# Regulatory Subfunctionalization drives OXPHOS evolution in teleosts

**DOI:** 10.1101/2024.07.30.605748

**Authors:** A Tsipourlianos, JCR Cardoso, R Angelakopoulos, A Kotoula, DM Power, Z Mamuris, KA Moutou

## Abstract

Oxidative phosphorylation (OXPHOS) supplies over 90% of the cellular ATP requirements through the orchestrated function of five multiprotein complexes located in the inner mitochondrial membrane. The study explores how the evolutionary constraints imposed by the indispensable function of OXPHOS guide the outcome of Whole Genome Duplication (WGD) events. Two teleosts, the gilthead seabream (*Sparus aurata*) and the European seabass (*Dicentrarchus labrax*), which have undergone three rounds of WGDs were used as models and OXPHOS gene expression was assessed during the highly plastic and energy-demanding period of early development. Fish ontogeny is a unique period in fish life marked with dramatic changes in morphology, physiology, metabolism and behavior. The tightly regulated landscape of cell divisions, migrations and differentiation driving these dramatic changes demands high energy supplies. The discovery of 24 and 22 OXPHOS gene families in gilthead seabream and European seabass genome, respectively, and the subsequent phylogenetic analysis showed in most cases divergence from a common ancestor at the base of the teleost lineage, a process attributed to teleost-specific WGD. Overall results indicate that the WGD events have resulted in early retention of OXPHOS paralogue genes and subsequent species- or lineage-specific losses. OXPHOS paralogue gene expression levels were compared following RNA sequencing within and between distinct developmental stages in gilthead seabream and European seabass. Different expression patterns between paralogs were revealed; some were in dosage balance, others were expressed only in particular stage(s) and a lot of them were differentially expressed between stages. Differences in both the number and location of SNPs were revealed between paralogs, after merging the RNA sequencing data with whole genome sequencing data. A considerable number of mutations were mapped in the UTRs and mostly synonymous substitutions were identified in the CDS. The ratio of non-synonymous to synonymous substitutions when comparing the CDS variants revealed neutral positions while others are subject to purifying selection, safeguarding protein structural integrity and/or function. Overall, regulatory subfunctionalization of OXPHOS paralogs appeared as the evolutionary mechanism behind the retention of the paralogs in gilthead seabream and European seabass genomes in favor of ontogenetic plasticity.

## 1. Introduction

The publication of Susumu Ohno’s book “Evolution by Gene Duplication” in 1970 established gene and whole genome duplication (WGD) as key evolutionary forces; they create new genetic material that may be altered by mutations and natural selection, leading to the emergence of new characteristics adding to species complexity. Whole genome duplication, segmental duplication, tandem duplication, and transposon-induced duplication are some of the mechanisms that produce duplicate genes (Jiang et al., 2004; Drouin & Dover, 1990; Bailey et al., 2002; Zhang, 2003; Panchy et al., 2016). Each of these mechanisms may have a different effect on the function, evolutionary fate, and genomic architecture of duplicate genes. Following gene duplication, newly acquired paralogous sequences are presumed to operate similarly to the original gene, although functional diversity may occur via neofunctionalization (Ohno 1970) or subfunctionalization (Force et al., 1999) that can contribute to the preservation of both paralogous genes (Drobek, 2022). In both circumstances, mutations may arise in the coding or regulatory sequence at different positions of each duplicate, leading to changes in the coding sequence and the protein structure, or the spatiotemporal regulation of the gene (Duarte et al., 2006; Kocha et al., 2015; Maouche et al., 2018; Martins et al., 2013; Shew et al., 2021; Sun et al., 2010). In contrast, the accumulation of harmful mutations can lead to non-functionalization eliminating the redundant copies. Within the same organism, various tissues and organs may exhibit varying degrees of functional redundancy and divergence across duplicates (Bermejo-Nogales et al., 2015; Gallagher & Macqueen, 2016; Mankiewicz & Cleveland, 2021; Porplycia et al., 2017). In addition, certain duplicated genes have the capacity to carry out redundant tasks throughout development and to switch to different roles in adulthood (Bochkis et al., 2012).

Oxidative phosphorylation (OXPHOS) is a fundamental biological process providing almost 90% of cellular energy in the form of ATP. OXPHOS is located in the inner membrane of mitochondria and involves five multiprotein complexes. Four of them (complexes I-IV) build the respiratory chain, responsible for creating the proton gradient across the inner mitochondrial membrane, while complex V is the ATP synthase that uses the proton gradient to produce ATP. The genes for the subunits of four of the five complexes are located in both nuclear and mitochondrial DNA (Wolff et al., 2014). As a result, the cross-talking of the two genomes (mitonuclear interactions), protein synthesis and targeting, and the assembly of OXPHOS complexes and super complexes (Signes & Fernandez-Vizarra, 2018) require complex regulatory mechanisms.

Given the complexity of OXPHOS, it is possible that the unique evolutionary opportunities provided by the simultaneous duplication by WGD, had a considerable impact on its evolution. Two WGD events (1R and 2R) took place early in the evolution of vertebrates, while a third WGD event (3R) or teleost-specific genome duplication (TSGD) took place before the divergence of the teleosts (Meyer & Van De Peer, 2005; Sémon & Wolfe, 2007; Van De Peer et al., 2009). TSGD is considered pivotal to the fast divergence of the teleosts, in part because it allowed for the acquisition of novel gene functions (Glasauer & Neuhauss, 2014; McClintock et al., 2001; Volff, 2005; Woods et al., 2005). The taxonomic group of teleosts contains more than 26K species exhibiting an extraordinary range of adaptations that allow them to inhabit highly varied environments and make them excellent model organisms for the study of evolutionary mechanisms. The evolution of OXPHOS nuclear genes has been studied in fish and other vertebrates, providing insights into the impact of WGDs on the OXPHOS machinery (De Grassi et al., 2005; De Grassi et al., 2008; Little et al., 2010) and their possible evolutionary fates (Porplycia et al., 2017). While the impact of gene and genome duplication on OXPHOS genes is well acknowledged, the existing literature tends to focus on specific genes, somewhat constraining the comprehensive understanding of evolutionary dynamics within the entire OXPHOS gene repertoire. This study investigates the fate of nuclear genes encoding OXPHOS proteins post whole-genome duplication (WGD). With focus on gilthead seabream (*Sparus aurata*) and the European seabass (*Dicentrarchus labrax*), two euryhaline and omnivorous species of high economic importance for Mediterranean marine fish farming. Both species are emblematic for Mediterranean aquaculture possessing the 3rd and 4th position, respectively, in European Union aquaculture total production (Eurostat 2021). Employing multiple methodologies, including gene mapping, subcellular localization inference, phylogenetic analysis, and examination of gene polymorphism, it aims to understand the evolutionary trajectories and regulatory mechanisms of these duplicated genes. Additionally, integrating transcriptome-level analyses during developmental stages provides insights into the functional implications of WGD on OXPHOS gene evolution and their roles in cellular functions across different developmental contexts.

## 2. Materials and methods

### 2.1. Database mining and sequence retrieval

In order to study gene evolution and potential divergent function of OXPHOS in gilthead seabream (*Sparus aurata*) and European seabass (*Dicentrarchus labrax*), the genome of the two fish species from Ensembl (database Release 105) (Cunningham et al., 2022) was searched for orthologs of Human OXPHOS genes based on HUGO Gene Nomenclature Committee (HGNC), accessed in March 2022, using the human (*Homo sapiens*) sequences as queries and the identity of the sequences retrieved were further confirmed by a reciprocal BLAST approach. In order to find potential OXPHOS gene families, the genomes of the two species were searched for paralogs using a reciprocal BLAST approach, using the previously found OXPHOS genes (best hit) of each species as query sequence. Only the OXPHOS-related paralogs were taken into account for the rest of the analysis in cases where paralogs were found. For comparative analysis searches for homologues in other species genomes were also performed for the genes encoding for the five complexes of OXPHOS. The species that were included in the analysis were several teleost representatives, including Nile tilapia (*Oreochromis niloticus*), medaka (*Oryzias latipes*), amazon molly (*Poecilia Formosa*), platyfish (*Xiphophorus maculatus*), stickleback (*Gasterosteus aculeatus*), greater amberjack (*Seriola dumerili*), fugu (*Takifugu rubripes*) and tetraodon (*Tetraodon nigroviridis*). Searches also included the genomes of the jawless fish sea lamprey (*Petromyzon marinus*) and the cartilaginous fish elephant shark (*Callorhinchus milii*), as representatives of basal vertebrate lineages; the early diverging ray-finned fish, spotted gar (*Lepisosteus oculatus*); the basal Sarcopterygian representative, the African coelacanth (*Latimeria chalumnae*); two tetrapods, mouse (*Mus musculus*) and chicken (*Gallus gallus*); and finally amphioxus (*Branchiostoma floridae*) as a representative of early chordata. Using the BLAST algorithm (Altschul et al., 1990), homolog genes were identified in the Ensembl and/or NCBI (Sayers et al., 2022) genomic databases (Ensembl Release 105) based on sequence similarity and e-value (<1 x 10^-5^) and, once again, only the OXPHOS related paralogs were taken into account. Their identity was then confirmed by reciprocal BLAST against the human, gilthead seabream and European seabass genomes. At this point we need to acknowledge that the pipeline used to identify orthologs and putative paralogs is prone to potential errors due to assembly quality and annotation methods.

### 2.2. Sequence alignments and phylogenetic analysis

Although peptide sequences are preferred (Foster, & Hickey, 1999) for phylogenetic research, the deduced protein sequences of the examined genes often yielded short sequences, which had a negative impact on phylogenetic signal. For the majority of the reconstructed phylogenies, nucleotide coding sequences (CDS) were used. The MAFT (Katoh, 2002) algorithm was used to perform multiple sequence alignment. Alignments where visualized using AliView (Larsson, A. (2014) and gaps were removed. After manually editing for removing the gaps in the alignment and removing poorly aligned sequences, the edited alignment was used as input to MEGAX (Kumar et al., 2018) in order to find the optimal nucleotide/amino acid substitution model (Table S1) that would describe the evolution of gene families most effectively. Lastly, Maximum Likelihood algorithm was used in the phylogenetic analysis. MEGAX was used to reconstruct the phylogenetic trees, and 100 bootstrap repetitions were performed to statistically analyze the sequence branches that were constructed. The generated topologies of the phylogenetic trees for each gene family were evaluated in respect to the three rounds of vertebrate genome duplications, more over we have inferred the evolution of the genes of interest based on the tree topology and the bootstrap support values of the branches considering the evolution of the species included.

### 2.3. Collinearity analysis and duplicate gene origin classification

The Multiple Collinearity Scan Toolkit (MCScanX) (Wang et al., 2012) was used to examine the degree of collinearity in the genomes of the gilthead seabream and European seabass. Blastp searches were run using the command-line program BLAST+ (Camacho et al., 2009) with an e-value threshold set to <1 x 10^-5^ in order to find paralogous regions in both fish genomes. Ensembl annotation files for both species were parsed to produce the gff files required by MCScanX, with the MATCH SIZE option set to 2, which indicates that two genes are necessary to create a collinear block. The same files were used to investigate the genes evolutionary origins, in the MCScanX tool called duplicate gene classifier, which classified each duplicated gene in the gilthead seabream and European seabass according to their origin. Genes were categorized into whole-genome or segmental duplicates, which include anchor genes in collinear blocks, tandem duplicates, which are paralogs that are physically next to one another in the genome, proximal duplicates, which have multiple genes between them but are mapped in a nearby chromosomal region, and dispersed duplicates, which do not fit into any of the above-mentioned categories. While running the duplicate gene classifier the MATCH SIZE and N PROXIMAL parameters, which specify the maximum number of genes between two proximal duplicates, were set to 2 and 10, respectively.

### 2.4. Prediction of subcellular localization

The prediction tools DeepLoc 2.0 (Thumuluri et al., 2022) and TargetP 2.0 (Almagro Armenteros et al., 2019) were used to determine the subcellular location of the gilthead seabream and seabass proteins generated by the paralog genes. The mitochondrial localization was confirmed using DeepLoc, and the presence of the mitochondrial transit peptide and the location of the cleavage site were predicted using TargetP.

### 2.5. Functional conservation estimate

The InterPro 91.0 database (Paysan-Lafosse et al., 2023), accessed in November 2022, was utilized to investigate the degree of conserved function between the human, gilthead seabream and European seabass homologues. The protein sequences were entered into InterPro, and the outcomes of the fish paralogs and their human homologs were compared. On the basis of conserved domains or sites related to function as identified on their peptide sequences, the human and fish proteins were categorized into families and GO terms related to their biological processes and molecular functions. Lastly, CDS, UTRs, introns, and Pfam (Mistry et al., 2021) domains were obtained from Ensembl and plotted using GSDS 2.0 (Hu et al., 2015) for comparisons in order to estimate the degree of gene structure and protein domain conservation.

### 2.6. Larvae rearing and sampling

The pseudo-green approach was used for rearing gilthead seabream and European seabass, in accordance with the accepted methodology for intense larval rearing (Papandroulakis et al., 2001). Larval rearing was performed at the Institute of Marine Biology, Biotechnology and Aquaculture (HCMR). Recirculation systems with 500 L, black, cylindro-conical tanks (one tank per duplicate) and biological filters were utilized. Chillers and heaters were used to change the water’s temperature to the required levels for the tests, and they were automatically controlled by electronic sensors (Eliwell, EU). From the egg stage (epiboly) until mid metamorphosis, the temperature was adjusted at 20°C and 17°C for gilthead seabream and European seabass, respectively. Larvae were successively fed on enriched (DHA Protein Selco, INVE) rotifers until they reached 6 mm total length (TL), enriched rotifers and *Artemia* nauplii (Easy DHA Selco, INVE) when they were between 6- and 10-mm TL and enriched *Artemia* nauplii after 10 mm TL. The rotifer concentration in the tanks was adjusted twice daily to 6–8 ind ml^−1^. The concentration of *Artemia* nauplii was adjusted twice daily to 0.5–2 ind ml^−1^. During the rotifer-feeding period, live microalgae *Chlorella* sp. were added thrice daily at a concentration of 6.5 ± 3. 10^5^ cells ml^−1^. Weaning to inert diets (O-range, INVE) started when fish were approximately 8 mm TL. During the rotifer-feeding period, the excess rotifers were removed from the culture medium by means of planktonic traps (50 μm mesh size), positioned at the water-outlet of tanks. During the first twenty days of feeding, air-blowing skimmers were installed to keep the water surface free from lipids and enhance swim bladder inflation. Water oxygen saturation was 5.8 ± 1.0 to 6.0 ± 0.9 mg/L^-1^, pH 7.8 ± 0.3, salinity 35–36 ‰ and photoperiod 18L:6D. The water turnover rate ranged from 20% of the tank volume d^-1^ during the autotrophic phase to 70% h^-1^ at the end of larval rearing. Water was pumped from a deep (> 100 m) borehole.

The same protocol for larvae sampling was used for both species. A random sample of larvae from each tank was collected, larvae were anaesthetized using 2-Phenoxyethanol (Sigma), rinsing with clean sea water and then placing in fixative, RNA later® solution; stored at 4°C overnight (to allow the solution to thoroughly penetrate the larvae), then at –20°C for longer-term storage. From both species samples were collected at three developmental stages, namely first feeding (FF), notochord flexion (FL), and mid metamorphosis (MM).

### 2.7. RNA extraction and transcriptome sequencing

Using the E.Z.N.A.® Total RNA Kit I, RNA was isolated from the total body of individuals at each developmental stage in both species. In order to have an adequate total RNA yield 3-5 individuals were pooled to generate a sample at FF, whereas individual RNA extractions were performed at FL and MM. Subsequently, twelve RNA samples were pooled for each developmental stage, and the pooled samples were sent to Novogene inc. (Europe) for evaluation. The RNA fragments were utilized to create cDNA, construct libraries and sequence using the HiSeq 2000 platform (Illumina, USA) with 150bp paired-end reads, producing about 30M reads per sample with over 97% of the reads passing Q20.

Fastp (Chen et al., 2018) was used to initially process the raw data (raw reads) in FASTQ format. In this stage, reads with low quality and reads including adaptor and poly-N sequences were removed from the raw data to provide clean data (clean reads). The reference genome and gene annotation files were directly obtained from the Ensembl genome browser website. Using the HISAT2 program (Kim et al., 2019), paired-end clean reads were mapped to the reference genomes (fSpaAur1.1, INSDC Assembly GCA 900880675.1, July 2019 and seabass_V1.0, INSDC Assembly GCA_000689215.1, Apr 2014). Based on recommendations for datasets lacking biological replicates, edgeR (Robinson et al., 2010) was used to analyze the differential expression of two conditions. By utilizing the Benjamini and Hochberg approaches, the P values were modified. For significantly different expression, a corrected pvalue of 0.05 and a log2 (Fold Change) of 0.6 were established as the thresholds. Following that, data on the genes of interest were retrieved using R base commands (R Core Team, 2022; RStudio Team, 2022).

### 2.8. SNP identification and selection analysis

To identify SNPs to the transcript sequences of gilthead seabream and European seabass OXPHOS paralogs, the transcriptome data for both species were used in addition to whole genome resequencing data from four individuals of gilthead seabream, one from France and three from Greece, and from five individuals of European seabass from Greece.

According to the manufacturer’s instructions, PureLink Genomic DNA Mini Kit (Invitrogen, Catalogue number: K182002) was used for genomic DNA extraction. Agarose electrophoresis was used to determine the quality of the DNA, and the Qubit dsDNA BR Assay Kit was used to determine its amount (Invitrogen, Catalogue number: Q32850). The DNA samples were delivered to Novogene, where they underwent Illumina HiSeq 3000 sequencing at a mean sequencing coverage of 100x, assuming a uniform 25x coverage for each sample. FastQC (Andrews, 2010) was used to evaluate the quality of FASTQ files, and any sequences with PHRED scores lower than 30 were discarded. From the Ensembl Database, the reference genomes (fSpaAur1.1, INSDC Assembly GCA 900880675.1, seabass_V1.0, INSDC Assembly GCA_000689215.1) were obtained and indexed with bwa. Using bwa-mem (Li, 2013), pairs of sequences that satisfied the quality standards were mapped in the indexed genome. The SAM files of the alignments were converted to BAM files, sorted and indexed for the subsequent analysis using SAMtools (Li et al., 2009). Individual BAM files were merged using SAMtools. The merged BAM file was used for the variant calling using bcftools (Li, 2011). The quality filtering of SNPs was performed using bcftools with the following arguments, QUAL>20 and DP>10. The variant annotation and effect prediction was carried out using the tool SnpEff (Cingolani et al., 2012), with the gtf annotation file provided by Ensembl as input. FEL (Kosakovsky Pond et al., 2006) algorithm as included in Datamonkey (Delport et al., 2010) was sued to estimate the rates of synonymous and non-synonymous substitutions and to identify polymorphic sites under positive or negative selection.

## 3. Results

### 3.1. OXPHOS gene families in gilthead seabream and European seabass

Several OXPHOS gene families were discovered in both species after thorough searches using reciprocal BLAST to find OXPHOS orthologs between the human and fish genomes and within European seabass and gilthead seabream genome respectively, to find putative paralogs. Gene families were discovered in all five OXPHOS complexes, increasing the total number of OXPHOS genes to 105 and 103 for gilthead seabream and European seabass, respectively, in comparison with the 85 OXPHOS genes in the human genome (Figure 1). Spatial distribution and organization of OXPHOS genes across the genomes of the gilthead seabream and the European seabass are shown in Figures 2–3. Gene names were not consistently assigned to fish paralogs and/or human homologs based on Ensembl annotation, and some of them were designated as novel genes. The identified genes were renamed based on the gene name of their homolog at the human genome (best BLAST hit), adding at random an ascending number for each paralog to be more systematic and make it simpler to follow the gene families throughout the manuscript. The correspondence between the used gene name and Ensembl gene IDs is shown in Table S2.

**Figure 1.**
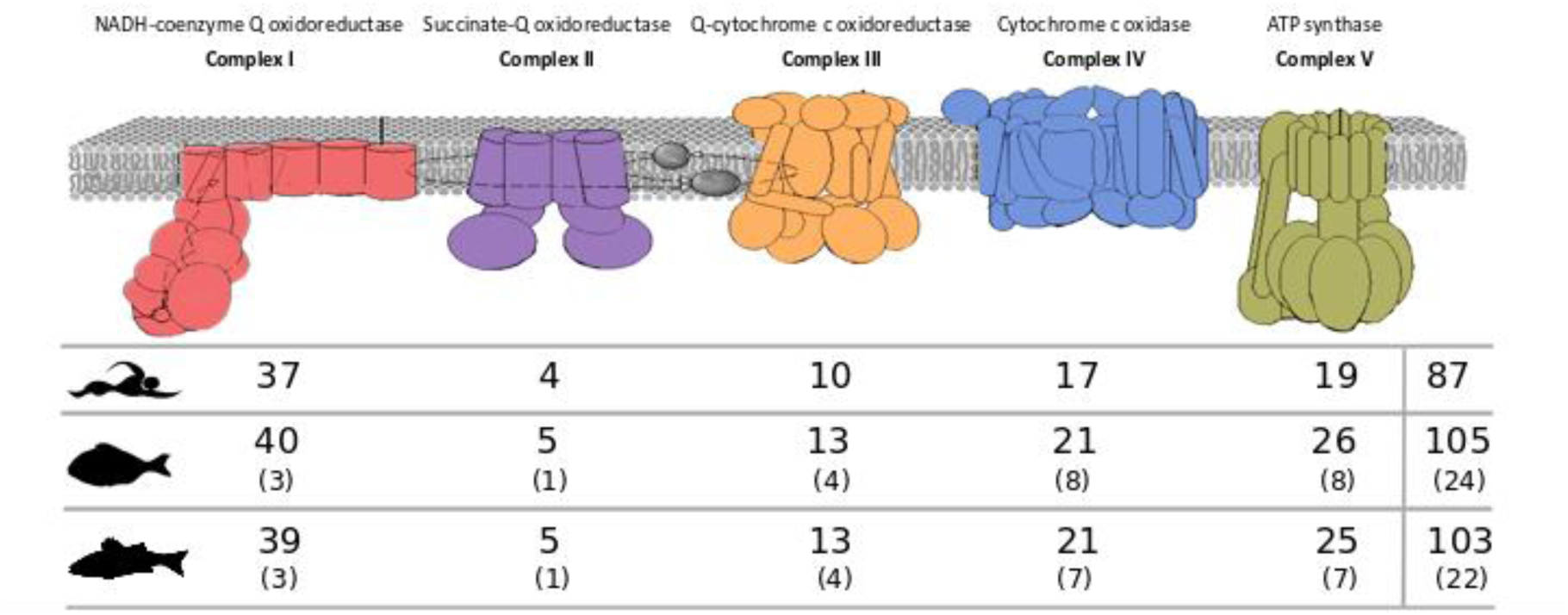
Number of OXPHOS genes per complex in the human, gilthead seabream and European seabass. In parenthesis, the number of gene families identified in the genomes of gilthead seabream and European seabass, respectively, are given.

**Figure 2.**
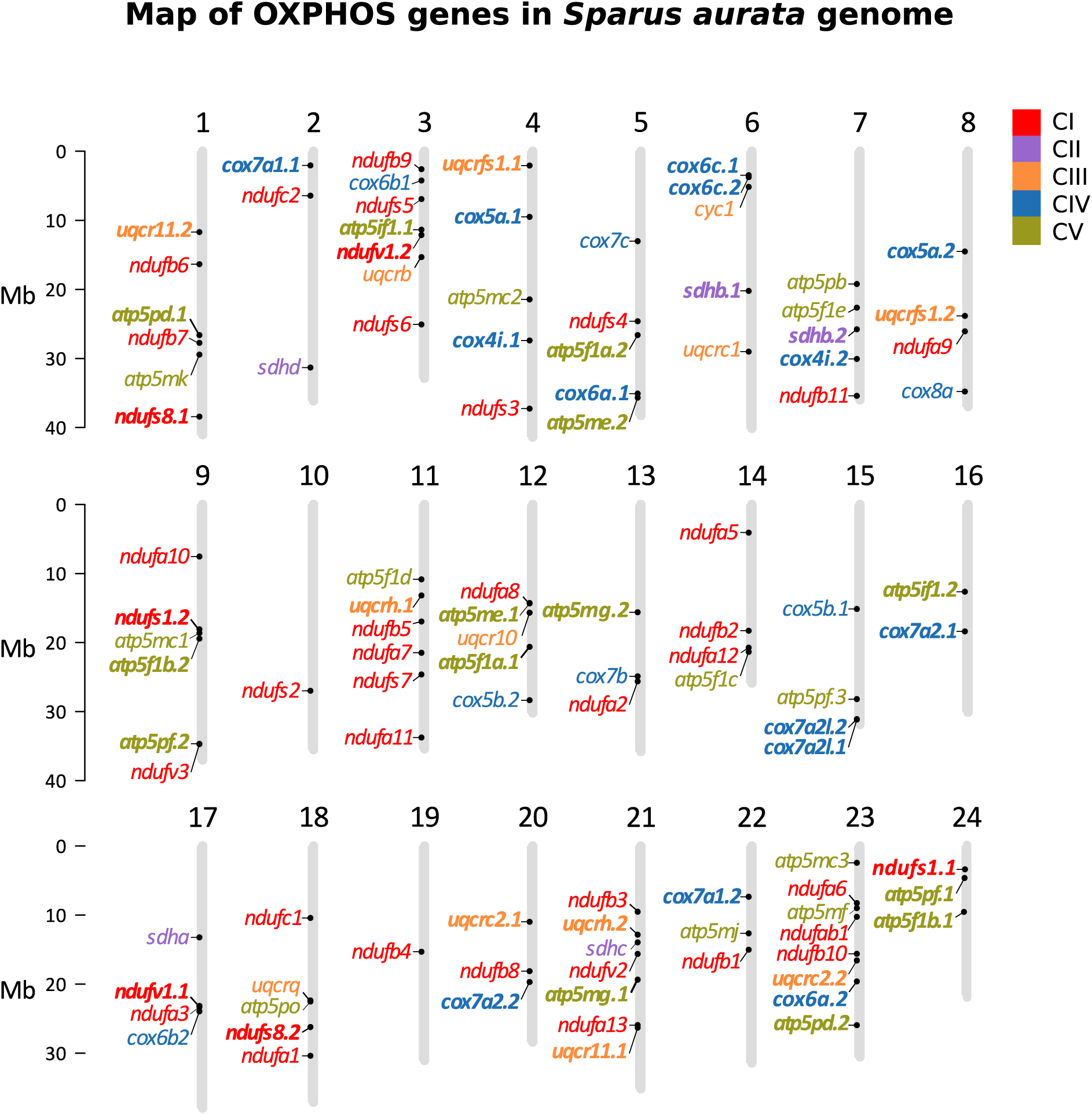
Mapping of OXPHOS genes onto the gilthead seabream genome (fSpaAur1.1) at the chromosome level was conducted using the R package chromPlot (Oróstica & Verdugo, 2016). Different colors were assigned to indicate the five OXPHOS complexes, and genes belonging to OXPHOS gene families are highlighted in bold.

**Figure 3.**
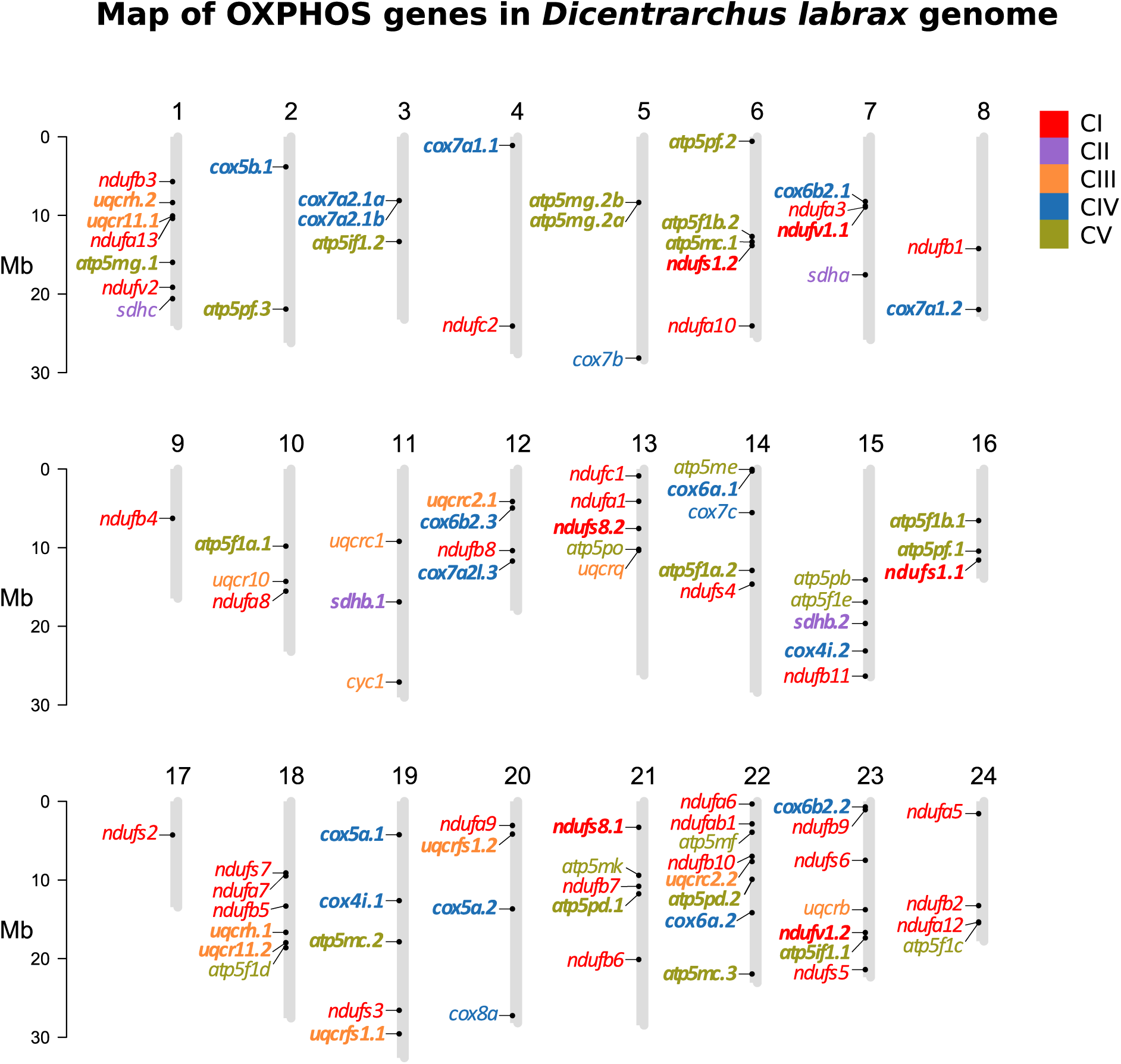
Mapping of OXPHOS genes onto the European seabass genome (seabass_V1.0) at the scaffold level was conducted using the R package chromPlot (Oróstica & Verdugo, 2016). Different colors were assigned to indicate the five OXPHOS complexes, and genes belonging to OXPHOS gene families are highlighted in bold.

While the findings between the two species were relatively similar, some differences were observed. Twenty-four (24) OXPHOS gene families were identified in gilthead seabream and 22 in European seabass with *atp5me* and *cox6c* being the extra gilthead seabream families. Most shared gene families between the two species had the same gene number, except for the *atp5mg*, *cox6b2*, *cox7a1* and *cox7a2l* gene families (Table 1).

**Table 1.** Predictions in silico of subcellular localization of the products of OXPHOS paralogs in gilthead seabream and European seabass. Most gene products were predicted to be targeted in the mitochondrion. The exceptions are indicated in red.

| *Sparus aurata* |  |  | *Dicentrarchus labrax* |  |  |
| --- | --- | --- | --- | --- | --- |
| Gene name | Localization | Probability | Gene name | Localization | Probability |
| **Complex I** |  |  |  |  |  |
| *ndufs1.1* | Cytoplasm | 0.7156 | *ndufs1.1* | Mitochondrion | 0.9113 |
| *ndufs1.2* | Mitochondrion | 0.9175 | *ndufs1.2* | Mitochondrion | 0.8681 |
| *ndufs8.1* | Mitochondrion | 0.9443 | *ndufs8.1* | Mitochondrion | 0.9514 |
| *ndufs8.2* | Mitochondrion | 0.9431 | *ndufs8.2* | Mitochondrion | 0.9398 |
| *ndufv1.1* | Mitochondrion | 0.8658 | *ndufv1.1* | Mitochondrion | 0.8198 |
| *ndufv1.2* | Mitochondrion | 0.9270 | *ndufv1.2* | Mitochondrion | 0.9207 |
| **Complex II** |  |  |  |  |  |
| *sdhb.1* | Mitochondrion | 0.8927 | *sdhb.1* | Mitochondrion | 0.8516 |
| *sdhb.2* | Mitochondrion | 0.8736 | *sdhb.2* | Mitochondrion | 0.8858 |
| **Complex III** |  |  |  |  |  |
| *uqcr11.1* | Mitochondrion | 0.9766 | *uqcr11.1* | Mitochondrion | 0.9752 |
| *uqcr11.2* | Mitochondrion | 0.9645 | *uqcr11.2* | Mitochondrion | 0.9690 |
| *uqcrc2.1* | Mitochondrion | 0.9682 | *uqcrc2.1* | Mitochondrion | 0.9669 |
| *uqcrc2.2* | Mitochondrion | 0.9359 | *uqcrc2.2* | Mitochondrion | 0.9302 |
| *uqcrfs1.1* | Mitochondrion | 0.8842 | *uqcrfs1.1* | Mitochondrion | 0.9105 |
| *uqcrfs1.2* | Cytoplasm, Mitochondrion | 0.6149, 0.6460 | *uqcrfs1.2* | Mitochondrion | 0.9415 |
| *uqcrh.1* | Cytoplasm, Mitochondrion | 0.5312, 0.6471 | *uqcrh.1* | Cytoplasm, Mitochondrion | 0.5497, 0.6586 |
| *uqcrh.2* | Cytoplasm | 0.5133 | *uqcrh.2* | Cytoplasm | 0.5530 |
| **Complex IV** |  |  |  |  |  |
| *cox4i.1* | Mitochondrion | 0.9329 | *cox4i.1* | Mitochondrion | 0.9374 |
| *cox4i.2* | Mitochondrion | 0.9513 | *cox4i.2* | Mitochondrion | 0.9257 |
| *cox5a.1* | Mitochondrion | 0.9204 | *cox5a.1* | Mitochondrion | 0.9286 |
| *cox5a.2* | Mitochondrion | 0.9314 | *cox5a.2* | Mitochondrion | 0.9572 |
| *cox5b.1* | Mitochondrion | 0.9448 | *cox5b.1* | Mitochondrion | 0.9208 |
| *cox5b.2* | Mitochondrion | 0.9555 | *cox5b.2* | Mitochondrion | 0.9339 |
| *cox6a.1* | Mitochondrion | 0.9032 | *cox6a.1* | Mitochondrion | 0.892 |
| *cox6a.2* | Mitochondrion | 0.9095 | *cox6a.2* | Mitochondrion | 0.9241 |
| *cox6b2.1* | Cytoplasm, Mitochondrion | 0.5627, 0.6472 | *cox6b2.1* | Cytoplasm | 0.5835 |
| *cox6b2.2* | Cytoplasm | 0.5038 | *cox6b2.2* | Cytoplasm | 0.5590 |
|  |  |  | *cox6b2.3* | Cytoplasm | 0.5590 |
| *cox6c.1* | Mitochondrion | 0.9246 |  |  |  |
| *cox6c.2* | Mitochondrion | 0.9350 |  |  |  |
| *cox7a1.1* | Mitochondrion | 0.9392 | *cox7a1.1* | Mitochondrion | 0.9376 |
| *cox7a1.2* | Mitochondrion | 0.9363 | *cox7a1.2* | Mitochondrion | 0.9401 |
| *cox7a2.1* | Mitochondrion | 0.9337 | *cox7a2.1a* | Mitochondrion | 0.9286 |
|  |  |  | *cox7a2.1b* | Mitochondrion | 0.9008 |
| *cox7a2l.1* | Mitochondrion | 0.9299 | *cox7a2l.1* | Mitochondrion | 0.9190 |
| *cox7a2l.2* | Mitochondrion | 0.9299 |  |  |  |
| *cox7a2l.3* | Mitochondrion | 0.9223 | *cox7a2l.3* | Mitochondrion | 0.9114 |
| **Complex V** |  |  |  |  |  |
| *atp5f1a.1* | Mitochondrion | 0.8939 | *atp5f1a.1* | Mitochondrion | 0.8726 |
| *atp5f1a.2* | Mitochondrion | 0.6299 | *atp5f1a.2* | Mitochondrion | 0.8594 |
| *atp5f1b.1* | Mitochondrion | 0.8256 | *atp5f1b.1* | Mitochondrion | 0.7482 |
| *atp5f1b.2* | Mitochondrion | 0.8942 | *atp5f1b.2* | Mitochondrion | 0.8984 |
| *atp5if1.1* | Mitochondrion | 0.9714 | *atp5if1.1* | Mitochondrion | 0.9720 |
| *atp5if1.2* | Mitochondrion | 0.9747 | *atp5if1.2* | Mitochondrion | 0.9737 |
| *atp5mc.1* | Mitochondrion | 0.8216 | *atp5mc.1* | Mitochondrion | 0.8281 |
| *atp5mc.2* | Mitochondrion | 0.8046 | *atp5mc.2* | Mitochondrion | 0.7364 |
| *atp5mc.3* | Mitochondrion | 0.7420 | *atp5mc.3* | Mitochondrion | 0.7385 |
| *atp5me.1* | Mitochondrion | 0.8718 |  |  |  |
| *atp5me.2* | Mitochondrion | 0.8841 |  |  |  |
| *atp5mg.1* | Mitochondrion | 0.9382 | *atp5mg.1* | Mitochondrion | 0.9398 |
| *atp5mg.2* | Mitochondrion | 0.9532 | *atp5mg.2a* | Mitochondrion | 0.9354 |
|  |  |  | *atp5mg.2b* | Cytoplasm, Nucleus | 0.5851, 0.5860 |
| *atp5pd.1* | Mitochondrion | 0.8477 | *atp5pd.1* | Mitochondrion | 0.9097 |
| *atp5pd.2* | Mitochondrion | 0.9526 | *atp5pd.2* | Mitochondrion | 0.9349 |
| *atp5pf.1* | Mitochondrion | 0.9568 | *atp5pf.1* | Mitochondrion | 0.9415 |
| *atp5pf.2* | Mitochondrion | 0.9124 | *atp5pf.2* | Mitochondrion | 0.9010 |
| *atp5pf.3* | Cytoplasm | 0.6992 | *atp5pf.3* | Cytoplasm, Mitochondrion | 0.5936, 0.6593 |

### 3.2. Evolutionary origin of paralogs

Different mechanisms, including small-scale duplications like single-gene or segmental duplications, and WGD, can lead to the emergence of duplicated genes. In contrast to humans and other non-teleost vertebrates, which have undergone only the first two rounds of vertebrate genome duplication, teleost evolution was accompanied by a third round of WGD, the TSGD. Phylogenetic and synteny analyses were performed to determine the evolutionary origin of gilthead seabream and European seabass paralogs. Three main topologies for gene paralogs were identified and supported by bootstrap confidence values (Figure 4). Phylogenetic trees for each gene family can be found in Figures S1-S22.

**Figure 4.**
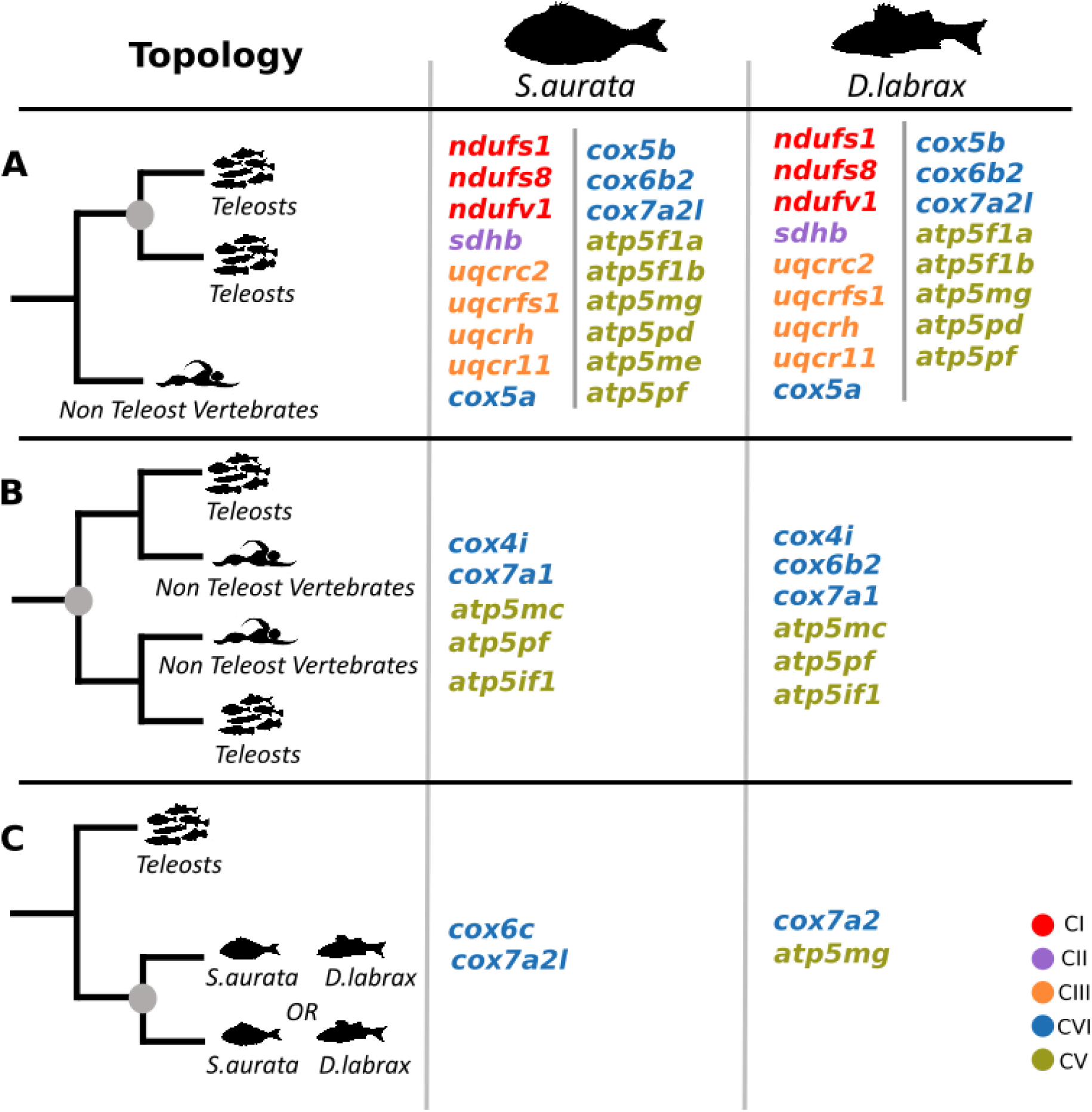
Observed topologies from the phylogenetic analysis of the OXPHOS gene families in gilthead seabream and European seabass. Grey cycles on the topology trees represent duplication nodes. The genes of each OXPHOS complex share the same colour.

Topology A demonstrates that both paralogs descended from a duplication that occurred in the common ancestor of teleosts and non-teleost vertebrates, most likely the first or second round of vertebrate WGDs. Topology B indicates a WGD event, most likely the TSGD, as both paralogs descended from a duplication that occurred at the common ancestor of teleosts. Topology C shows that the grouping of paralogs is species-specific, which suggests either a within-species or a lineage-specific duplication that took place at the outer branches of teleosts and it is not represented by any other species in the analysis. The *cox7a* gene family (Figure S14) was the only exception to the above, as the observed topology could not be fully explained by the vertebrate WGD events alone, indicating that there was at least one ancestral duplication that resulted in the *cox7a* and *cox7a2l* gene families.

Eighteen (18) and 17 paralog pairs in gilthead seabream and European seabass, respectively, followed topology B, which matches the expected topology of TSGD (Figure 4). The observed topology in the case of the gene families *atp5f1b* (Figure S16) and *cox7a2l* (Figure S14) may suggest potential subsequently gene loss. However, we cannot discard that gene absence may also be the result of errors in genome assemblies or gene annotation mistakes. Six gene families have members that are following topology A (Figure 4), which is in accordance with one of the first two rounds of vertebrate genome duplications. From those, five families were shared between gilthead seabream and European seabass (*atp5if1, atp5mc, atp5pf, cox4i, cox7a1*) and one (*cox6b2*) was European seabass specific. Topology C was observed in two different gene families in each species. Species- or lineage-specific duplication events gave rise to members of *cox6c* and *cox7a2l* gene families in gilthead seabream and to *atp5mg* and *cox7a1* in European seabass. Lastly, the phylogenetic tree for *cox6a* gene family was not resolved.

Synteny analysis (Figures 5-6) classified OXPHOS paralogs into four categories: i) WGD/segmental, when paralogs are members of collinear blocks; ii) proximal, indicating that the paralogs are near each other but interrupted by several other genes; iii) tandem that indicates that paralogs are next to each other; and iv) dispersed, indicating that, while the genes are paralogs, they are not near each other on chromosomes nor show conserved synteny (Table S3). In gilthead seabream, the relationships between the members of 7 gene families were classified as WGD/segmental, 17 as dispersed and 2 as proximal, while in European seabass 7 were classified as WGD/segmental, 17 as dispersed, 1 as proximal and 1 as tandem. In both species, the families classified as proximal and/or tandem are the same that, based on phylogenetic analysis, were characterized as the outcome of species/ lineage specific duplications (Topology C). From the paralog pairs that were inferred as the possible result of TSGD (Topology B), the members of 5 gene families (*sdhb, uqcrc2, uqcr11, cox6b2* and *atp5f1a*) in gilthead seabream and 6 families (*sdhb, uqcrc2, uqcr11, cox6b2, atp5f1a and atp5f1b*) in European seabass were identified as members of collinear blocks, suggesting a conserved chromosomal neighborhood for those genes.

**Figure 5.**
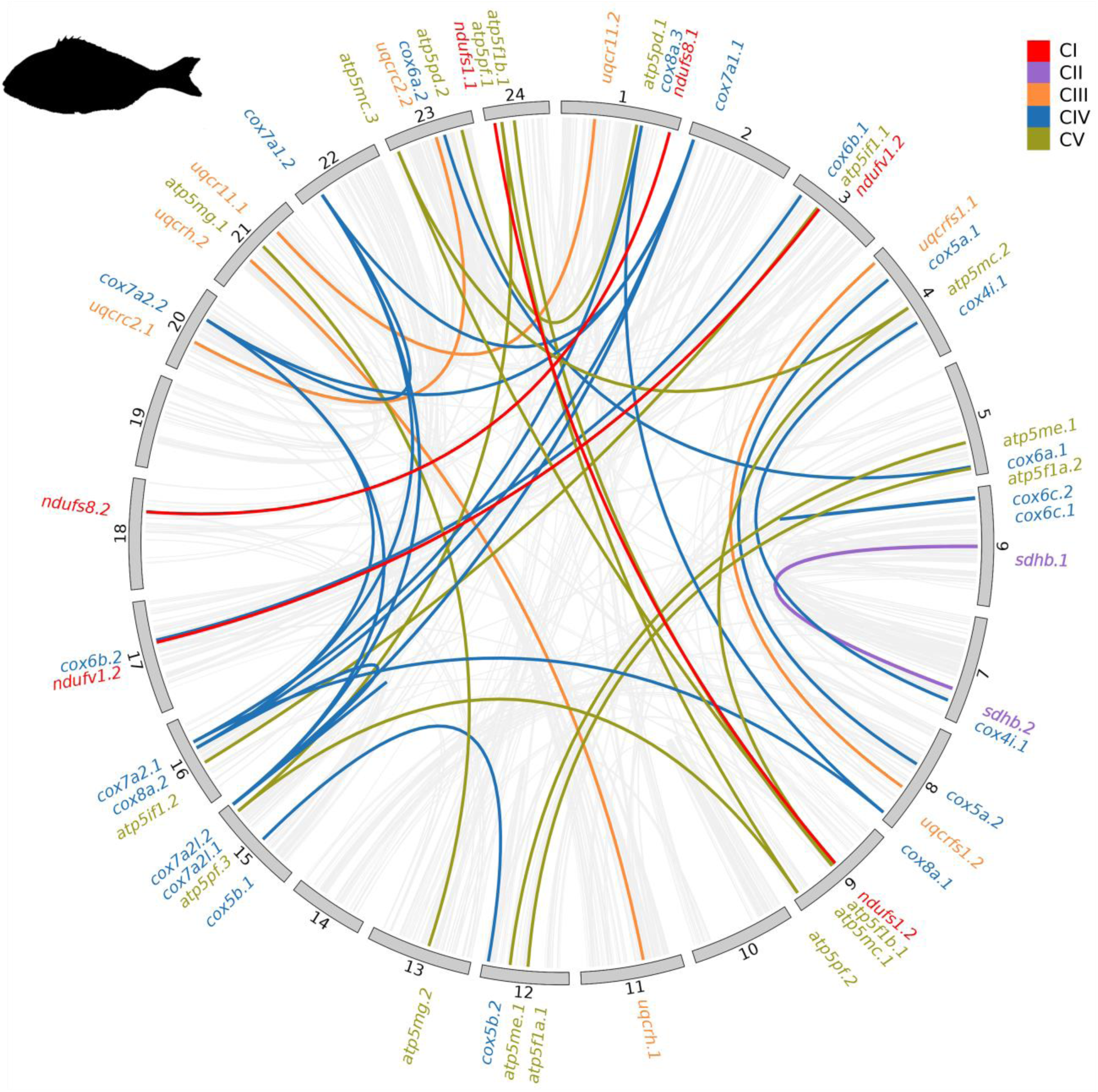
Circos plot depicting the paralogue relationships between gilthead seabream genes. Duplicated gene relationships between gilthead sea bream chromosomes are represented by inner lines. Different colors represent the five OXPHOS complexes.

**Figure 6.**
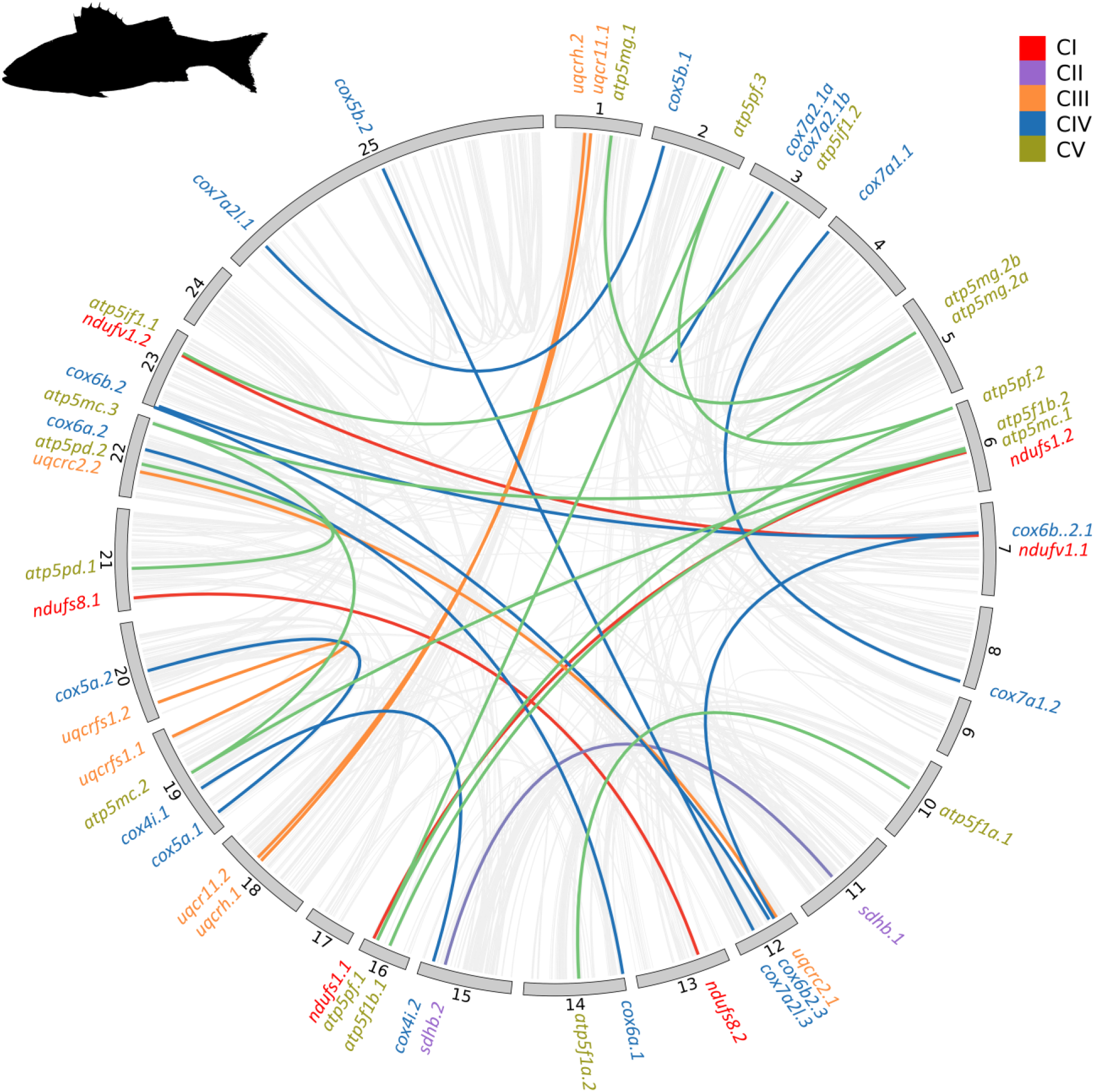
Circos plot depicting the paralogue relationships between European seabass genes. Duplicated gene relationships between European seabass chromosomes are represented by inner lines. Different colors represent the five OXPHOS complexes.

On the basis of phylogenetic tree topology, members of the remaining gene families for both species that were inferred to be TSGD paralogs, were not identified in collinear blocks and were categorized as dispersed duplicates. Given that they were members of two district teleost clades with a well-supported duplication node at their bases, it was hypothesized that these paralogs are the result of TSGD followed by 1) significant gene losses to the paralogs chromosomal environment, 2) chromosomal rearrangements close to the paralog locations, or 3) translocations mediated by transposons or retrotransposons, at least for the one paralog. The majority of the members of the gene families resulting from one of the first rounds of vertebrate WGDs based on phylogenetic tree Topology A (Figure 4) were also characterized as dispersed, and their current location may be the result of the same mechanisms as described above, leading to low conservation of the chromosomal gene environment of those paralogs, too.

### 3.3. *In silico* prediction of functional characteristics of OXPHOS paralogs

The prediction *in silico* of the subcellular localization for OXPHOS paralogs in both species suggests mitochondria as the most probable localization for the majority of the genes (Table 1), with similar predictions for both species. In the cases the prediction indicated mitochondria and cytosol, the probability for cytosol localization was lower than for mitochondria. In both species all members of gene families *cox6b2* and *uqcrh* were predicted to be localized in cytosol, yet with a relatively low probability for that prediction. While this could be an indication of diverged evolution for those gene families it seems that this is not the case. Upon further investigation, when the same analysis was followed for the human orthologs of those genes, similar results were yielded as with the fish genes; predictions for both mitochondria and cytosol with a low probability score. Even in the cases without a clear prediction for mitochondria localization, that prediction had always a higher probability compared with cytosol. Based on that and in respect to the level of genome and gene annotation of the human genome it is most probable that those genes are also localized in the mitochondria. In addition, based on InterPro’s predictions, functional analysis of OXPHOS gene families revealed that the majority of gilthead seabream and European seabass OXPHOS paralogs share protein family membership, molecular function, and biological processes with their human homolog for both species. The few exceptions to the above were the members of *uqcrc2* family in both species and *cox6c* family in gilthead seabream. Lastly the majority of OXPHOS paralogs had conserved gene structure (number of exons-introns; (Figures S23-S45). Certain exceptions were recorded and even in those cases a high conservation of protein domains was observed; *ndufs1.2* in gilthead seabream missing the NADH-quinone oxidoreductase domain that exists at the human ortholog and *ndufs1.1* gilthead seabream paralog; *uqcrfs1.2* missing two domains, UCR_TM and Ubiq-Cytc-red_N, in both species studied, while those domains exist in its human ortholog and in the *uqcrfs1.1* paralog of gilthead seabream and European seabass. In addition, there was no prediction for a protein domain in Pfam for European seabass genes *cox7a2.1b* and *cox7a2l.1*.

### 3.4. Genetic polymorphism and selection pressure

The total number of SNPs was identified along with the position (CDS, UTRs) in the sequence and the type of mutation (synonymous, missense) for all OXPHOS genes in both species (Table 2, Table S4-S5). Following the identification of the polymorphic sites at the coding sequences, selection analysis revealed the evolutionary mode in action for each position, again for all OXPHOS genes in both species (Table 2). No sequence mutations were found in seven (7) genes (*ndufv1.2, uqcrfs1.2, cox6a.2, cox7a2.1, atp5mc.2, atp5me.1 and atp5mg.2)* and nine (9) genes *(ndufv1.1, uqcr11.2, uqcrfs1.1, uqcrh.1, cox7a2.1b, cox7a2l.1, atp5f1b.1, atp5pd.1, atp5mc.2)* in gilthead seabream and European seabass, respectively. Three (3) gene families *(ndufv1, uqcrfs1 and cox7a2)* with no members bearing mutations were identified in common between both species. The majority of mutations in CDSs were synonymous, and those sites were either neutral or under purifying selection. Diversifying selection was observed only in four (4) genes of gilthead seabream (*ndufs1.1, atp5f1b.1, atp5pf.2, atp5pf.3*).

**Table 2.** The number of SNPs identified in the 52 genes of gilthead seabream and the 50 genes of European seabass, respectively, under different selection mode, and at different gene domains.

|  | *Gilthead seabream* | *European seabass* |
| --- | --- | --- |
| **Purifying** | 114 | 40 |
| **Diversifying** | 18 | 0 |
| **Neutral** | 129 | 73 |
| **Synonymous** | 182 | 96 |
| **Missense** | 99 | 30 |
| **5'UTR** | 42 | 15 |
| **3'UTR** | 148 | 60 |
| **Splice region** | 14 | 1 |

Different number of mutations were observed between paralogs for several gene families where; one paralog appeared to be highly conserved (few or even no mutations), whereas the other paralog showed higher number of mutation accumulation (e.g. *ndufv1, uqcr11, uqcrfs1, atp5pd, atp5me*); on the other hand, the paralogs of several gene families appeared to have similar number of mutations (e.g. *uqcrc2, cox4i, cox5a, cox6c, atp5f1a, atp5if1*); lastly, five (5) gene families (*uqcrc2, cox4i*, *atp5f1a*, *atp5if1*, *atp5mc*) exhibited similar pattern of mutation accumulation between the paralogs and that pattern did not differentiate between the two species (Figures S46 – S48).

### 3.5. OXPHOS gene expression in development

While we conducted a differential gene expression analysis, we acknowledge that the absence of biological replicates may compromise the reliability of the analysis. Thus, our findings are presented here solely for exploratory purposes.

Transcriptome analysis in gilthead seabream revealed that all OXPHOS genes were expressed at least at one of the three developmental stages examined, with the only exception the gene *cox7a2l.1,* which was not detected at any stage. Distinct ontogenetic profiles were drawn for several genes (Figure 7), as a number of genes identified as DEGs (log2 fold change above 0.6) in the comparisons between the developmental stages (Figure 7, Tables S6-S8). Most of the DEGs were downregulated during the transition from FF to FL, while the opposite was observed for the transition from FL to MM. Additionally, a number of genes were expressed only at one stage, with traces or no expression in the other developmental stage(s), indicative of a stage-specific regulation of expression. Interestingly, 36 of the 52 seabream OXPHOS paralogs appeared to be DEGs, at least according to one stage-to-stage comparison and they were among the five and eight genes for gilthead seabream and European seabass respectively, that exhibited stage-specific expression patterns. Within each developmental stage, differences between the expression levels of members of several gene families were observed, and those patterns were changing ontogenetically.

**Figure 7.**
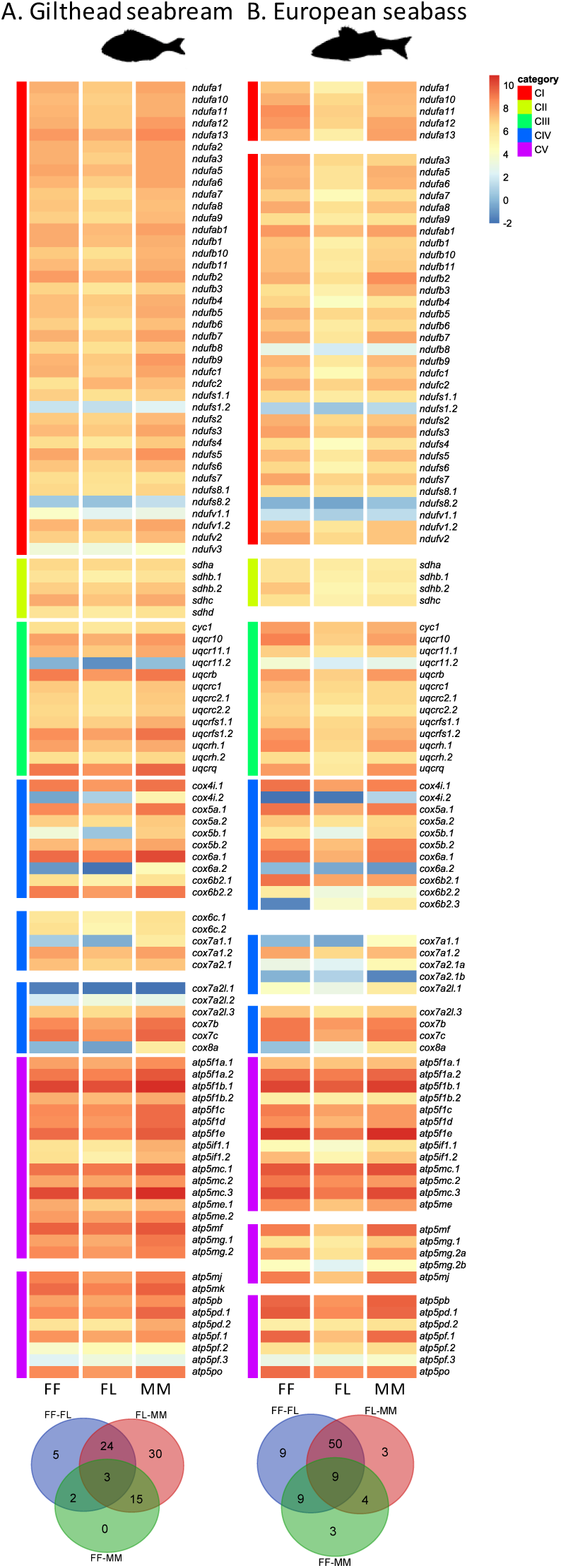
Heat map of gene expression of all OXPHOS genes and Venn diagrams for DEGs between the developmental stages for A) gilthead seabream and B) European seabass. Color bars on the left denote the OXPHOS complex the gene contributes to. FF, First Feeding; FL, Flexion of notochord; MM, Mid metamorphosis.

The ontogenetic expression patterns of OXPHOS genes in European seabass presented many similarities to gilthead seabream. The majority of genes were expressed at least in one stage with the only exception the gene *cox7a2.1b* for which no transcripts were traced in any stage. Overall, 37 of 50 OXPHOS paralogs appear to be DEGs (Figure 7, Tables S9-S11), at least according to one stage-to-stage comparison. Lower expression at FL stage compared with FF and MM was a widespread pattern.

In general, there were many similarities in the OXPHOS gene expression profiles between the two species. Within each developmental stage, genes were differentially expressed, indicative of different regulation mechanisms in place. The VENN diagrams revealed a big-scale transition in OXPHOS gene expression around the FL stage, evident through the number of DEGS at the comparisons FF-FL and FL-MM (Figure 7). In gilthead seabream 34 and 82 DEGs were identified in FF-FL and FL-MM comparisons, respectively, as opposed to 20 in the FF-MM comparison. In European seabass, the differences in DEG number were even higher with 77 and 66 DEGs identified in FF-FL and FL-MM comparisons, respectively, as opposed to 25 in the FF-MM comparison. This transition in expression around FL was the most prominent in the genes of complex I and complex III in European seabass (Figure 7). Complex-specific differences in expression levels were also observed; the genes of complex V exhibited the highest expression levels, followed by the genes of complex IV, and this observation was consistent in both species (Figure 7).

### 3.6. Expression profiles of OXPHOS paralogs

OXPHOS paralogs showed a variety of expression patterns between and within developmental stages (Figure 8). Three patterns were observed: 1) similar expression levels between paralogs, 2) one paralog is overexpressed compared with the other, and 3) one or more paralogs are expressed at a specific stage while the other is consistently expressed across stages.

**Figure 8.**
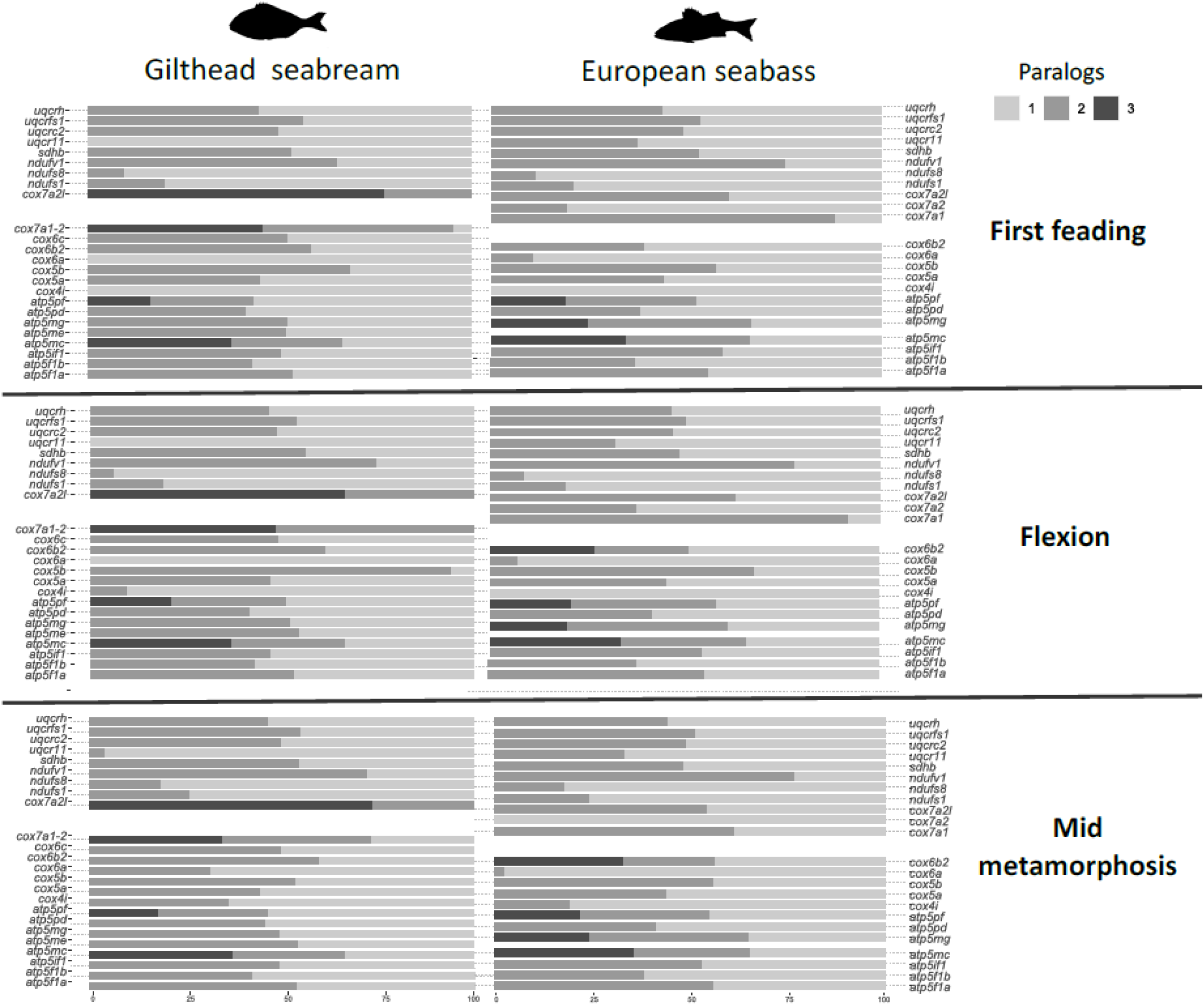
Relative expression levels of each paralog in each gene family expressed as percentage by developmental stage for gilthead seabream (upper panel) and European seabass (lower panel).

A great example of the first pattern was the relative expression of complex III paralogs. The expression levels of the members of the gene families *uqcrh, uqcrfs1*, and *uqcr2* were nearly equal throughout the stages, and this pattern was consistent in both species. The fourth gene family, *uqcr11*, was the exception to the above; while the genes in European seabass shared the same pattern with the other gene families, in gilthead seabream the *uqcr11.2* paralog was not expressed at all during FF and FL and it was only discovered at low levels at the stage of MM. The similar expression levels pattern was also followed by the single gene family of complex II, *sdhb* in both species.

The expression profiles of the gene families of complex I make a good example of the second pattern observed. Complex I has three gene families, *ndufv1*, *ndufs1* and *ndufs8,* with two members each. The genes *ndufv1.2*, *ndufs1.1* and *ndufs8.1* were highly expressed across the stages while their paralogs were expressed at low levels. The paralogs of these families followed the same pattern in both species.

Finally, there were several genes that at least one paralog demonstrated stage specific expression pattern. In European seabass *cox4i.2* was moderately expressed only in MM, while the gilthead seabream homolog was identified earlier in ontogenesis at the stage of FL and increased its levels at MM. In gilthead seabream, the gene *cox6a.2* was not expressed at all at the FF and FL stages and was moderately expressed at MM, whereas in European seabass the homolog was expressed in all three developmental stages but at very low levels. The *cox6b* gene family had only two members in gilthead seabream, which were expressed in all stages, whereas in European seabass one more paralog, the *cox6b.3*, was identified that was expressed only at the later stages of FL and MM.

## 4. Discussion

Gene and genome duplication have significantly altered the OXPHOS gene palette in gilthead seabream and European seabass compared with human and probably terrestrial vertebrates in general. With 24 gene families and a total of 105 genes coding for OXPHOS in gilthead seabream, and 22 gene families and a total of 103 genes coding for OXPHOS in European seabass, compared with 87 in human, and considering those species as representatives for their linages, it is evident that OXPHOS has differentiated significantly between the land vertebrates and teleosts, at least at the genomic level.

### 4.1 Gene duplications modulated OXPHOS genes in gilthead seabream and European seabass

The OXPHOS genes in teleosts appear to have been significantly impacted by TSGD (De Grassi et al., 2005; De Grassi et al., 2008; Little et al., 2010; Porplycia et al., 2017). The majority of paralogs in both species were outgrouped from the single gene of other non-teleost vertebrates and formed two distinct teleost clades. These results indicated that those OXPHOS paralogs diverged from a common ancestor located at the base of teleost lineage. Taking into consideration that those genes can only be identified in teleost species that underwent the teleost-specific genome duplication (TSGD) (Meyer & Van De Peer, 2005; Voldoire et al., 2017), it is proposed that these genes originated from TSGD. The early teleost genome preserved a sizable number of paralogs after the duplication, but afterwards there were species- and/or lineage-specific gene losses, perhaps reflecting the various evolutionary pressures or potential adaptations for the diverse teleost lineages (Brunet et al., 2006; Inoue et al., 2015). Acknowledging the multitude of adaptations seen within the vast radiation of teleosts and their diverse ecological niches, we recognize the presence of more divergent teleost representatives. Given this diversity, it would be intriguing to explore, in the future, the evolution of the OXPHOS gene family in a broader context within the teleost radiation.

### 4.2 Functional Complexity of OXPHOS Subunits

Oxidative phosphorylation (OXPHOS), stands out as one of the most complex processes within the cell due to the intricate coordination of its five multiprotein complexes. Its complexity is evident not only in the specific functions of diverse proteins within these complexes, serving catalytic, structural, regulatory and assembly roles, but also in the collaborative formation of supercomplexes (Habersetzer et al., 2013; Papa et al., 2012; Porplycia et al., 2017; Signes & Fernandez-Vizarra, 2018; Tang et al., 2020; D. F. Wilson, 2017). Notably, gene families with retained paralogs appear to encompass all these functions of OXPHOS subunits. Some interesting observations about the roles of subunits from the identified gene families were recorded. While the majority of complex I subunits are supernumerary, mainly contributing to the assembly and the structure of the complex, none of those subunits have retained paralogs. The subunits coded by the identified gene families (*ndufs1*, *ndufs8*, *ndufv1*) are part of the 7 nuclear encoded core subunits of the complex. Subunits *ndufs1* and *ndufs8* are components of the iron-sulfur fragment that is required in the electron transfer process, while *ndufs1* may form part of the active site crevice where NADH is oxidized. In addition subunit *ndufv1* carries the NADH-binding site as well as flavin mononucleotide (FMN)- and Fe-S-biding sites (Wirth et al., 2016).

Complex III gene families encode catalytic and structural subunits. Subunits *uqcrc11* and *uqcrh* belong to the supernumerary subunits, they act as structural components of the enzyme, and they both play a role at the assembly of the complex. Subunit *uqcrfs1* is the last subunit to be incorporated in complex III, and it’s a member of the catalytic core of the enzyme and essential for enzymatic activity (Fernandez-Vizarra & Zeviani, 2018).

In complex IV, there are eleven different proteins among the nuclear-encoded subunits, with six of them existing in copies in humans. The catalytic core of complex IV is constituted by three mitochondria-encoded subunits (COX1, COX2, COX3), while the nuclear-encoded subunits, although not essential for catalytic function, build the conserved catalytic core. These nuclear-encoded subunits may modulate COX activity under diverse conditions, and some of them have tissue-specific and development-specific isoforms (paralogs), potentially allowing for the fine-tuning of COX function in individual tissues (Čunátová et al., 2020). Between those subunits there were 8 and 7 gene families identified for gilthead seabream and for European seabass respectively, adding to the complexity of teleosts complex IV gene palette.

A number of gene families of complex V were identified in both species under study, sharing between them catalytic structural and assembly roles. Two gene families, *atp5f1a* and *atp5f1b,* code for the water-soluble catalytic domain F1 found in the mitochondrial matrix. In addition, the *atp5mc* gene family codes for F0 domain of the complex where protons enter the matrix through the F0 pore and the F0 motor rotates anticlockwise to turn around F1, thereby driving ATP synthesis. Subunits d and F6 coded by the members of gene families *atp5pd* and *atp5pf* respectively, are part of the peripheral stalk, which connect F1 to F0 and keep the stators from spinning along with the rotor (Neupane et al., 2019; Signes & Fernandez-Vizarra, 2018). Furthermore, probably the most interesting complex V subunits are e and g which are coded by the members of *atp5me* and *atp5mg*. These subunits play a role in the formation of this supramolecular structure. Deleting the genes responsible for these subunits doesn’t impact the assembly or activity of the ATP synthase monomer but specifically influences its dimerization and oligomerization (Habersetzer et al., 2013) that play a role in the creation of cristae. As a result, lack of subunits e and g, and consequently, ATP synthase supercomplexes, leads to alterations in mitochondrial ultrastructure (Habersetzer et al., 2013; Paumard et al., 2002).

In complexes I, II, and III, the subunits coded by the identified gene families were either iron-sulfur proteins (*ndufs1*, *ndufs8*, *sdhd*, and *uqcrfs1*) or featured an Fe-S-binding site (*ndufv1*). Fe– S clusters play an important role in the production of mitochondrial reactive oxygen species. Complex I has two sites, I_F_ (FMN site) and I_Q_ (CoQ binding site), complex II has one site, the II_F_, and complex III has two sites, the Qo and Qi, for ROS production. Overall, the identified OXPHOS gene families encode for subunits that participate in three of the total five ROS production sites of the respiratory chain - *ndufv1* and *ndufs1* are part of the I_F_, *sdhd* is part of the II_F_ and *uqcrfs1* is part of Qo (Mazat et al., 2020). This highlights an additional crucial function of oxidative phosphorylation (OXPHOS) beyond energy generation as ROS play a vital role as intracellular messengers in processes such as proliferation, migration, adhesion, and differentiation (Huo et al., 2009; Ray et al., 2012) apart from the adverse effects of ROS-induced oxidative stress on mitochondria and, consequently, the cell.

For most of the cases, WGDs were the evolutionary origin of that extra genetic material that, after gene losses and gains based on species- or lineage-specific pressures and adaptations, have shaped the OXPHOS gene collection of teleosts as a whole. However, variations were spotted in the gene families and the number of members in each family between the various teleost lineages and even between gilthead seabream and European seabass, which can be considered closely related species. While the number of OXPHOS gene families is high in both species, the majority of OXPHOS genes come in a single copy, suggesting non-functionalization as the fate of most of the OXPHOS paralogs initially generated after the WGDs. For a few of the genes retained in the teleost genomes there was some evidence pointing towards neo functionalization as the possible evolutionary fate. An additional subcellular localization besides mitochondria was identified *in silico* for those genes in both species, while missing protein domains were observed in some of them; both of those findings could be an indication of functional divergence. While the above is a possible scenario, taking into consideration that this evidence is produced by *in silico* analysis and that the genomes of those species are relatively new assemblies with automatic annotation, it is possible that those findings are based on sequencing and/or annotation artifacts and functional evidence is required.

### 4.3 Selective Pressure and Functional Conservation in OXPHOS paralogs

The fact that the majority of identified mutations at the coding region for the genes under study were synonymous substitutions, and no missense substitutions were observed, this suggest that there was selection pressure to preserve the gene and the structure and/or function of the encoded protein. It’s worth noting that a considerable number of the identified mutations were found in the untranslated regions (UTRs) of the transcripts, where genetic alterations do not affect protein function, although they may influence gene regulation. In addition, selection analysis suggested that the mutations identified in the CDS were either neutral or under purifying selection, evidence of pressure to keep the existing protein structure and function by removing mutations that would compromise the protein’s normal activity (Soskine & Tawfik, 2010). Purifying selection suggests that the functional integrity of the protein is critical for the organism, and mutations that alter its function are likely to be harmful. The observed conservation level and purifying selection in the duplicated genes (Nielsen et al., 2010), as well as the fact that these characteristics are similar across gene families, imply that subfunctionalization may be the most plausible evolutionary scenario. The duplicated gene copies may have kept the ancestral functionalities and are susceptible to selective pressure to maintain their respective roles.

### 4.4 Transcriptional Regulation and Developmental Dynamics

The transcriptome analysis in both species provided evidence that the OXPHOS paralogs are functioning genes at least at the transcriptional level. In each species almost all genes were identified at least in one stage. The sole exceptions were the genes *cox7a2.1b* and *cox7a2l.1* for European seabass and gilthead seabream, respectively, which were represented by less than 20 reads and were therefore regarded as not expressed. However, the evidence produced here is not considered adequate to render those paralogs as non-functional, given that the transcriptome analysis covered only a fraction of the species life cycle. Thus, the appearance of those paralogs at later stages cannot be excluded.

The observation of different patterns in the relative expression of the paralogs is indicative of different mechanisms shaping the fate of paralogs. Given that all paralogs encode for proteins that participate in big protein complexes, the need to maintain dosage balance and avoid imbalanced protein stoichiometry is expected to have driven evolutionary decisions right after the WGD event that created them (Almeida-Silva & Van De Peer, 2023; Ishikawa et al., 2020; Picolo et al., 2021). The first pattern that conforms with the dosage balance hypothesis was the most often observed and the genes following this trend exhibited moderate to high expression across all stages. It is possible that these genes are absolutely essential for the overall stability, robustness and function of the complex that their copies have been retained right after the WGD event based on the dosage effect restrains, and dosage balance has generated a time-dependent selective barrier to subfunctionalization or to neofunctionalization (A. E. Wilson & Liberles, 2023).

Paralogs with differentiated relative expression (second pattern) may have been shaped through different pathways starting with the need to maintain dosage balance right after the WGD event. However, subsequent losses of genes or function of the proteins they interact with in their complex might have created the need for a new dosage status (Birchler & Veitia, 2021; Edger & Pires, 2009). The specific needs of the organism might have also added variation in expression patterns. A number of those genes are amongst the catalytic subunits of their corresponding complexes. A great example is the paralogs of complex I, where all three gene families follow this trend in both species. The recruitment of subunits encoded by different paralogs may have an impact on the efficiency of energy production or the availability of that extra material may provide an advantage in rapid changes of the energy needs. The capacity for swift adaptation to new energy demand statuses provides an immense advantage to developing organisms as their organs differentiate and the processes of respiration, feeding and digestion, nutrient absorption and growth mature. This hypothesis is reinforced by the observation that certain paralogs were transcribed only at particular developmental stage. There are numerus examples including myosins, growth factors, opsins and a lot more that stage specific isoforms (paralogs) are known to exist and gene families with different members for embryonic, larvae, juvenile and adult forms (Davies et al., 2011; Georgiou et al., 2016; Gu et al., 2021). It seems that gene paralogs are a crucial feature of fish development and play a significant part in supplying the genetic variety and redundancy needed for the complicated developmental processes involved in the formation of varied fish body plans and adaptations. While all those expression patterns are shared between OXPHOS paralogs, they are not uniformly divided between the five complexes. The paralogs with similar expression levels were mostly members of the gene families of complexes II, III, and V, whereas all the gene families of complex I followed the second pattern with one paralog identified as low expressed, and the majority of complex IV paralogs were following the stage-specific pattern. This observation adds to the evidence that different selective pressures may act on the different OXPHOS complexes driving differentiating adaptations.

This first systematic recording of OXPHOS regulation in fish ontogeny made clear that OXPHOS is not a static process in early fish life. The differential expression analysis identified a number of OXPHOS genes as DEGs between the stage-to-stage comparisons. In both species the comparisons that exhibited the highest number of DEGs were the FF-FL and FL-MM, with the majority of OXPHOS genes identified as DEGs, while the comparison FF-MM had significantly less DEGs again in both species. The fact that numerus genes are identified as DEGs between the developmental stages but a lot of them with low log2 fold change (0.6 - 1), could be an indication of micro adjustments in OXPHOS regulation through the course of development.

Comparing the expression levels between the genes of the five complexes for all developmental stages, stark differences were observed, with the genes coding for the subunits of complex V exhibiting the highest expression. It is known that the ratios of OXPHOS complexes are not 1:1 but closer to 1.1:1.3:3:6.7:3.5 (Hatefi, 1985; Schägger & Pfeiffer, 2001). In our study, genes of complex V were more highly expressed than genes of complex IV, which could be explained by the different species and/or tissue specific needs. The ontogenetic profile of OXPHOS and the up-regulation trend towards metamorphosis may reflect the transition from larva to juvenile form, which comes along with major morphological, histological and behavioral changes and is considered a high energy demanding transition (Metcalfe & Monaghan, 2001; Olivotto et al., 2011). Mid metamorphosis was the stage when the paralogs with stage specific patterns were observed, mostly evident at the complex IV paralogs. The existence of paralogs may add one more level of regulation to the already tightly regulated landscape of cell divisions, migrations and differentiation, driving these dramatic changes and the ultimate transition to the adult form through metamorphosis.

## 5. Conclusions

Our data supports that different gene duplication events, mostly the TSGD, were responsible for the multiple OXPHOS paralogs identified in the gilthead seabream and European seabass genomes and genomic redundancy in respect of dosage balance followed genome duplication The location of mutations, the frequency of synonymous replacements, and the existence of neutral sites or sites under purifying selection, all imply that the paralogs and their proteins have experienced minimal evolutionary alterations. In addition, the retention of mitochondrial targeting sequences and the conservation of peptide structure offer evidence that the majority of OXPHOS paralogs in both species have retained their ancestral function, at least a part of it. The ontogenetic pattern of paralog expression and the marked differences between major developmental stages suggest that the evolution of OXPHOS in teleosts based on the paradigm of gilthead seabream and European seabass is characterized both by plasticity and conservation. The evolutionary constraints appear to have preserved the functional integrity of the OXPHOS machinery, while providing the molecular “space” for adaptations based on regulatory subfunctionalization and the fine tuning of this crucial biological process. This “space” is also recorded in the differences in the number of OXPHOS gene families between the teleost species screened and it reinforces the idea that the fate of the initially duplicated genes is dictated by the evolutionary constraints and adaptations of each species.

## Submission declaration and verification

With the submission of this manuscript, we would like to confirm that the above-mentioned manuscript has not been published elsewhere, accepted for publication elsewhere or under editorial review for publication elsewhere.

## Author contributions

TA: Conceptualization, Data curation, Formal Analysis, Investigation, Methodology, Visualization, Writing – original draft, JCRC: Methodology, Writing – review & editing, AR: Methodology, Visualization, KA: Data curation, Formal Analysis, Visualization DM: Writing – review & editing, Supervision, Methodology, MZ: Writing – review & editing, Supervision, MKA: Conceptualization, Funding acquisition, Project administration, Resources, Supervision, Writing – original draft.

## Acknowledgements

The authors acknowledge HCMR Hellenic Centre for Marine Research for supplying the samples. This work funded by the European Union through project H2020-SFS2016-2-727610 “PerformFISH: Consumer-driven production: Integrating Innovative Approaches for Competitive and Sustainable Performance across the Mediterranean Aquaculture Value Chain”. This study also received Portuguese national funds from FCT - Foundation for Science and Technology through project UIDB/04326/2020 (DOI:10.54499/UIDB/04326/2020), UIDP/04326/2020 (DOI:10.54499/UIDP/04326/2020) and LA/P/0101/2020 (DOI 10.54499/LA/P/0101/2020). This output reflects only the authors; views, and the European Union cannot be held responsible for any use that may be made of the information contained herein.

