## Supplementary material for "Regulatory Subfunctionalization drives OXPHOS evolution in teleosts"

**Table S1.** *Substitution model per gene family*

| Gene | Model |
|---|---|
| *ndufv1* | LG+G |
| *ndufs1* | JTT+G |
| *ndufs8* | T92+G+I |
| *sdhb* | T93+G+I |
| *uqcr11* | K2+G |
| *uqcrc2* | K2+G+I |
| *uqcrfs1* | T92+G+I |
| *uqcrh* | K2+G |
| *cox5a* | K2+G+I |
| *cox5b* | T92+G+I |
| *cox6b2* | T92+G |
| *cox6c* | K2+G+I |
| *cox7a* | LG+G+I |
| *cox4i* | T92+G+I |
| *atp5mc* | GTR+G+I |
| *atp5mg* | T92+G |
| *atp5pd* | K2+G+I |
| *atp5pf* | K2+G+I |
| *atp5f1a* | GTR+G+I |
| *atp5f1b* | GTR+G+I |
| *atp5mea* | K2+G |
| *atp5if1* | K2+G |

***Table S2.*** *Gene IDs and assigned names of OXPHOS genes of gilthead seabream and European seabass*

| ***Sparus aurata*** | | | | ***Dicentrarchus labrax*** | | | |
|---|---|---|---|---|---|---|---|
| Gene ID | Gene name | Gene ID | Gene name | Gene ID | Gene name | Gene ID | Gene name |
| ENSSAUG00010001552 | *atp5f1a.1* | ENSSAUG00010018219 | *ndufa1* | ENSDLAG00005007330 | *atp5f1a.1* | ENSDLAG00005015965 | *ndufa1* |
| ENSSAUG00010008214 | *atp5f1a.2* | ENSSAUG00010027521 | *ndufa10* | ENSDLAG00005018532 | *atp5f1a.2* | ENSDLAG00005008059 | *ndufa10* |
| ENSSAUG00010007970 | *atp5f1b.1* | ENSSAUG00010023201 | *ndufa11* | ENSDLAG00005018338 | *atp5f1b.1* | ENSDLAG00005021448 | *ndufa11* |
| ENSSAUG00010026506 | *atp5f1b.2* | ENSSAUG00010018631 | *ndufa12* | ENSDLAG00005025099 | *atp5f1b.2* | ENSDLAG00005024691 | *ndufa12* |
| ENSSAUG00010022681 | *atp5f1c* | ENSSAUG00010017787 | *ndufa13* | ENSDLAG00005025077 | *atp5f1c* | ENSDLAG00005024977 | *ndufa13* |
| ENSSAUG00010005650 | *atp5f1d* | ENSSAUG00010002150 | *ndufa2* | ENSDLAG00005013665 | *atp5f1d* | ENSDARG00000021984 | *ndufa2* |
| ENSSAUG00010011548 | *atp5f1e* | ENSSAUG00010020916 | *ndufa3* | ENSDLAG00005016988 | *atp5f1e* | ENSDLAG00005002836 | *ndufa3* |
| ENSSAUG00010020382 | *atp5if1.1* | ENSSAUG00010010015 | *ndufa5* | ENSDLAG00005004647 | *atp5if1.1* | ENSDLAG00005008847 | *ndufa5* |
| ENSSAUG00010003151 | *atp5if1.2* | ENSSAUG00010007020 | *ndufa6* | ENSDLAG00005000620 | *atp5if1.2* | ENSDLAG00005015610 | *ndufa6* |
| ENSSAUG00010024766 | *atp5mc.1* | ENSSAUG00010007465 | *ndufa7* | ENSDLAG00005029965 | *atp5mc.1* | ENSDLAG00005025335 | *ndufa7* |

| | | | | | | | |
|---|---|---|---|---|---|---|---|
| ENSSAUG00010001003 | *atp5mc.2* | ENSSAUG00010022401 | *ndufa8* | ENSDLAG00005027449 | *atp5mc.2* | ENSDLAG00005022483 | *ndufa8* |
| ENSSAUG00010003390 | *atp5mc.3* | ENSSAUG00010008168 | *ndufa9* | ENSDLAG00005033252 | *atp5mc.3* | ENSDLAG00005015966 | *ndufa9* |
| ENSSAUG00010022955 | *atp5me.1* | ENSSAUG00010021118 | *ndufab1* | ENSDLAG00005030491 | *atp5me* | ENSDLAG00005001413 | *ndufab1* |
| ENSSAUG00010005603 | *atp5me.2* | ENSSAUG00010022839 | *ndufb1* | ENSDLAG00005007485 | *atp5mf* | ENSDLAG00005001413 | *ndufb1* |
| ENSSAUG00010015404 | *atp5mf* | ENSSAUG00010023733 | *ndufb10* | ENSDLAG00005019335 | *atp5mg.1* | ENSDLAG00005011028 | *ndufb10* |
| ENSSAUG00010011394 | *atp5mg.1* | ENSSAUG00010012559 | *ndufb11* | ENSDLAG00005006347 | *atp5mg.2a* | ENSDLAG00005019348 | *ndufb11* |
| ENSSAUG00010004998 | *atp5mg.2* | ENSSAUG00010013498 | *ndufb2* | ENSDLAG00005006335 | *atp5mg.2b* | ENSDLAG00005019817 | *ndufb2* |
| ENSSAUG00010012064 | *atp5mj* | ENSSAUG00010025848 | *ndufb3* | ENSDLAG00005024092 | *atp5mk* | ENSDLAG00005024263 | *ndufb3* |
| ENSSAUG00010005272 | *atp5mk* | ENSSAUG00010015751 | *ndufb4* | ENSDLAG00005021874 | *atp5pb* | ENSDLAG00005018458 | *ndufb4* |
| ENSSAUG00010001248 | *atp5pb* | ENSSAUG00010027131 | *ndufb5* | ENSDLAG00005003781 | *atp5pd.1* | ENSDLAG00005026097 | *ndufb5* |
| ENSSAUG00010003235 | *atp5pd.1* | ENSSAUG00010018717 | *ndufb6* | ENSDLAG00005021473 | *atp5pd.2* | ENSDLAG00005008294 | *ndufb6* |
| ENSSAUG00010027145 | *atp5pd.2* | ENSSAUG00010018909 | *ndufb7* | ENSDLAG00005015257 | *atp5pf.1* | ENSDLAG00005020963 | *ndufb7* |
| ENSSAUG00010017260 | *atp5pf.1* | ENSSAUG00010023063 | *ndufb8* | ENSDLAG00005000158 | *atp5pf.2* | ENSDLAG00005022803 | *ndufb8* |
| ENSSAUG00010001316 | *atp5pf.2* | ENSSAUG00010000482 | *ndufb9* | ENSDLAG00005015632 | *atp5pf.3* | ENSDLAG00005022603 | *ndufb9* |
| ENSSAUG00010012045 | *atp5pf.3* | ENSSAUG00010012757 | *ndufc1* | ENSDLAG00005017184 | *atp5po* | ENSDLAG00005006810 | *ndufc1* |
| ENSSAUG00010004698 | *atp5po* | ENSSAUG00010000182 | *ndufc2* | ENSDLAG00005016960 | *cox4i.1* | ENSDLAG00005024193 | *ndufc2* |
| ENSSAUG00010008794 | *cox4i.1* | ENSSAUG00010006080 | *ndufs1.1* | ENSDLAG00005020602 | *cox4i.2* | ENSDLAG00005025864 | *ndufs1.1* |
| ENSSAUG00010018002 | *cox4i.2* | ENSSAUG00010021399 | *ndufs1.2* | ENSDLAG00005023073 | *cox5a.1* | ENSDLAG00005009594 | *ndufs1.2* |
| ENSSAUG00010022937 | *cox5a.1* | ENSSAUG00010004779 | *ndufs2* | ENSDLAG00005014376 | *cox5a.2* | ENSDLAG00005024309 | *ndufs2* |
| ENSSAUG00010016180 | *cox5a.2* | ENSSAUG00010004194 | *ndufs3* | ENSDLAG00005009814 | *cox5b.1* | ENSDLAG00005019820 | *ndufs3* |
| ENSSAUG00010021488 | *cox5b.1* | ENSSAUG00010021984 | *ndufs4* | ENSDLAG00005014851 | *cox5b.2* | ENSDLAG00005014329 | *ndufs4* |
| ENSSAUG00010012698 | *cox5b.2* | ENSSAUG00010020202 | *ndufs5* | ENSDLAG00005011849 | *cox6a.1* | ENSDLAG00005015202 | *ndufs5* |
| ENSSAUG00010019745 | *cox6a.1* | ENSSAUG00010008813 | *ndufs6* | ENSDLAG00005010421 | *cox6a.2* | ENSDLAG00005002523 | *ndufs6* |
| ENSSAUG00010024406 | *cox6a.2* | ENSSAUG00010010516 | *ndufs7* | ENSDLAG00005001169 | *cox6b2.1* | ENSDLAG00005022657 | *ndufs7* |
| ENSSAUG00010020219 | *cox6b2.1* | ENSSAUG00010003862 | *ndufs8.1* | ENSDLAG00005008900 | *cox6b2.2* | ENSDLAG00005012483 | *ndufs8.1* |
| ENSSAUG00010014837 | *cox6b2.2* | ENSSAUG00010009712 | *ndufs8.2* | ENSDLAG00005001205 | *cox6b2.3* | ENSDLAG00005015082 | *ndufs8.2* |
| ENSSAUG00010005990 | *cox6c.1* | ENSSAUG00010018928 | *ndufv1.1* | ENSDLAG00005031075 | *cox6c* | ENSDLAG00005004491 | *ndufv1.1* |
| ENSSAUG00010005612 | *cox6c.2* | ENSSAUG00010012863 | *ndufv1.2* | ENSDLAG00005012733 | *cox7a1.1* | ENSDLAG00005013355 | *ndufv1.2* |
| ENSSAUG00010002730 | *cox7a1.1* | ENSSAUG00010012337 | *ndufv2* | ENSDLAG00005022694 | *cox7a1.2* | ENSDLAG00005025947 | *ndufv2* |
| ENSSAUG00010007755 | *cox7a1.2* | ENSSAUG00010001379 | *ndufv3* | ENSDLAG00005001713 | *cox7a2.1a* | ENSDLAG00005001080 | *sdha* |
| ENSSAUG00010024392 | *cox7a2.1* | ENSSAUG00010014692 | *sdha* | ENSDLAG00005001717 | *cox7a2.1b* | ENSDLAG00005008624 | *sdhb.1* |
| ENSSAUG00010023051 | *cox7a2l.1* | ENSSAUG00010000613 | *sdhb.1* | ENSDLAG00005007600 | *cox7a2l.1* | ENSDLAG00005016007 | *sdhb.2* |
| ENSSAUG00010023048 | *cox7a2l.2* | ENSSAUG00010011676 | *sdhb.2* | ENSDLAG00005023340 | *cox7a2l.3* | ENSDLAG00005006491 | *sdhc* |
| ENSSAUG00010026799 | *cox7a2l.3* | ENSSAUG00010008991 | *sdhc* | ENSDLAG00005023940 | *cox7b* | ENSDLAG00005034190 | *sdhd* |
| ENSSAUG00010006458 | *cox7b* | ENSSAUG00010007110 | *sdhd* | ENSDLAG00005025866 | *cox7c* | ENSDLAG00005021401 | *uqcr10* |
| ENSSAUG00010024581 | *cox7c* | ENSSAUG00010021526 | *uqcr10* | ENSDLAG00005009958 | *cox8a* | ENSDLAG00005024816 | *uqcr11.1* |
| ENSSAUG00010021868 | *cox8a* | ENSSAUG00010018172 | *uqcr11.1* | ENSDLAG00005010064 | *cyc1* | ENSDLAG00005007897 | *uqcr11.2* |
| ENSSAUG00010006809 | *cyc1* | ENSSAUG00010016776 | *uqcr11.2* | | | ENSDLAG00005019495 | *uqcrb* |
| | | ENSSAUG00010003983 | *uqcrb* | | | ENSDLAG00005019752 | *uqcrc1* |
| | | ENSSAUG00010010721 | *uqcrc1* | | | ENSDLAG00005014442 | *uqcrc2.1* |
| | | ENSSAUG00010002780 | *uqcrc2.1* | | | ENSDLAG00005014897 | *uqcrc2.2* |
| | | ENSSAUG00010022860 | *uqcrc2.2* | | | ENSDLAG00005020221 | *uqcrfs1.1* |
| | | ENSSAUG00010000160 | *uqcrfs1.1* | | | ENSDLAG00005021271 | *uqcrfs1.2* |
| | | ENSSAUG00010025510 | *uqcrfs1.2* | | | ENSDLAG00005003877 | *uqcrh.1* |
| | | ENSSAUG00010009981 | *uqcrh.1* | | | ENSDLAG00005009892 | *uqcrh.2* |

| | | | | | | | |
|---|---|---|---|---|---|---|---|
| | | ENSSAUG00010022471 | *uqcrh.2* | | | ENSDLAG00005001473 | *uqcrq* |
| | | ENSSAUG00010018590 | *uqcrq* | | | | |

***Table S3.*** *Classification of paralogs for both species based on the level of collinearity between the paralogue pairs as provided by MCScanX*

| **Gene name** | **Collinearity** | |
|---|---|---|
| | ***Sparus aurata*** | ***Dicentrarchus labrax*** |
| ***Complex I*** | | |
| *ndufs1* | dispersed | dispersed |
| *ndufs8* | dispersed | dispersed |
| *ndufv1* | dispersed | dispersed |
| ***Complex II*** | | |
| *sdhb* | WGD/segmental | WGD/segmental |
| ***Complex III*** | | |
| *uqcr11* | WGD/segmental | WGD/segmental |
| *uqcrc2* | WGD/segmental | WGD/segmental |
| *uqcrfs1* | dispersed | dispersed |
| *uqcrh* | dispersed | dispersed |
| ***Complex IV*** | | |
| *cox4i* | dispersed | dispersed |
| *cox5a* | dispersed | dispersed |
| *cox5b* | dispersed | dispersed |
| *cox6a* | dispersed | dispersed |
| *cox6b2* | WGD/segmental | WGD/segmental, dispersed |
| *cox6c* | proximal | - |
| *cox7a1* | dispersed, WGD/segmental | dispersed, tandem |
| *cox7a2l* | dispersed, proximal | dispersed |
| ***Complex V*** | | |
| ***atp5f1a*** | WGD/segmental | WGD/segmental |
| ***atp5f1b*** | dispersed | WGD/segmental |
| ***atp5if1*** | dispersed | dispersed |

| | | |
|---|---|---|
| ***atp5mc*** | WGD/segmental, dispersed | WGD/segmental, dispersed |
| ***atp5me*** | dispersed | - |
| ***atp5mg*** | dispersed | dispersed, proximal |
| ***atp5pd*** | dispersed | dispersed |
| ***atp5pf*** | dispersed | dispersed |

***Table S4.*** *Number, location, and selection mode of polymorphic sites for gilthead seabream OXPHOS paralogs. Dashes used when there were not enough unique sequences existed for selection analysis.*

| Gene name | Complex | Purifying | Diversifying | Neutral | Synonymous | Missense | 5UTR | 3UTR | Splice region |
|---|---|---|---|---|---|---|---|---|---|
| ***ndufs1.1*** | CI | 19 | 1 | 18 | 24 | 16 | | 3 | 2 |
| ***ndufs1.2*** | CI | 1 | | 3 | 3 | 1 | | | |
| ***ndufs8.1*** | CI | 2 | | 7 | 5 | 4 | | | |
| ***ndufs8.2*** | CI | - | - | - | | 1 | | 3 | |
| ***ndufv1.1*** | CI | - | - | - | 1 | | | | |
| ***ndufv1.2*** | CI | - | - | - | | | | | |
| ***sdhb.1*** | CII | | | 3 | 1 | 2 | | 1 | |
| ***sdhb.2*** | CII | 3 | | 1 | 4 | | | 1 | 1 |
| ***uqcr11.1*** | CIII | 1 | | 2 | 3 | | 1 | 8 | |
| ***uqcr11.2*** | CIII | - | - | - | 1 | | | | |
| ***uqcrc2.1*** | CIII | - | - | - | | 1 | 2 | 2 | |
| ***uqcrc2.2*** | CIII | 1 | | 4 | 2 | 3 | | 1 | |
| ***uqcrfs1.1*** | CIII | - | - | - | 1 | | | 3 | |
| ***uqcrfs1.2*** | CIII | - | - | - | | | | | |
| ***uqcrh.1*** | CIII | - | - | - | | | 1 | | |
| ***uqcrh.2*** | CIII | - | - | - | 1 | | | 2 | |
| ***cox4i.1*** | CIV | - | - | - | | | | 3 | 2 |
| ***cox4i.2*** | CIV | 1 | | 2 | 2 | 1 | | 4 | |
| ***cox5a.1*** | CIV | 2 | | 3 | 5 | | | | |
| ***cox5a.2*** | CIV | - | - | - | | 1 | | 3 | |
| ***cox5b.1*** | CIV | - | - | - | 1 | | | | |
| ***cox5b.2*** | CIV | 7 | | 1 | 7 | 1 | 2 | 16 | 1 |

| | | | | | | | | | |
|---|---|---|---|---|---|---|---|---|---|
| ***cox6a.1*** | CIV | 1 | | 2 | 2 | 1 | | 7 | |
| ***cox6a.2*** | CIV | - | - | - | | | | | |
| ***cox6b2.1*** | CIV | 3 | | 2 | 4 | 1 | 4 | 5 | |
| ***cox6b2.2*** | CIV | 1 | | 1 | 1 | 1 | | 1 | |
| ***cox6c.1*** | CIV | - | - | - | | | 3 | 2 | 1 |
| ***cox6c.2*** | CIV | - | - | - | | 1 | 3 | | 1 |
| ***cox7a1.1*** | CIV | 3 | | 2 | 4 | | | 5 | |
| ***cox7a1.2*** | CIV | 1 | | 1 | 1 | 1 | 1 | | |
| ***cox7a2.1*** | CIV | - | - | - | | | | | |
| ***cox7a2l.1*** | CIV | - | - | - | 1 | | | 2 | |
| ***cox7a2l.2*** | CIV | - | - | - | 1 | | 1 | | |
| ***cox7a2l.3*** | CIV | 1 | | 3 | 4 | | | 2 | 1 |
| ***atp5f1a.1*** | CV | 4 | | 3 | 4 | 3 | | 11 | |
| ***atp5f1a.2*** | CV | 6 | | 3 | 8 | 1 | | 2 | 1 |
| ***atp5f1b.1*** | CV | 10 | 1 | 5 | 13 | 4 | | 6 | 1 |
| ***atp5f1b.2*** | CV | 2 | | | 2 | | 2 | | |
| ***atp5if1.1*** | CV | - | - | - | | 1 | 5 | 1 | |
| ***atp5if1.2*** | CV | - | - | - | 1 | | | 2 | |
| ***atp5mc.1*** | CV | | | 2 | | 2 | | 1 | 1 |
| ***atp5mc.2*** | CV | - | - | - | | | | | |
| ***atp5mc.3*** | CV | - | - | - | 1 | | | | |
| ***atp5me.1*** | CV | - | - | - | | | | | |
| ***atp5me.2*** | CV | 3 | | 2 | 5 | | 4 | 36 | |
| ***atp5mg.1*** | CV | - | - | - | | | 1 | 2 | 1 |
| ***atp5mg.2*** | CV | - | - | - | | | | | |
| ***atp5pd.1*** | CV | - | - | - | | 1 | | | |
| ***atp5pd.2*** | CV | 3 | | 4 | 7 | | 7 | 5 | |
| ***atp5pf.1*** | CV | 2 | | 4 | 5 | 1 | 2 | 1 | 1 |
| ***atp5pf.2*** | CV | 24 | 7 | 20 | 38 | 14 | 3 | 5 | |
| ***atp5pf.3*** | CV | 13 | 9 | 31 | 19 | 36 | | 2 | |

***Table S5.*** *Number, location, and selection mode of polymorphic sites for European seabass OXPHOS paralogs. Dashes used when there were not enough unique sequences existed for selection analysis.*

| Gene name | Complex | Purifying | Diversifying | Neutral | Synonymous | Missense | 5UTR | 3UTR | Splice region |
|---|---|---|---|---|---|---|---|---|---|
| ***ndufs1.1*** | CI | 5 | | 7 | 10 | 3 | | | |
| ***ndufs1.2*** | CI | 5 | | 3 | 7 | 1 | | 2 | |
| ***ndufs8.1*** | CI | 1 | | 1 | 2 | | | 2 | 1 |
| ***ndufs8.2*** | CI | 2 | | 8 | 8 | 2 | | 2 | |
| ***ndufv1.1*** | CI | | - | - | | | | | |
| ***ndufv1.2*** | CI | 5 | | 2 | 6 | 1 | | 3 | |
| ***sdhb.1*** | CII | | | 2 | 2 | | | | |
| ***sdhb.2*** | CII | | | | 1 | | | | |
| ***uqcr11.1*** | CIII | | | | 1 | | 3 | 4 | |
| ***uqcr11.2*** | CIII | | - | - | | | | | |
| ***uqcrc2.1*** | CIII | 2 | | 2 | 3 | 1 | | | |
| ***uqcrc2.2*** | CIII | | | 2 | 1 | 1 | | | |
| ***uqcrfs1.1*** | CIII | | - | - | | | | | |
| ***uqcrfs1.2*** | CIII | 2 | | 5 | 5 | 2 | | | |
| ***uqcrh.1*** | CIII | | - | - | | | | | |
| ***uqcrh.2*** | CIII | | | 5 | 3 | 2 | | 4 | |
| ***cox4i.1*** | CIV | 1 | | 1 | 1 | 1 | 1 | | |
| ***cox4i.2*** | CIV | 1 | | 1 | 2 | | | 4 | |
| ***cox5a.1*** | CIV | 1 | | 1 | 2 | | | 1 | |
| ***cox5a.2*** | CIV | | | | 1 | | | | |
| ***cox5b.1*** | CIV | | | 5 | 3 | 2 | 1 | 1 | |
| ***cox5b.2*** | CIV | 1 | | 1 | 2 | 1 | | | |
| ***cox6a.1*** | CIV | | | | 1 | | | 1 | |
| ***cox6a.2*** | CIV | | | 1 | 1 | | | 6 | |
| ***cox6b2.1*** | CIV | - | - | - | | | | 3 | |
| ***cox6b2.2*** | CIV | 1 | | 1 | 1 | 1 | | | |
| ***cox6b2.3*** | CIV | | | | | 2 | | 1 | |
| ***cox7a1.1*** | CIV | 1 | | | 1 | | | | |
| ***cox7a1.2*** | CIV | | | 3 | 1 | 2 | | 5 | |
| ***cox7a2.1a*** | CIV | | | | | 1 | 2 | | |

| | | | | | | | | |
|---|---|---|---|---|---|---|---|---|
| ***cox7a2.1b*** | CIV | - | - | - | | | | |
| ***cox7a2l.1*** | CIV | - | - | - | | | | |
| ***cox7a2l.3*** | CIV | | | | 1 | | | 1 |
| ***atp5f1a.1*** | CV | 3 | | 5 | 7 | 1 | | |
| ***atp5f1a.2*** | CV | 1 | | 3 | 4 | | | 3 |
| ***atp5f1b.1*** | CV | - | - | - | | | | |
| ***atp5f1b.2*** | CV | 5 | | 2 | 7 | | | 3 |
| ***atp5if1.1*** | CV | - | - | - | | | 3 | 1 |
| ***atp5if1.2*** | CV | - | - | - | | | | 3 |
| ***atp5mc.1*** | CV | 1 | | 2 | 3 | | | 3 |
| ***atp5mc.2*** | CV | - | - | - | | | | |
| ***atp5mc.3*** | CV | | | | 1 | | | 2 |
| ***atp5mg.1*** | CV | | | | 1 | | | 1 |
| ***atp5mg.2a*** | CV | | | | 1 | | 1 | |
| ***atp5mg.2b*** | CV | | | 3 | 2 | 1 | | |
| ***atp5pd.1*** | CV | - | - | - | | | | |
| ***atp5pd.2*** | CV | - | - | - | | | | 2 |
| ***atp5pf.1*** | CV | | | 2 | 1 | 1 | | |
| ***atp5pf.2*** | CV | 2 | | 3 | 3 | 2 | 4 | 2 |
| ***atp5pf.3*** | CV | | | 2 | | 2 | | |

***Table S6.*** *Gilthead seabream OXPHOS DEGs between first feeding (FF) and flexion (FL) stages*

| Gene ID | logFC | logCPM | PValue | FDR | neglog10FDR | DEG | gene_name | assigned_name | OXPHOS complex |
|---|---|---|---|---|---|---|---|---|---|
| ENSSAUG00010023201 | -0.60838 | 6.76655 | 0.003721 | 0.015471225 | 1.810475294 | **DOWN** | NDUFA11 | *ndufa11* | CI |
| ENSSAUG00010018631 | -0.65729 | 6.958678 | 0.001683 | 0.007861031 | 2.104520501 | **DOWN** | NDUFA12 | *ndufa12* | CI |
| ENSSAUG00010020916 | -0.8491 | 6.394953 | 6.05E-05 | 0.000435645 | 3.360867736 | **DOWN** | NDUFA3 | *ndufa3* | CI |
| ENSSAUG00010007465 | -0.79699 | 6.61397 | 0.000155 | 0.000996071 | 3.001709531 | **DOWN** | NDUFA7 | *ndufa7* | CI |
| ENSSAUG00010023733 | -0.6547 | 6.822671 | 0.001791 | 0.008293791 | 2.081246903 | **DOWN** | NDUFB10 | *ndufb10* | CI |
| ENSSAUG00010013498 | -0.677 | 7.401605 | 0.001149 | 0.005676951 | 2.245884879 | **DOWN** | NDUFB2 | *ndufb2* | CI |
| ENSSAUG00010025848 | -0.71878 | 6.686957 | 0.000629 | 0.003367447 | 2.472699217 | **DOWN** | NDUFB3 | *ndufb3* | CI |
| ENSSAUG00010018909 | -0.60662 | 7.081087 | 0.00367 | 0.015275237 | 1.816012039 | **DOWN** | NDUFB7 | *ndufb7* | CI |
| ENSSAUG00010012757 | -0.73931 | 6.739576 | 0.000434 | 0.002440533 | 2.612515383 | **DOWN** | NDUFC1 | *ndufc1* | CI |
| ENSSAUG00010000182 | 1.287757 | 7.95149 | 8.24E-10 | 1.75E-08 | 7.755780885 | **UP** | NDUFC2 | *ndufc2* | CI |
| ENSSAUG00010021984 | -0.71608 | 6.959896 | 0.000627 | 0.003359146 | 2.47377114 | **DOWN** | ndufs4 | *ndufs4* | CI |
| ENSSAUG00010018928 | -1.62518 | 4.752917 | 1.69E-12 | 5.56E-11 | 10.25479479 | **DOWN** | NDUFV1 | *ndufv1.1* | CI |
| ENSSAUG00010000613 | -0.75775 | 6.641202 | 0.000319 | 0.001867339 | 2.728776785 | **DOWN** | SDHB | *sdhb.1* | CII |
| ENSSAUG00010008991 | -0.60111 | 7.503902 | 0.003826 | 0.015840092 | 1.800242307 | **DOWN** | SDHC | *sdhc* | CII |
| ENSSAUG00010003983 | -0.6081 | 8.083334 | 0.003296 | 0.013959407 | 1.855133027 | **DOWN** | UQCRB | *uqcrb* | CIII |
| ENSSAUG00010009981 | -0.79974 | 7.113707 | 0.000132 | 0.000865331 | 3.062817713 | **DOWN** | UQCRH | *uqcrh.1* | CIII |
| ENSSAUG00010018590 | -0.61284 | 7.932847 | 0.003091 | 0.013220967 | 1.878736787 | **DOWN** | UQCRQ | *uqcrq* | CIII |
| ENSSAUG00010008794 | -0.73361 | 8.80013 | 0.000382 | 0.002189293 | 2.659696197 | **DOWN** | Cox4a | *cox4i.1* | CIV |
| ENSSAUG00010018002 | 1.744559 | 0.209029 | 0.00179 | 0.008292567 | 2.081311003 | **UP** | Cox4b | *cox4i.2* | CIV |
| ENSSAUG00010022937 | -1.15388 | 7.943167 | 3.36E-08 | 5.26E-07 | 6.279155231 | **DOWN** | COX5a2 | *cox5a.1* | CIV |
| ENSSAUG00010021488 | -3.19077 | 2.26313 | 1.57E-19 | 1.16E-17 | 16.93542513 | **DOWN** | COX5b1 | *cox5b.1* | CIV |
| ENSSAUG00010020219 | -1.19595 | 5.211188 | 6.66E-08 | 9.77E-07 | 6.010095673 | **DOWN** | cox6b2 | *cox6b2.1* | CIV |
| ENSSAUG00010014837 | -0.68203 | 8.423963 | 0.000969 | 0.004895556 | 2.310198009 | **DOWN** | cox6b2 | *cox6b2.2* | CIV |

| | | | | | | | | | |
|---|---|---|---|---|---|---|---|---|---|
| ENSSAUG00010005990 | -0.73822 | 4.943381 | 0.00086 | 0.004411896 | 2.355374712 | **DOWN** | COX6c1 | *cox6c.1* | CIV |
| ENSSAUG00010005612 | -1.44712 | 5.252524 | 7.83E-11 | 1.98E-09 | 8.703058983 | **DOWN** | COX6c1 | *cox6c.2* | CIV |
| ENSSAUG00010007755 | -0.68023 | 7.170087 | 0.00112 | 0.005551347 | 2.2556016 | **DOWN** | COX7a2 | *cox7a1.2* | CIV |
| ENSSAUG00010023048 | 1.283496 | 2.473303 | 1.37E-05 | 0.000117078 | 3.93152438 | **UP** | - | *cox7a2l.2* | CIV |
| ENSSAUG00010006458 | -0.74144 | 8.033359 | 0.000349 | 0.002023832 | 2.693825598 | **DOWN** | - | *cox7b* | CIV |
| ENSSAUG00010024581 | -0.7192 | 8.002034 | 0.000522 | 0.002867575 | 2.542485263 | **DOWN** | COX7c | *cox7c* | CIV |
| ENSSAUG00010003151 | -1.11451 | 5.732481 | 2.68E-07 | 3.47E-06 | 5.460127927 | **DOWN** | atp5if1b | *atp5if1.2* | CV |
| ENSSAUG00010022955 | -0.91205 | 5.995557 | 1.99E-05 | 0.000163118 | 3.787498189 | **DOWN** | atp5mea | *atp5me.1* | CV |
| ENSSAUG00010012064 | -0.68817 | 7.378575 | 0.000954 | 0.004826685 | 2.316351061 | **DOWN** | - | *atp5mj* | CV |
| ENSSAUG00010001316 | 0.857325 | 6.760232 | 4.68E-05 | 0.000348307 | 3.458037353 | **UP** | si:ch211-140m22.7 | *atp5pf.2* | CV |
| ENSSAUG00010012045 | 1.018596 | 5.146167 | 4.24E-06 | 4.16E-05 | 4.380951318 | **UP** | - | *atp5pf.3* | CV |

***Table S7.*** *Gilthead seabream OXPHOS DEGs between flexion (FL) and mid metamorphosis (MM) stages*

| Gene ID | logFC | logCPM | PValue | FDR | neglog10FDR | DEG | gene_name | assigned_name | OXPHOS complex |
|---|---|---|---|---|---|---|---|---|---|
| ENSSAUG00010018219 | 0.771216 | 6.717771 | 0.000235 | 0.001104 | 2.957227 | **UP** | NDUFA1 | *ndufa1* | CI |
| ENSSAUG00010027521 | 0.644622 | 7.907949 | 0.001846 | 0.006786 | 2.168374 | **UP** | ndufa10 | *ndufa10* | CI |
| ENSSAUG00010023201 | 0.652024 | 6.32463 | 0.001976 | 0.007202 | 2.142543 | **UP** | NDUFA11 | *ndufa11* | CI |
| ENSSAUG00010018631 | 1.099147 | 6.775862 | 1.81E-07 | 1.69E-06 | 5.772288 | **UP** | NDUFA12 | *ndufa12* | CI |
| ENSSAUG00010017787 | 0.905105 | 7.49443 | 1.40E-05 | 8.95E-05 | 4.048269 | **UP** | NDUFA13 | *ndufa13* | CI |
| ENSSAUG00010002150 | 0.813611 | 6.381413 | 0.000115 | 0.000582 | 3.235436 | **UP** | NDUFA2 | *ndufa2* | CI |
| ENSSAUG00010020916 | 0.959625 | 5.997943 | 6.58E-06 | 4.49E-05 | 4.347421 | **UP** | NDUFA3 | *ndufa3* | CI |
| ENSSAUG00010007020 | 0.882536 | 6.63665 | 2.70E-05 | 0.00016 | 3.795327 | **UP** | NDUFA6 | *ndufa6* | CI |

| | | | | | | | | | |
|---|---|---|---|---|---|---|---|---|---|
| ENSSAUG00010007465 | 1.137219 | 6.36987 | 8.19E-08 | 8.26E-07 | 6.083017 | **UP** | NDUFA7 | *ndufa7* | CI |
| ENSSAUG00010008168 | 0.85443 | 7.624379 | 3.99E-05 | 0.000226 | 3.645654 | **UP** | NDUFA9 | *ndufa9* | CI |
| ENSSAUG00010021118 | 0.618377 | 7.555135 | 0.002877 | 0.009981 | 2.000823 | **UP** | ndufab1b | *ndufab1* | CI |
| ENSSAUG00010023733 | 1.125042 | 6.659024 | 9.81E-08 | 9.75E-07 | 6.011127 | **UP** | NDUFB10 | *ndufb10* | CI |
| ENSSAUG00010012559 | 0.686491 | 6.381765 | 0.001116 | 0.004378 | 2.358699 | **UP** | NDUFA11 | *ndufb11* | CI |
| ENSSAUG00010025848 | 0.87938 | 6.320167 | 3.17E-05 | 0.000184 | 3.734938 | **UP** | NDUFB3 | *ndufb3* | CI |
| ENSSAUG00010027131 | 0.88502 | 6.776047 | 2.48E-05 | 0.000148 | 3.828414 | **UP** | NDUFB5 | *ndufb5* | CI |
| ENSSAUG00010018717 | 0.814084 | 6.024703 | 0.000127 | 0.000635 | 3.197111 | **UP** | NDUFB6 | *ndufb6* | CI |
| ENSSAUG00010018909 | 0.956861 | 6.833796 | 5.17E-06 | 3.63E-05 | 4.440299 | **UP** | NDUFB7 | *ndufb7* | CI |
| ENSSAUG00010023063 | 0.668573 | 6.149064 | 0.001572 | 0.005891 | 2.229834 | **UP** | ndufb8 | *ndufb8* | CI |
| ENSSAUG00010000482 | 1.133814 | 7.394714 | 6.15E-08 | 6.36E-07 | 6.196223 | **UP** | NDUFB9 | *ndufb9* | CI |
| ENSSAUG00010012757 | 0.980987 | 6.426523 | 3.43E-06 | 2.50E-05 | 4.602244 | **UP** | NDUFC1 | *ndufc1* | CI |
| ENSSAUG00010021399 | 0.942584 | 3.241936 | 0.000172 | 0.000834 | 3.079074 | **UP** | NDUFS1 | *ndufs1.2* | CI |
| ENSSAUG00010004194 | 0.849018 | 7.342747 | 4.64E-05 | 0.000259 | 3.586628 | **UP** | ndufs3 | *ndufs3* | CI |
| ENSSAUG00010021984 | 1.100184 | 6.741542 | 1.78E-07 | 1.67E-06 | 5.777538 | **UP** | ndufs4 | *ndufs4* | CI |
| ENSSAUG00010020202 | 0.917909 | 6.855913 | 1.21E-05 | 7.80E-05 | 4.107957 | **UP** | NDUFS5 | *ndufs5* | CI |
| ENSSAUG00010008813 | 0.765986 | 5.952428 | 0.000315 | 0.001425 | 2.846034 | **UP** | NDUFS6 | *ndufs6* | CI |
| ENSSAUG00010009712 | 1.226513 | 0.884088 | 0.001789 | 0.006604 | 2.180217 | **UP** | NDUFS8 | *ndufs8.2* | CI |
| ENSSAUG00010012337 | 0.661651 | 7.031377 | 0.001516 | 0.005698 | 2.244293 | **UP** | ndufv2 | *ndufv2* | CI |
| ENSSAUG00010001379 | 0.740143 | 5.913822 | 0.000503 | 0.002155 | 2.666553 | **UP** | NDUFV3 | *ndufv3* | CI |
| ENSSAUG00010000613 | 0.70026 | 6.136861 | 0.000936 | 0.003732 | 2.428073 | **UP** | SDHB | *sdhb.1* | CII |
| ENSSAUG00010006809 | 0.662778 | 7.955723 | 0.001365 | 0.005194 | 2.284497 | **UP** | Cyc1 | *cyc1* | CIII |
| ENSSAUG00010021526 | 0.694521 | 7.112121 | 0.000865 | 0.003476 | 2.458954 | **UP** | UQCR10 | *uqcr10* | CIII |
| ENSSAUG00010018172 | 0.671483 | 6.78801 | 0.001333 | 0.005088 | 2.293466 | **UP** | UQCR11-B | *uqcr11.1* | CIII |
| ENSSAUG00010003983 | 0.665923 | 7.650416 | 0.001325 | 0.005061 | 2.295766 | **UP** | UQCRB | *uqcrb* | CIII |
| ENSSAUG00010010721 | 0.702831 | 8.617001 | 0.000662 | 0.002753 | 2.560168 | **UP** | uqcrc1 | *uqcrc1* | CIII |

| | | | | | | | | | |
|---|---|---|---|---|---|---|---|---|---|
| ENSSAUG00010022860 | 0.757147 | 7.393519 | 0.000274 | 0.001261 | 2.899274 | **UP** | uqcrc2a | *uqcrc2.2* | CIII |
| ENSSAUG00010000160 | 0.653075 | 7.698811 | 0.001632 | 0.006085 | 2.215767 | **UP** | UQCRFS1 | *uqcrfs1.1* | CIII |
| ENSSAUG00010025510 | 1.152369 | 7.351788 | 3.82E-08 | 4.14E-07 | 6.383079 | **UP** | UQCRFS1 | *uqcrfs1.2* | CIII |
| ENSSAUG00010009981 | 0.611918 | 6.529101 | 0.00356 | 0.012038 | 1.919428 | **UP** | UQCRH | *uqcrh.1* | CIII |
| ENSSAUG00010018590 | 0.950094 | 7.677698 | 5.05E-06 | 3.55E-05 | 4.450039 | **UP** | UQCRQ | *uqcrq* | CIII |
| ENSSAUG00010008794 | 0.869992 | 8.418707 | 2.66E-05 | 0.000158 | 3.801778 | **UP** | Cox4a | *cox4i.1* | CIV |
| ENSSAUG00010018002 | 4.466957 | 3.734163 | 7.38E-50 | 1.59E-47 | 46.79971 | **UP** | Cox4b | *cox4i.2* | CIV |
| ENSSAUG00010022937 | 1.53874 | 7.750779 | 2.95E-13 | 6.83E-12 | 11.16557 | **UP** | COX5a2 | *cox5a.1* | CIV |
| ENSSAUG00010021488 | 6.698974 | 5.151455 | 9.58E-96 | 7.18E-93 | 92.144 | **UP** | COX5b1 | *cox5b.1* | CIV |
| ENSSAUG00010019745 | 1.281683 | 8.712407 | 7.98E-10 | 1.14E-08 | 7.94381 | **UP** | COX6a1 | *cox6a1* | CIV |
| ENSSAUG00010024406 | 10.81198 | 2.166252 | 3.03E-38 | 4.20E-36 | 35.37663 | **UP** | COX6a2 | *cox6a2* | CIV |
| ENSSAUG00010020219 | 1.00298 | 4.609326 | 8.14E-06 | 5.46E-05 | 4.262502 | **UP** | cox6b2 | *cox6b2.1* | CIV |
| ENSSAUG00010014837 | 0.886065 | 8.084874 | 1.95E-05 | 0.00012 | 3.919706 | **UP** | cox6b2 | *cox6b2.2* | CIV |
| ENSSAUG00010005990 | 1.160463 | 4.749528 | 2.27E-07 | 2.09E-06 | 5.678988 | **UP** | COX6c1 | *cox6c.1* | CIV |
| ENSSAUG00010005612 | 1.224155 | 4.621897 | 6.04E-08 | 6.25E-07 | 6.203904 | **UP** | COX6c1 | *cox6c.2* | CIV |
| ENSSAUG00010002730 | 6.441541 | 3.678219 | 2.28E-68 | 8.09E-66 | 65.0918 | **UP** | COX7a1 | *cox7a1.1* | CIV |
| ENSSAUG00010007755 | 0.746639 | 6.742953 | 0.000367 | 0.001631 | 2.787562 | **UP** | COX7a2 | *cox7a1.2* | CIV |
| ENSSAUG00010026799 | 0.993402 | 6.352207 | 2.66E-06 | 1.99E-05 | 4.701727 | **UP** | cox7a2l | *cox7a2l.3* | CIV |
| ENSSAUG00010006458 | 1.211318 | 7.876359 | 6.83E-09 | 8.38E-08 | 7.076527 | **UP** | - | *cox7b* | CIV |
| ENSSAUG00010024581 | 0.914044 | 7.658047 | 1.13E-05 | 7.33E-05 | 4.13499 | **UP** | COX7c | *cox7c* | CIV |
| ENSSAUG00010021868 | 6.975317 | 4.623382 | 1.48E-89 | 9.54E-87 | 86.02036 | **UP** | COX8b | *cox8a* | CIV |
| ENSSAUG00010001552 | 0.645129 | 9.579008 | 0.001713 | 0.00635 | 2.197239 | **UP** | atp5fa1 | *atp5f1a.1* | CV |
| ENSSAUG00010008214 | 0.980939 | 10.55919 | 2.03E-06 | 1.55E-05 | 4.808315 | **UP** | atp5fa1 | *atp5f1a.2* | CV |
| ENSSAUG00010007970 | 0.830788 | 11.54342 | 5.43E-05 | 0.000297 | 3.527221 | **UP** | atp5f1b | *atp5f1b.1* | CV |
| ENSSAUG00010022681 | 0.76334 | 9.174677 | 0.000214 | 0.001017 | 2.99278 | **UP** | ATP5C1 | *atp5f1c* | CV |
| ENSSAUG00010005650 | 1.05242 | 8.46095 | 4.06E-07 | 3.57E-06 | 5.446873 | **UP** | ATP5D | *atp5f1d* | CV |

| | | | | | | | | | |
|---|---|---|---|---|---|---|---|---|---|
| ENSSAUG00010011548 | 0.803622 | 8.136408 | 0.000105 | 0.000538 | 3.269095 | **UP** | ATP5E | *atp5f1e* | CV |
| ENSSAUG00010020382 | 1.666153 | 6.04345 | 1.64E-14 | 4.46E-13 | 12.35099 | **UP** | atp5if1a | *atp5if1.1* | CV |
| ENSSAUG00010003151 | 2.123771 | 6.020761 | 7.07E-22 | 3.68E-20 | 19.43433 | **UP** | atp5if1b | *atp5if1.2* | CV |
| ENSSAUG00010024766 | 0.764596 | 9.594603 | 0.000207 | 0.000984 | 3.007098 | **UP** | ATP5G1 | *atp5mc.1* | CV |
| ENSSAUG00010003390 | 0.86244 | 10.05665 | 2.88E-05 | 0.000169 | 3.771379 | **UP** | ATP5G3 | *atp5mc.3* | CV |
| ENSSAUG00010011394 | 1.319305 | 7.364447 | 3.54E-10 | 5.29E-09 | 8.276236 | **UP** | ATP5L | *atp5mg.1* | CV |
| ENSSAUG00010012064 | 0.815853 | 6.990301 | 9.56E-05 | 0.000496 | 3.304914 | **UP** | - | *atp5mj* | CV |
| ENSSAUG00010005272 | 0.666573 | 7.723071 | 0.001303 | 0.004984 | 2.302439 | **UP** | atp5md | *atp5mk* | CV |
| ENSSAUG00010001248 | 0.634952 | 8.327361 | 0.002107 | 0.007619 | 2.118115 | **UP** | atp5pb | *atp5pb* | CV |
| ENSSAUG00010003235 | 0.90566 | 8.382742 | 1.24E-05 | 8.02E-05 | 4.096036 | **UP** | atp5pd | *atp5pd.1* | CV |
| ENSSAUG00010027145 | 1.799595 | 6.611352 | 5.30E-17 | 1.84E-15 | 14.73635 | **UP** | atp5pd | *atp5pd.2* | CV |
| ENSSAUG00010012045 | -0.66802 | 4.805868 | 0.002729 | 0.009534 | 2.020742 | **DOWN** | - | *atp5pf.3* | CV |

***Table S8.*** *Gilthead seabream OXPHOS DEGs between first feeding (FF) and mid metamorphosis (MM) stages*

| **Gene ID** | **logFC** | **logCPM** | **PValue** | **FDR** | **neglog10FDR** | **DEG** | **gene_name** | **assigned_name** | **OXPHOS complex** |
|---|---|---|---|---|---|---|---|---|---|
| ENSSAUG00010000482 | 0.659065 | 7.569029 | 0.001511 | 0.004871 | 2.312417 | **UP** | NDUFB9 | ndufb9 | CI |
| ENSSAUG00010000182 | 1.016243 | 7.292003 | 1.21E-06 | 7.49E-06 | 5.125621 | **UP** | NDUFC2 | ndufc2 | CI |
| ENSSAUG00010021399 | 0.711764 | 3.338327 | 0.004572 | 0.013148 | 1.88113 | **UP** | NDUFS1 | ndufs1.2 | CI |
| ENSSAUG00010018928 | -1.10834 | 4.433457 | 1.68E-06 | 1.01E-05 | 4.997089 | **DOWN** | NDUFV1 | ndufv1.1 | CI |
| ENSSAUG00010000160 | 0.865791 | 7.627671 | 3.21E-05 | 0.00015 | 3.823841 | **UP** | UQCRFS1 | uqcrfs1.1 | CI |
| ENSSAUG00010018002 | 6.218516 | 3.702216 | 4.39E-63 | 6.06E-61 | 60.21757 | **UP** | Cox4b | cox4i.2 | CIV |
| ENSSAUG00010021488 | 3.521233 | 5.269738 | 7.54E-46 | 6.27E-44 | 43.20265 | **UP** | COX5b1 | cox5b.1 | CIV |

| | | | | | | | | | |
|---|---|---|---|---|---|---|---|---|---|
| ENSSAUG00010019745 | 0.775891 | 8.886962 | 0.000172 | 0.000691 | 3.160642 | **UP** | COX6a1 | cox6a1 | CIV |
| ENSSAUG00010024406 | 6.280244 | 2.212241 | 5.55E-37 | 3.51E-35 | 34.4548 | **UP** | COX6a2 | cox6a2 | CIV |
| ENSSAUG00010002730 | 5.146639 | 3.717386 | 1.55E-54 | 1.67E-52 | 51.77787 | **UP** | COX7a1 | cox7a1.1 | CIV |
| ENSSAUG00010021868 | 6.166504 | 4.643526 | 9.22E-79 | 2.04E-76 | 75.68968 | **UP** | COX8b | cox8a | CIV |
| ENSSAUG00010008214 | 0.67821 | 10.67607 | 0.000967 | 0.003264 | 2.486215 | **UP** | atp5fa1 | atp5f1a.2 | CV |
| ENSSAUG00010022681 | 0.729034 | 9.195353 | 0.000407 | 0.001499 | 2.824085 | **UP** | ATP5C1 | atp5f1c | CV |
| ENSSAUG00010005650 | 0.797383 | 8.556931 | 0.000116 | 0.00048 | 3.318376 | **UP** | ATP5D | atp5f1d | CV |
| ENSSAUG00010020382 | 1.32508 | 6.141802 | 7.25E-10 | 7.42E-09 | 8.129576 | **UP** | atp5if1a | atp5if1.1 | CV |
| ENSSAUG00010003151 | 1.020945 | 6.310654 | 1.52E-06 | 9.17E-06 | 5.037462 | **UP** | atp5if1b | atp5if1.2 | CV |
| ENSSAUG00010024766 | 0.6881 | 9.631272 | 0.00083 | 0.002837 | 2.547087 | **UP** | ATP5G1 | atp5mc.1 | CV |
| ENSSAUG00010011394 | 1.014957 | 7.466486 | 1.20E-06 | 7.42E-06 | 5.129429 | **UP** | ATP5L | atp5mg.1 | CV |
| ENSSAUG00010003235 | 0.686751 | 8.470709 | 0.000888 | 0.003017 | 2.520477 | **UP** | atp5pd | atp5pd.1 | CV |
| ENSSAUG00010027145 | 1.676791 | 6.648144 | 5.15E-15 | 9.73E-14 | 13.01191 | **UP** | atp5pd | atp5pd.2 | CV |

***Table S9.*** *European seabass OXPHOS DEGs between first feeding (FF) and flexion (FL) stages*

| Gene ID | logFC | logCPM | PValue | FDR | neglog10FDR | DEG | gene_name | assigned_name | OXPHOS complex |
|---|---|---|---|---|---|---|---|---|---|
| **ENSDLAG00005001413** | -0.70445 | 8.256581 | 0.000678 | 0.002999 | 2.522988561 | **DOWN** | ndufab1b | ndufab1 | CI |
| **ENSDLAG00005002523** | -1.48555 | 6.126992 | 1.03E-11 | 2.35E-10 | 9.629284241 | **DOWN** | ndufs6 | ndufs6 | CI |
| **ENSDLAG00005002836** | -1.04983 | 6.558052 | 8.13E-07 | 7.49E-06 | 5.125725335 | **DOWN** | ndufa3 | ndufa3 | CI |
| **ENSDLAG00005004491** | -0.68613 | 3.300529 | 0.009363 | 0.028908 | 1.538987047 | **DOWN** | NDUFV1 | ndufv1.1 | CI |
| **ENSDLAG00005006810** | -2.02051 | 5.753171 | 3.58E-19 | 2.24E-17 | 16.65050077 | **DOWN** | - | ndufc1 | CI |

| | | | | | | | | | |
|---|---|---|---|---|---|---|---|---|---|
| **ENSDLAG00005008059** | -0.85669 | 8.455135 | 3.65E-05 | 0.000225 | 3.64816344 | **DOWN** | ndufa10 | ndufa10 | CI |
| **ENSDLAG00005008294** | -1.24766 | 6.486466 | 5.57E-09 | 7.98E-08 | 7.09784577 | **DOWN** | ndufb6 | ndufb6 | CI |
| **ENSDLAG00005008847** | -1.30567 | 6.526013 | 1.08E-09 | 1.76E-08 | 7.75396646 | **DOWN** | ndufa5 | ndufa5 | CI |
| **ENSDLAG00005011028** | -1.26714 | 7.133062 | 2.17E-09 | 3.35E-08 | 7.47446084 | **DOWN** | ndufb10 | ndufb10 | CI |
| **ENSDLAG00005013355** | -1.16739 | 8.363301 | 2.27E-08 | 2.87E-07 | 6.542476368 | **DOWN** | ndufv1 | ndufv1.2 | CI |
| **ENSDLAG00005014329** | -1.90579 | 6.858027 | 1.95E-18 | 1.13E-16 | 15.94594856 | **DOWN** | ndufs4 | ndufs4 | CI |
| **ENSDLAG00005015202** | -1.40378 | 6.432511 | 7.99E-11 | 1.59E-09 | 8.798478693 | **DOWN** | ndufs5 | ndufs5 | CI |
| **ENSDLAG00005015610** | -1.45502 | 6.826454 | 9.89E-12 | 2.26E-10 | 9.646257323 | **DOWN** | ndufa6 | ndufa6 | CI |
| **ENSDLAG00005015965** | -1.7492 | 6.637423 | 8.49E-16 | 3.59E-14 | 13.44444702 | **DOWN** | ndufa1 | ndufa1 | CI |
| **ENSDLAG00005018458** | -2.23475 | 5.991291 | 4.93E-23 | 4.50E-21 | 20.34663221 | **DOWN** | NDUB4 | ndufb4 | CI |
| **ENSDLAG00005019348** | -1.30979 | 6.542041 | 9.48E-10 | 1.57E-08 | 7.805060546 | **DOWN** | ndufb11 | ndufb11 | CI |
| **ENSDLAG00005019817** | -1.12811 | 6.791651 | 1.06E-07 | 1.18E-06 | 5.927707281 | **DOWN** | ndufb2 | ndufb2 | CI |
| **ENSDLAG00005019820** | -0.97414 | 8.203083 | 2.90E-06 | 2.34E-05 | 4.630804753 | **DOWN** | ndufs3 | ndufs3 | CI |
| **ENSDLAG00005020963** | -1.6498 | 7.072826 | 1.28E-14 | 4.48E-13 | 12.34849083 | **DOWN** | ndufb7 | ndufb7 | CI |
| **ENSDLAG00005021448** | -1.44661 | 7.732764 | 7.47E-12 | 1.75E-10 | 9.757829178 | **DOWN** | ndufa11 | ndufa11 | CI |
| **ENSDLAG00005022483** | -1.19363 | 8.198219 | 1.15E-08 | 1.56E-07 | 6.806493613 | **DOWN** | ndufa8 | ndufa8 | CI |
| **ENSDLAG00005022603** | -1.54095 | 7.474925 | 3.93E-13 | 1.12E-11 | 10.95124919 | **DOWN** | ndufb9 | ndufb9 | CI |
| **ENSDLAG00005022657** | -1.02766 | 8.073506 | 8.42E-07 | 7.71E-06 | 5.112695273 | **DOWN** | ndufs7 | ndufs7 | CI |
| **ENSDLAG00005022803** | -0.77658 | 5.896394 | 0.000321 | 0.001545 | 2.811189383 | **DOWN** | ndufb8 | ndufb8 | CI |
| **ENSDLAG00005023654** | -1.72051 | 5.899412 | 8.43E-15 | 3.01E-13 | 12.52158278 | **DOWN** | NDUFB1 | ndufb1 | CI |
| **ENSDLAG00005024193** | -0.87087 | 7.405356 | 3.19E-05 | 0.0002 | 3.699065199 | **DOWN** | ndufc2 | ndufc2 | CI |
| **ENSDLAG00005024263** | -1.02768 | 5.392867 | 3.45E-06 | 2.74E-05 | 4.562698834 | **DOWN** | ndufb3 | ndufb3 | CI |
| **ENSDLAG00005024309** | -0.83068 | 8.745731 | 6.04E-05 | 0.000351 | 3.454096642 | **DOWN** | ndufs2 | ndufs2 | CI |
| **ENSDLAG00005024691** | -1.42753 | 7.517111 | 1.51E-11 | 3.36E-10 | 9.474244881 | **DOWN** | ndufa12 | ndufa12 | CI |
| **ENSDLAG00005024977** | -1.85277 | 6.79397 | 1.57E-17 | 8.14E-16 | 15.08944468 | **DOWN** | NDUFA13 | ndufa13 | CI |
| **ENSDLAG00005025335** | -2.05257 | 5.877343 | 7.42E-20 | 4.99E-18 | 17.30160245 | **DOWN** | ndufa7 | ndufa7 | CI |

| | | | | | | | | | |
|---|---|---|---|---|---|---|---|---|---|
| **ENSDLAG00005025864** | -0.85785 | 8.450899 | 3.56E-05 | 0.00022 | 3.656833182 | **DOWN** | ndufs1 | ndufs1.1 | CI |
| **ENSDLAG00005025947** | -1.15224 | 8.04395 | 3.64E-08 | 4.46E-07 | 6.350476635 | **DOWN** | ndufv2 | ndufv2 | CI |
| **ENSDLAG00005026097** | -0.93144 | 7.34769 | 8.96E-06 | 6.45E-05 | 4.190260138 | **DOWN** | ndufb5 | ndufb5 | CI |
| **ENSDLAG00005006491** | -0.97705 | 7.129747 | 3.46E-06 | 2.74E-05 | 4.561917322 | **DOWN** | sdhc | sdhc | CII |
| **ENSDLAG00005008624** | -0.66892 | 6.552356 | 0.001578 | 0.006253 | 2.203915991 | **DOWN** | SDHB | sdhb.1 | CII |
| **ENSDLAG00005016007** | -1.93826 | 7.352673 | 2.10E-19 | 1.36E-17 | 16.86630644 | **DOWN** | SDHB | sdhb.2 | CII |
| **ENSDLAG00005001473** | -2.10543 | 6.734384 | 9.98E-22 | 7.93E-20 | 19.10055577 | **DOWN** | uqcrq | uqcrq | CIII |
| **ENSDLAG00005003877** | -1.9005 | 6.82499 | 2.63E-18 | 1.50E-16 | 15.82281476 | **DOWN** | uqcrh | uqcrh.1 | CIII |
| **ENSDLAG00005007897** | -1.38456 | 3.348448 | 1.99E-07 | 2.10E-06 | 5.677404691 | **DOWN** | - | uqcr11.2 | CIII |
| **ENSDLAG00005009892** | -0.8792 | 6.376813 | 3.69E-05 | 0.000227 | 3.643565667 | **DOWN** | - | uqcrh.2 | CIII |
| **ENSDLAG00005010064** | -1.03509 | 9.129857 | 6.10E-07 | 5.76E-06 | 5.239435087 | **DOWN** | cyc1 | cyc1 | CIII |
| **ENSDLAG00005014897** | -1.15164 | 7.730468 | 4.02E-08 | 4.87E-07 | 6.312863002 | **DOWN** | uqcrc2a | uqcrc2.2 | CIII |
| **ENSDLAG00005019495** | -1.16702 | 7.313231 | 3.08E-08 | 3.82E-07 | 6.417925835 | **DOWN** | uqcrb | uqcrb | CIII |
| **ENSDLAG00005021271** | -1.38742 | 7.951304 | 4.38E-11 | 9.05E-10 | 9.043471941 | **DOWN** | UQCRFS1 | uqcrfs1.2 | CIII |
| **ENSDLAG00005021401** | -1.73604 | 7.813145 | 3.42E-16 | 1.52E-14 | 13.81864111 | **DOWN** | uqcr10 | uqcr10 | CIII |
| **ENSDLAG00005024816** | -0.85254 | 6.66764 | 5.64E-05 | 0.00033 | 3.481215826 | **DOWN** | - | uqcr11.1 | CIII |
| **ENSDLAG00005001169** | -1.18299 | 8.673532 | 1.41E-08 | 1.87E-07 | 6.729056908 | **DOWN** | cox6b2 | cox6b2.1 | CIV |
| **ENSDLAG00005001205** | 5.409093 | 3.312495 | 5.57E-41 | 1.88E-38 | 37.72592172 | **UP** | CX6B1 | cox6b2.3 | CIV |
| **ENSDLAG00005007600** | -1.01325 | 2.082775 | 0.002237 | 0.008468 | 2.072237343 | **DOWN** | - | cox7a2l.1 | CIV |
| **ENSDLAG00005008900** | -2.10076 | 4.410784 | 1.38E-17 | 7.19E-16 | 15.14310892 | **DOWN** | cox6b2 | cox6b2.2 | CIV |
| **ENSDLAG00005009814** | -2.95887 | 5.18657 | 1.47E-33 | 2.98E-31 | 30.5263622 | **DOWN** | cox5b2 | cox5b.1 | CIV |
| **ENSDLAG00005009958** | 2.338964 | 2.065602 | 2.57E-10 | 4.73E-09 | 8.324800984 | **UP** | - | cox8a | CIV |
| **ENSDLAG00005011849** | -1.41471 | 8.634956 | 1.53E-11 | 3.41E-10 | 9.467820633 | **DOWN** | cox6a1 | cox6a.1 | CIV |
| **ENSDLAG00005014376** | -0.68445 | 7.23364 | 0.001075 | 0.004496 | 2.347195175 | **DOWN** | cox5aa | cox5a.2 | CIV |
| **ENSDLAG00005014851** | -1.18221 | 7.360492 | 2.02E-08 | 2.59E-07 | 6.586772729 | **DOWN** | COX5B | cox5b.2 | CIV |
| **ENSDLAG00005016960** | -1.06324 | 8.970568 | 3.09E-07 | 3.13E-06 | 5.504470677 | **DOWN** | cox4i1 | cox4i.1 | CIV |

| | | | | | | | | | |
|---|---|---|---|---|---|---|---|---|---|
| **ENSDLAG00005022694** | -1.10311 | 8.059628 | 1.30E-07 | 1.42E-06 | 5.848129481 | **DOWN** | - | cox7a1.2 | CIV |
| **ENSDLAG00005023073** | -1.3114 | 8.59349 | 3.68E-10 | 6.60E-09 | 8.180484188 | **DOWN** | COX5A | cox5a.1 | CIV |
| **ENSDLAG00005023340** | -1.08791 | 7.031331 | 2.61E-07 | 2.68E-06 | 5.572346433 | **DOWN** | COX7R | cox7a2l.3 | CIV |
| **ENSDLAG00005023940** | -0.63061 | 8.390518 | 0.002307 | 0.008707 | 2.060147381 | **DOWN** | - | cox7b | CIV |
| **ENSDLAG00005025866** | -1.2104 | 7.9336 | 7.79E-09 | 1.08E-07 | 6.965481308 | **DOWN** | cox7c | cox7c | CIV |
| **ENSDLAG00005000620** | -2.20388 | 6.860376 | 1.47E-23 | 1.42E-21 | 20.84850123 | **DOWN** | atp5if1b | atp5if1.2 | CV |
| **ENSDLAG00005003781** | -0.988 | 9.060952 | 1.90E-06 | 1.60E-05 | 4.79706307 | **DOWN** | atp5pd | atp5pd.1 | CV |
| **ENSDLAG00005004647** | -0.64911 | 4.617687 | 0.004651 | 0.015897 | 1.798685648 | **DOWN** | atp5if1a | atp5if1.1 | CV |
| **ENSDLAG00005006335** | -1.78254 | 3.035939 | 4.69E-10 | 8.22E-09 | 8.084881134 | **DOWN** | - | atp5mg.2b | CV |
| **ENSDLAG00005006347** | -1.80516 | 7.141081 | 4.37E-17 | 2.14E-15 | 14.66887518 | **DOWN** | atp5l | atp5mg.2a | CV |
| **ENSDLAG00005007485** | -1.71922 | 7.42999 | 8.13E-16 | 3.45E-14 | 13.4621238 | **DOWN** | atp5mf | atp5mf | CV |
| **ENSDLAG00005010974** | -1.2756 | 8.022423 | 1.21E-09 | 1.95E-08 | 7.709802314 | **DOWN** | - | atp5me | CV |
| **ENSDLAG00005013665** | -0.98863 | 8.764214 | 1.92E-06 | 1.62E-05 | 4.791652255 | **DOWN** | atp5f1d | atp5f1d | CV |
| **ENSDLAG00005015257** | -1.95292 | 8.103764 | 6.57E-20 | 4.44E-18 | 17.35229757 | **DOWN** | atp5pf | atp5pf.1 | CV |
| **ENSDLAG00005016988** | -1.05843 | 8.149599 | 3.92E-07 | 3.89E-06 | 5.410014104 | **DOWN** | atp5f1e | atp5f1e | CV |
| **ENSDLAG00005017184** | -0.60661 | 9.004443 | 0.003267 | 0.011751 | 1.929909348 | **DOWN** | atp5po | atp5po | CV |
| **ENSDLAG00005021874** | -0.92945 | 9.37781 | 7.07E-06 | 5.22E-05 | 4.282531644 | **DOWN** | atp5pb | atp5pb | CV |
| **ENSDLAG00005024092** | -1.55396 | 7.412036 | 2.64E-13 | 7.73E-12 | 11.11178677 | **DOWN** | atp5md | atp5mk | CV |
| **ENSDLAG00005024589** | -0.89256 | 10.14188 | 1.53E-05 | 0.000105 | 3.980090439 | **DOWN** | atp5mc1 | atp5mc.3 | CV |
| **ENSDLAG00005025077** | -0.70499 | 10.10942 | 0.000615 | 0.002749 | 2.560901333 | **DOWN** | atp5f1c | atp5f1c | CV |

***Table S10.*** *European seabass OXPHOS DEGs between flexion (FL) and mid metamorphosis (MM) stages*

| Gene ID | logFC | logCPM | PValue | FDR | neglog10FDR | DEG | gene_name | assigned_name | OXPHOS complex |
|---|---|---|---|---|---|---|---|---|---|
| ENSDLAG00005002523 | 1.305503 | 5.687814 | 1.98E-09 | 5.86E-08 | 7.232033 | **UP** | ndufs6 | ndufs6 | CI |
| ENSDLAG00005006810 | 1.999907 | 5.428119 | 7.30E-19 | 6.84E-17 | 16.16514 | **UP** | - | ndufc1 | CI |
| ENSDLAG00005008059 | 0.650139 | 8.016166 | 0.001683 | 0.009642 | 2.015851 | **UP** | ndufa10 | ndufa10 | CI |
| ENSDLAG00005008294 | 0.786943 | 5.869568 | 0.000228 | 0.001835 | 2.736388 | **UP** | ndufb6 | ndufb6 | CI |
| ENSDLAG00005008847 | 1.413724 | 6.294578 | 4.07E-11 | 1.60E-09 | 8.7961 | **UP** | ndufa5 | ndufa5 | CI |
| ENSDLAG00005009594 | 0.719364 | 3.121275 | 0.00488 | 0.023062 | 1.637111 | **UP** | NDUFS1 | ndufs1.2 | CI |
| ENSDLAG00005011028 | 1.031958 | 6.661889 | 1.01E-06 | 1.67E-05 | 4.77673 | **UP** | ndufb10 | ndufb10 | CI |
| ENSDLAG00005013355 | 1.035511 | 7.964034 | 6.66E-07 | 1.15E-05 | 4.937461 | **UP** | ndufv1 | ndufv1.2 | CI |
| ENSDLAG00005014329 | 1.634557 | 6.339475 | 4.00E-14 | 2.46E-12 | 11.60828 | **UP** | ndufs4 | ndufs4 | CI |
| ENSDLAG00005015202 | 1.490327 | 6.186711 | 5.11E-12 | 2.29E-10 | 9.640976 | **UP** | ndufs5 | ndufs5 | CI |
| ENSDLAG00005015610 | 1.386018 | 6.467141 | 8.21E-11 | 3.05E-09 | 8.515792 | **UP** | ndufa6 | ndufa6 | CI |
| ENSDLAG00005015965 | 2.12706 | 6.627692 | 2.60E-22 | 3.16E-20 | 19.50054 | **UP** | ndufa1 | ndufa1 | CI |
| ENSDLAG00005018458 | 1.656808 | 5.225019 | 1.31E-13 | 7.39E-12 | 11.13111 | **UP** | NDUB4 | ndufb4 | CI |
| ENSDLAG00005019348 | 1.144852 | 6.117511 | 8.31E-08 | 1.76E-06 | 5.754157 | **UP** | ndufb11 | ndufb11 | CI |
| ENSDLAG00005019817 | 2.071203 | 7.189475 | 7.84E-22 | 9.21E-20 | 19.03581 | **UP** | ndufb2 | ndufb2 | CI |
| ENSDLAG00005019820 | 0.669663 | 7.699378 | 0.001247 | 0.007557 | 2.121664 | **UP** | ndufs3 | ndufs3 | CI |
| ENSDLAG00005020963 | 1.784456 | 6.866862 | 8.79E-17 | 6.93E-15 | 14.15937 | **UP** | ndufb7 | ndufb7 | CI |
| ENSDLAG00005022483 | 0.805961 | 7.630648 | 0.000106 | 0.00097 | 3.013023 | **UP** | ndufa8 | ndufa8 | CI |
| ENSDLAG00005022603 | 1.397929 | 7.060694 | 3.97E-11 | 1.57E-09 | 8.805373 | **UP** | ndufb9 | ndufb9 | CI |
| ENSDLAG00005022803 | 0.641011 | 5.503287 | 0.002998 | 0.015556 | 1.808104 | **UP** | ndufb8 | ndufb8 | CI |
| ENSDLAG00005023654 | 1.404996 | 5.355437 | 1.98E-10 | 6.98E-09 | 8.155926 | **UP** | NDUFB1 | ndufb1 | CI |
| ENSDLAG00005024193 | 0.710288 | 6.994357 | 0.000677 | 0.00457 | 2.340116 | **UP** | ndufc2 | ndufc2 | CI |
| ENSDLAG00005024263 | 2.277893 | 6.027145 | 2.35E-24 | 3.50E-22 | 21.45646 | **UP** | ndufb3 | ndufb3 | CI |

| | | | | | | | | | |
|---|---|---|---|---|---|---|---|---|---|
| ENSDLAG00005024691 | 1.126572 | 6.995004 | 8.68E-08 | 1.83E-06 | 5.736803 | **UP** | ndufa12 | ndufa12 | CI |
| ENSDLAG00005024977 | 2.327198 | 6.868624 | 3.55E-26 | 6.03E-24 | 23.21948 | **UP** | NDUFA13 | ndufa13 | CI |
| ENSDLAG00005025335 | 1.793068 | 5.363768 | 1.13E-15 | 8.22E-14 | 13.08526 | **UP** | ndufa7 | ndufa7 | CI |
| ENSDLAG00005006491 | 0.664461 | 6.621099 | 0.001553 | 0.009003 | 2.045589 | **UP** | sdhc | sdhc | CII |
| ENSDLAG00005001473 | 2.449269 | 6.710147 | 2.35E-28 | 4.58E-26 | 25.33916 | **UP** | uqcrq | uqcrq | CIII |
| ENSDLAG00005003877 | 1.438636 | 6.165096 | 2.55E-11 | 1.03E-09 | 8.985321 | **UP** | uqcrh | uqcrh.1 | CIII |
| ENSDLAG00005009892 | 0.617132 | 5.903605 | 0.003748 | 0.018652 | 1.729268 | **UP** | - | uqcrh.2 | CIII |
| ENSDLAG00005010064 | 0.623337 | 8.557156 | 0.002519 | 0.013484 | 1.870176 | **UP** | cyc1 | cyc1 | CIII |
| ENSDLAG00005014897 | 0.765566 | 7.166484 | 0.000245 | 0.001947 | 2.710695 | **UP** | uqcrc2a | uqcrc2.2 | CIII |
| ENSDLAG00005019495 | 1.267753 | 7.074362 | 1.86E-09 | 5.52E-08 | 7.257712 | **UP** | uqcrb | uqcrb | CIII |
| ENSDLAG00005021271 | 0.873952 | 7.289913 | 2.85E-05 | 0.000314 | 3.502522 | **UP** | UQCRFS1 | uqcrfs1.2 | CIII |
| ENSDLAG00005021401 | 1.100961 | 7.042437 | 1.64E-07 | 3.25E-06 | 5.487892 | **UP** | uqcr10 | uqcr10 | CIII |
| ENSDLAG00005024816 | 0.824832 | 6.340637 | 9.56E-05 | 0.000884 | 3.05347 | **UP** | - | uqcr11.1 | CIII |
| ENSDLAG00005001205 | 1.76026 | 5.078701 | 7.70E-15 | 5.13E-13 | 12.28994 | **UP** | CX6B1 | cox6b2.3 | CIV |
| ENSDLAG00005001713 | 2.340445 | 3.066815 | 6.53E-17 | 5.21E-15 | 14.28302 | **UP** | cox7a2a | cox7a2.1a | CIV |
| ENSDLAG00005001717 | -2.5571 | -1.24886 | 0.009611 | 0.039753 | 1.400625 | **DOWN** | - | cox7a2.1b | CIV |
| ENSDLAG00005007600 | 2.541384 | 2.933389 | 1.57E-18 | 1.46E-16 | 15.83708 | **UP** | - | cox7a2l.1 | CIV |
| ENSDLAG00005009814 | 3.722372 | 5.571764 | 6.98E-51 | 4.27E-48 | 47.36913 | **UP** | cox5b2 | cox5b.1 | CIV |
| ENSDLAG00005009958 | 3.305936 | 4.889453 | 8.42E-40 | 3.26E-37 | 36.48709 | **UP** | - | cox8a | CIV |
| ENSDLAG00005011849 | 1.293799 | 8.238723 | 6.18E-10 | 1.98E-08 | 7.702424 | **UP** | cox6a1 | cox6a.1 | CIV |
| ENSDLAG00005012733 | 4.615523 | 2.874778 | 4.35E-37 | 1.53E-34 | 33.81607 | **UP** | cox7a1 | cox7a1.1 | CIV |
| ENSDLAG00005014851 | 1.560707 | 7.32396 | 1.81E-13 | 1.01E-11 | 10.99761 | **UP** | COX5B | cox5b.2 | CIV |
| ENSDLAG00005016960 | 0.712433 | 8.43357 | 0.000567 | 0.003931 | 2.405445 | **UP** | cox4i1 | cox4i.1 | CIV |
| ENSDLAG00005020602 | 3.34326 | 1.18869 | 3.74E-13 | 1.97E-11 | 10.70449 | **UP** | cox4i2 | cox4i.2 | CIV |
| ENSDLAG00005022694 | 0.710734 | 7.494601 | 0.00063 | 0.004317 | 2.364795 | **UP** | - | cox7a1.2 | CIV |
| ENSDLAG00005023073 | 1.031325 | 8.090279 | 7.23E-07 | 1.24E-05 | 4.906026 | **UP** | COX5A | cox5a.1 | CIV |

| | | | | | | | | | |
|---|---|---|---|---|---|---|---|---|---|
| ENSDLAG00005023340 | 1.024938 | 6.679605 | 1.18E-06 | 1.93E-05 | 4.714597 | **UP** | COX7R | cox7a2l.3 | CIV |
| ENSDLAG00005025866 | 1.263506 | 7.66156 | 1.71E-09 | 5.12E-08 | 7.2911 | **UP** | cox7c | cox7c | CIV |
| ENSDLAG00005000620 | 2.003383 | 6.388881 | 5.50E-20 | 5.70E-18 | 17.24433 | **UP** | atp5if1b | atp5if1.2 | CV |
| ENSDLAG00005003781 | 0.75454 | 8.600669 | 0.000261 | 0.00205 | 2.688221 | **UP** | atp5pd | atp5pd.1 | CV |
| ENSDLAG00005004321 | 0.631083 | 9.720717 | 0.00215 | 0.011862 | 1.925836 | **UP** | - | atp5mc.1 | CV |
| ENSDLAG00005004647 | 2.09098 | 5.339869 | 3.41E-20 | 3.56E-18 | 17.44797 | **UP** | atp5if1a | atp5if1.1 | CV |
| ENSDLAG00005006335 | 1.877528 | 2.804059 | 2.44E-11 | 9.94E-10 | 9.002827 | **UP** | - | atp5mg.2b | CV |
| ENSDLAG00005006347 | 2.086176 | 7.054863 | 4.80E-22 | 5.75E-20 | 19.2406 | **UP** | atp5l | atp5mg.2a | CV |
| ENSDLAG00005007485 | 2.178434 | 7.48516 | 6.19E-24 | 8.80E-22 | 21.05543 | **UP** | atp5mf | atp5mf | CV |
| ENSDLAG00005010974 | 0.958746 | 7.496384 | 4.36E-06 | 6.07E-05 | 4.216478 | **UP** | - | atp5me | CV |
| ENSDLAG00005013665 | 0.739748 | 8.294264 | 0.00035 | 0.002624 | 2.581108 | **UP** | atp5f1d | atp5f1d | CV |
| ENSDLAG00005015257 | 1.918095 | 7.766905 | 2.64E-19 | 2.57E-17 | 16.59065 | **UP** | atp5pf | atp5pf.1 | CV |
| ENSDLAG00005016988 | 1.443251 | 8.111028 | 6.43E-12 | 2.83E-10 | 9.548729 | **UP** | atp5f1e | atp5f1e | CV |
| ENSDLAG00005019335 | 1.218705 | 5.950208 | 1.50E-08 | 3.71E-07 | 6.430965 | **UP** | atp5l | atp5mg.1 | CV |
| ENSDLAG00005021874 | 1.136585 | 9.207519 | 4.42E-08 | 9.96E-07 | 6.001714 | **UP** | atp5pb | atp5pb | CV |
| ENSDLAG00005024092 | 1.875057 | 7.34873 | 1.89E-18 | 1.74E-16 | 15.75987 | **UP** | atp5md | atp5mk | CV |
| ENSDLAG00005024589 | 0.918996 | 9.849702 | 8.57E-06 | 0.00011 | 3.958601 | **UP** | atp5mc1 | atp5mc.3 | CV |

***Table S11.*** *European seabass OXPHOS DEGs between first feeding (FF) and mid metamorphosis (MM) stages*

| Gene ID | logFC | logCPM | PValue | FDR | neglog10FDR | DEG | gene_name | assigned_name | OXPHOS complex |
|---|---|---|---|---|---|---|---|---|---|
| ENSDLAG00005002836 | -0.99366 | 6.292763 | 2.96E-06 | 1.79E-05 | 4.746712 | **DOWN** | ndufa3 | ndufa3 | CI |
| ENSDLAG00005018458 | -0.62893 | 6.149405 | 0.003039 | 0.009588 | 2.01825 | **DOWN** | NDUB4 | ndufb4 | CI |
| ENSDLAG00005019817 | 0.892106 | 7.478111 | 1.93E-05 | 0.000101 | 3.996015 | **UP** | ndufb2 | ndufb2 | CI |
| ENSDLAG00005021448 | -1.08597 | 7.554912 | 2.12E-07 | 1.55E-06 | 5.808687 | **DOWN** | ndufa11 | ndufa11 | CI |
| ENSDLAG00005022657 | -0.67889 | 7.91397 | 0.001058 | 0.003762 | 2.424626 | **DOWN** | ndufs7 | ndufs7 | CI |
| ENSDLAG00005024263 | 1.199223 | 6.253495 | 2.08E-08 | 1.78E-07 | 6.749777 | **UP** | ndufb3 | ndufb3 | CI |
| ENSDLAG00005025947 | -0.75228 | 7.896465 | 0.000289 | 0.001179 | 2.928508 | **DOWN** | ndufv2 | ndufv2 | CI |
| ENSDLAG00005016007 | -1.59726 | 7.146347 | 6.08E-14 | 1.05E-12 | 11.97778 | **DOWN** | SDHB | sdhb.2 | CII |
| ENSDLAG00005007897 | -0.82602 | 3.24294 | 0.001234 | 0.004315 | 2.36503 | **DOWN** | - | uqcr11.2 | CIII |
| ENSDLAG00005021401 | -0.68606 | 7.847987 | 0.000942 | 0.003385 | 2.470442 | **DOWN** | uqcr10 | uqcr10 | CIII |
| ENSDLAG00005001169 | -1.05954 | 8.428544 | 3.50E-07 | 2.47E-06 | 5.606549 | **DOWN** | cox6b2 | cox6b2.1 | CIV |
| ENSDLAG00005001205 | 7.118177 | 4.696071 | 2.92E-80 | 1.05E-77 | 76.97697 | **UP** | CX6B1 | cox6b2.3 | CIV |
| ENSDLAG00005001713 | 1.717545 | 3.172191 | 2.26E-10 | 2.53E-09 | 8.596459 | **UP** | cox7a2a | cox7a2.1a | CIV |
| ENSDLAG00005007600 | 1.477009 | 3.1287 | 4.62E-08 | 3.73E-07 | 6.427825 | **UP** | - | cox7a2l.1 | CIV |
| ENSDLAG00005008900 | -1.66973 | 4.219499 | 2.89E-12 | 4.06E-11 | 10.39133 | **DOWN** | cox6b2 | cox6b2.2 | CIV |
| ENSDLAG00005009814 | 0.712465 | 6.12852 | 0.000809 | 0.002961 | 2.528537 | **UP** | cox5b2 | cox5b.1 | CIV |
| ENSDLAG00005009958 | 5.593796 | 4.759639 | 1.02E-69 | 3.04E-67 | 66.51674 | **UP** | - | cox8a | CIV |
| ENSDLAG00005012733 | 4.040594 | 2.89203 | 3.24E-30 | 2.17E-28 | 27.66409 | **UP** | cox7a1 | cox7a1.1 | CIV |
| ENSDLAG00005020602 | 3.642027 | 1.178278 | 2.87E-13 | 4.64E-12 | 11.33355 | **UP** | cox4i2 | cox4i.2 | CIV |
| ENSDLAG00005004647 | 1.390874 | 5.477727 | 3.17E-10 | 3.49E-09 | 8.457749 | **UP** | atp5if1a | atp5if1.1 | CV |
| ENSDLAG00005015632 | 0.754024 | 2.051071 | 0.016586 | 0.042482 | 1.371795 | **UP** | - | atp5pf.3 | CV |
| ENSDLAG00005019335 | 0.697829 | 6.10246 | 0.001041 | 0.003705 | 2.43116 | **UP** | atp5l | atp5mg.1 | CV |
| ENSDLAG00005021473 | 0.648233 | 6.003445 | 0.002362 | 0.007673 | 2.115057 | **UP** | atp5pd | atp5pd.2 | CV |

| | | | | | | | | | |
|---|---|---|---|---|---|---|---|---|---|
| ENSDLAG00005023876 | -0.77976 | 8.593294 | 0.000163 | 0.000703 | 3.153165 | **DOWN** | atp5mc3b | atp5mc.2 | CV |
| ENSDLAG00005025077 | -0.60516 | 9.864306 | 0.003243 | 0.010162 | 1.993 | **DOWN** | atp5f1c | atp5f1c | CV |

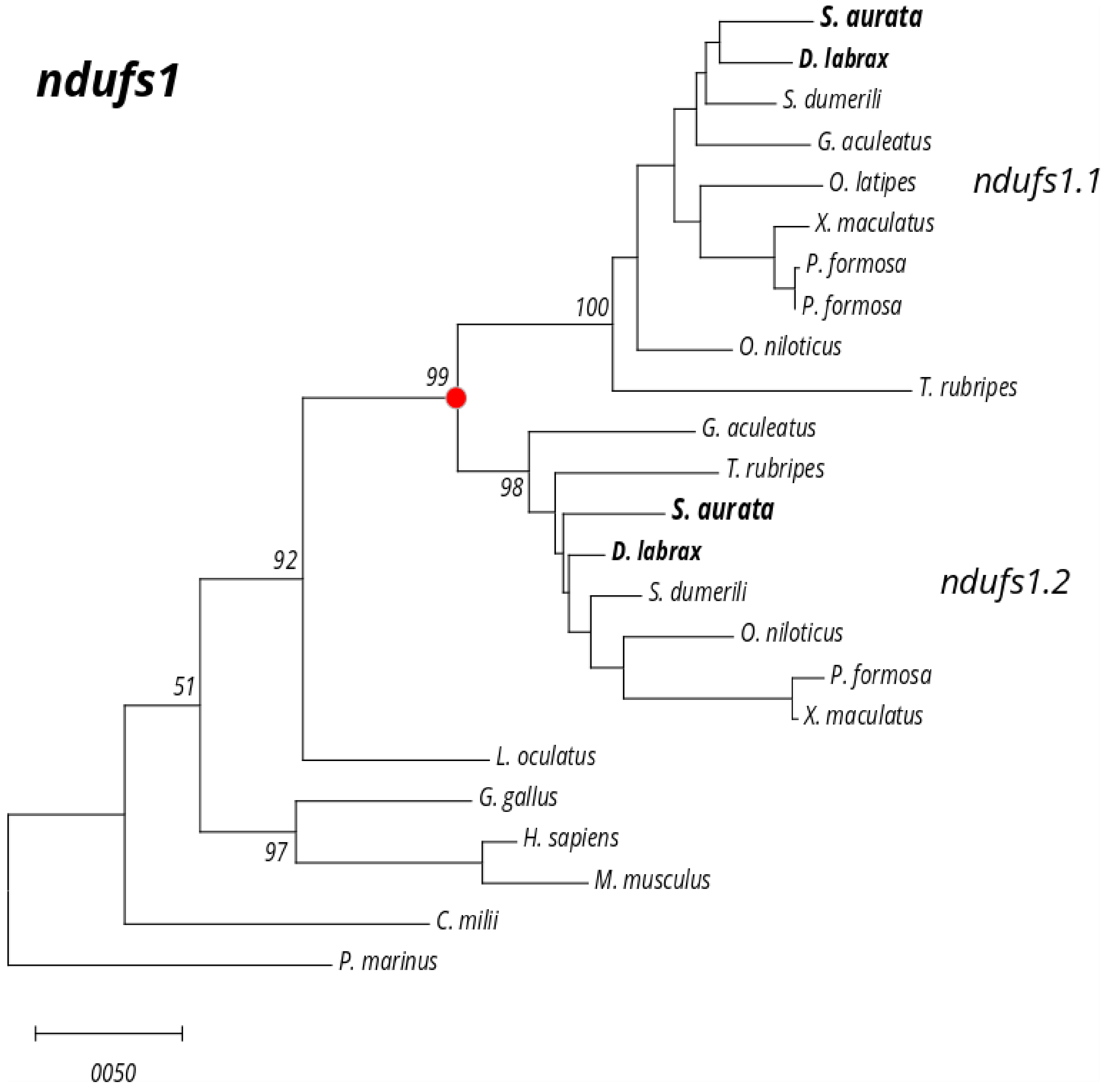

***Figure S1***. *Phylogenetic tree for ndufs1 gene family*

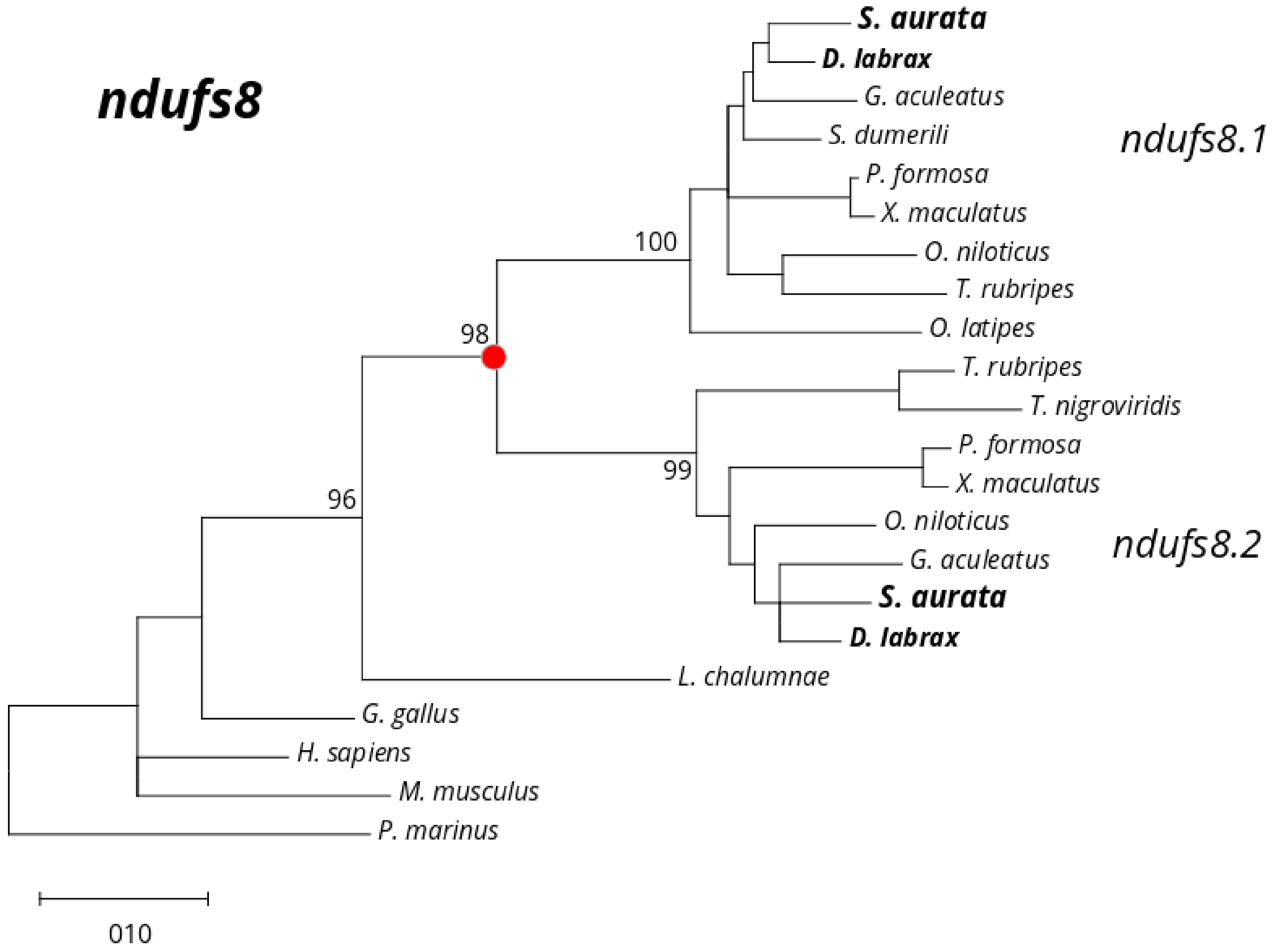

***Figure S2***. *Phylogenetic tree for ndufs8 gene family*

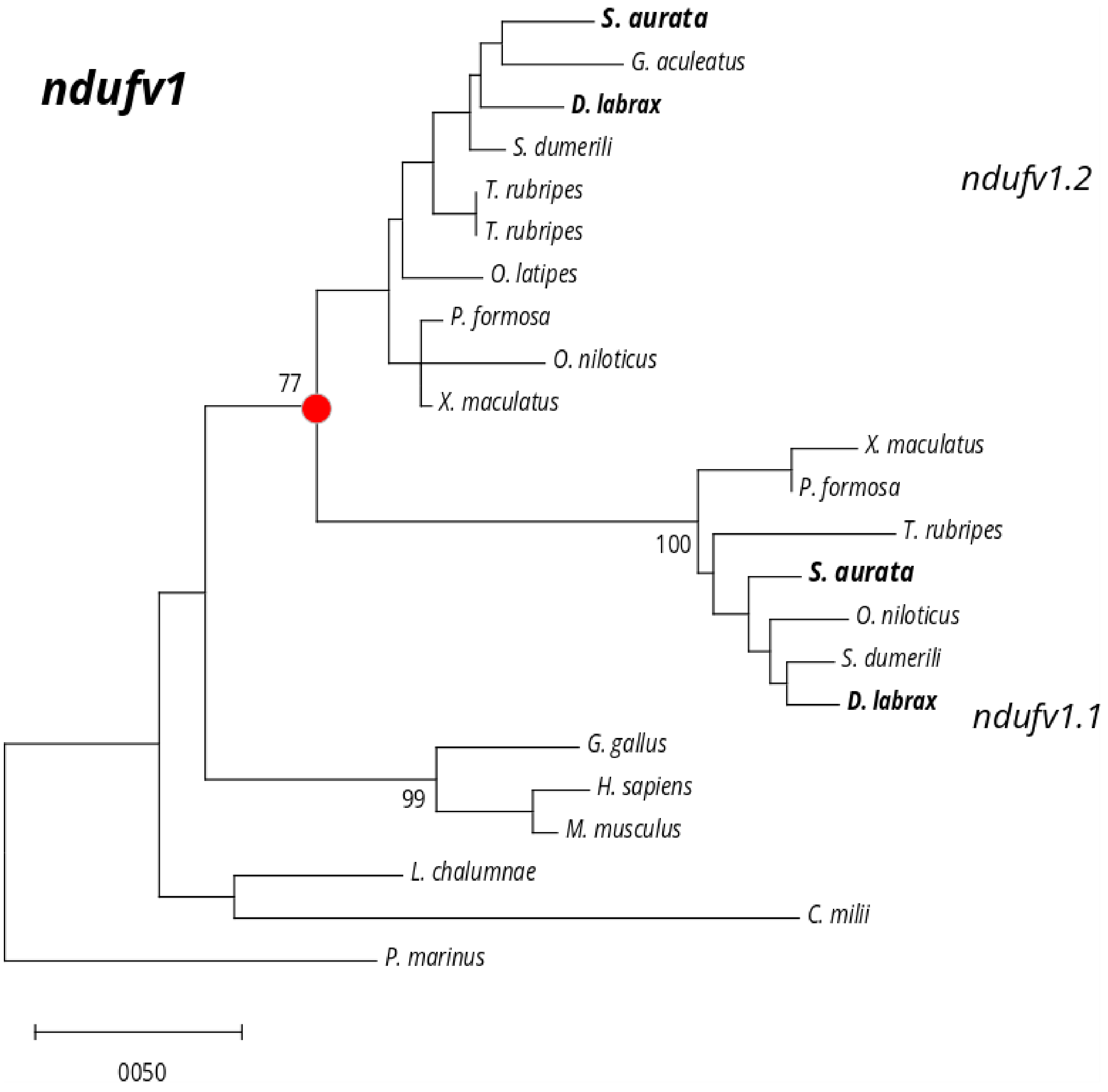

***Figure S3.*** *Phylogenetic tree for ndufv1 gene family*

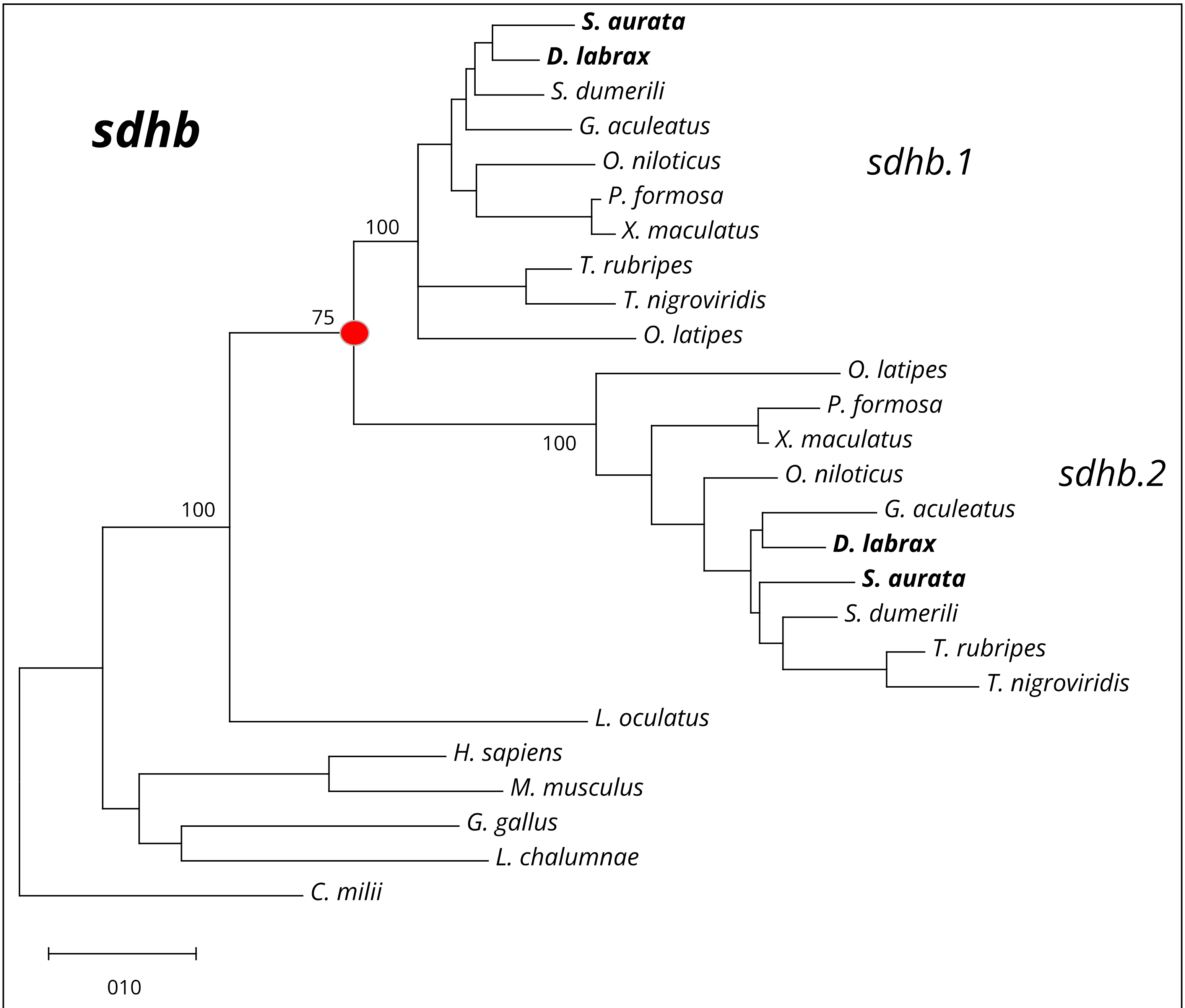

***Figure S4.*** *Phylogenetic tree for sdhb gene family*

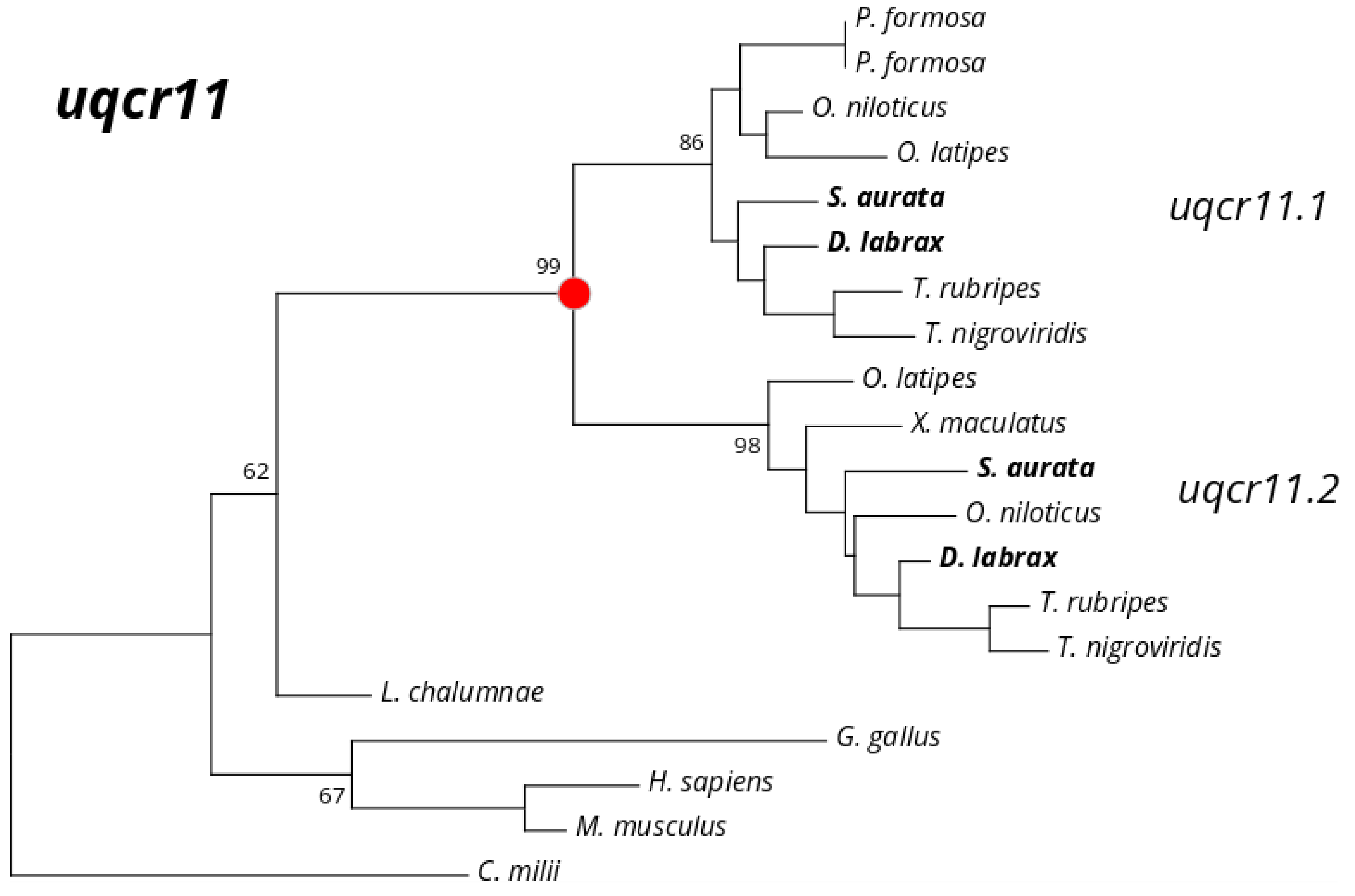

***Figure S5.*** *Phylogenetic tree for uqcr11 gene family*

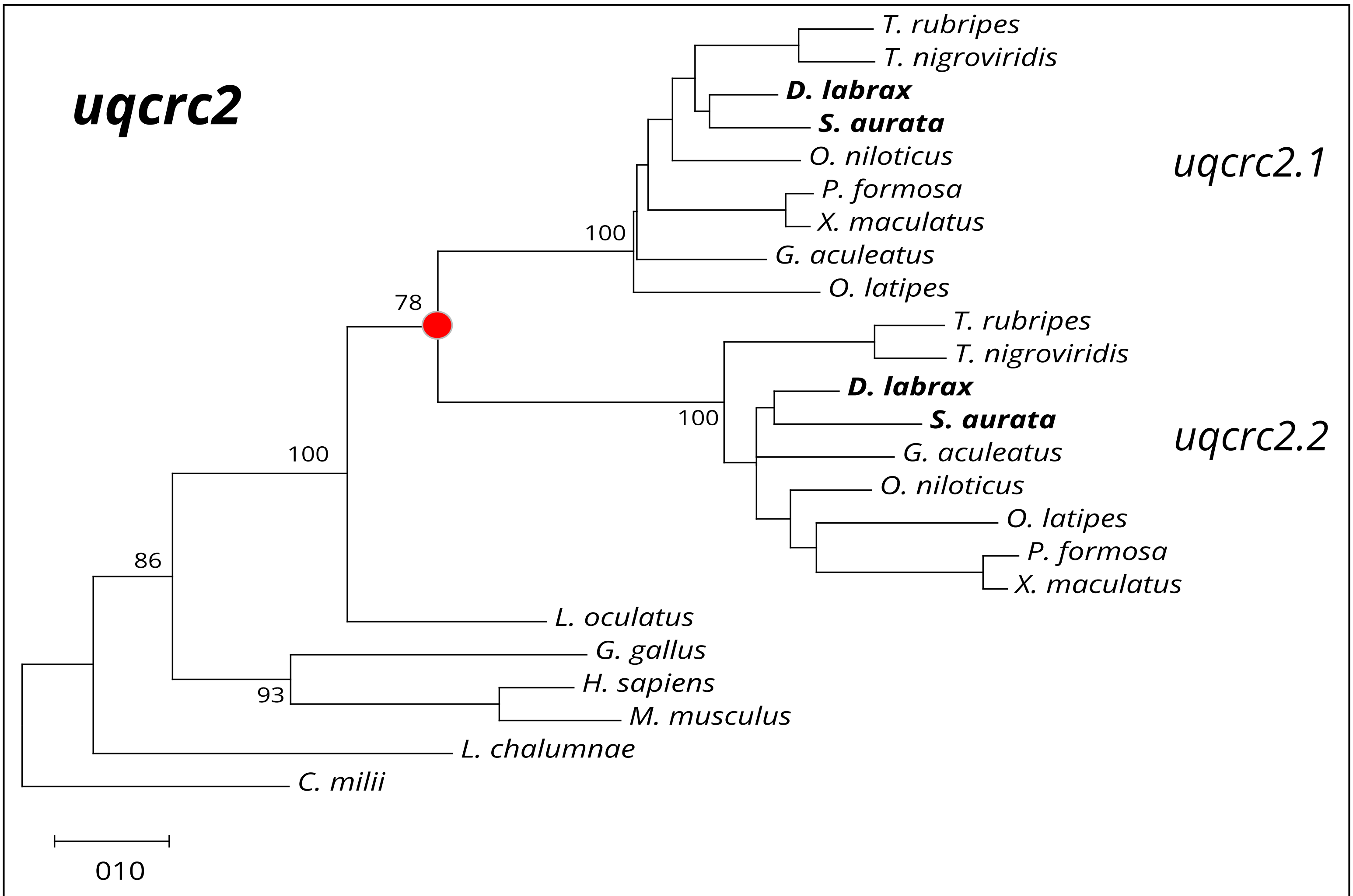

***Figure S6***. *Phylogenetic tree for uqcrc2 gene family*

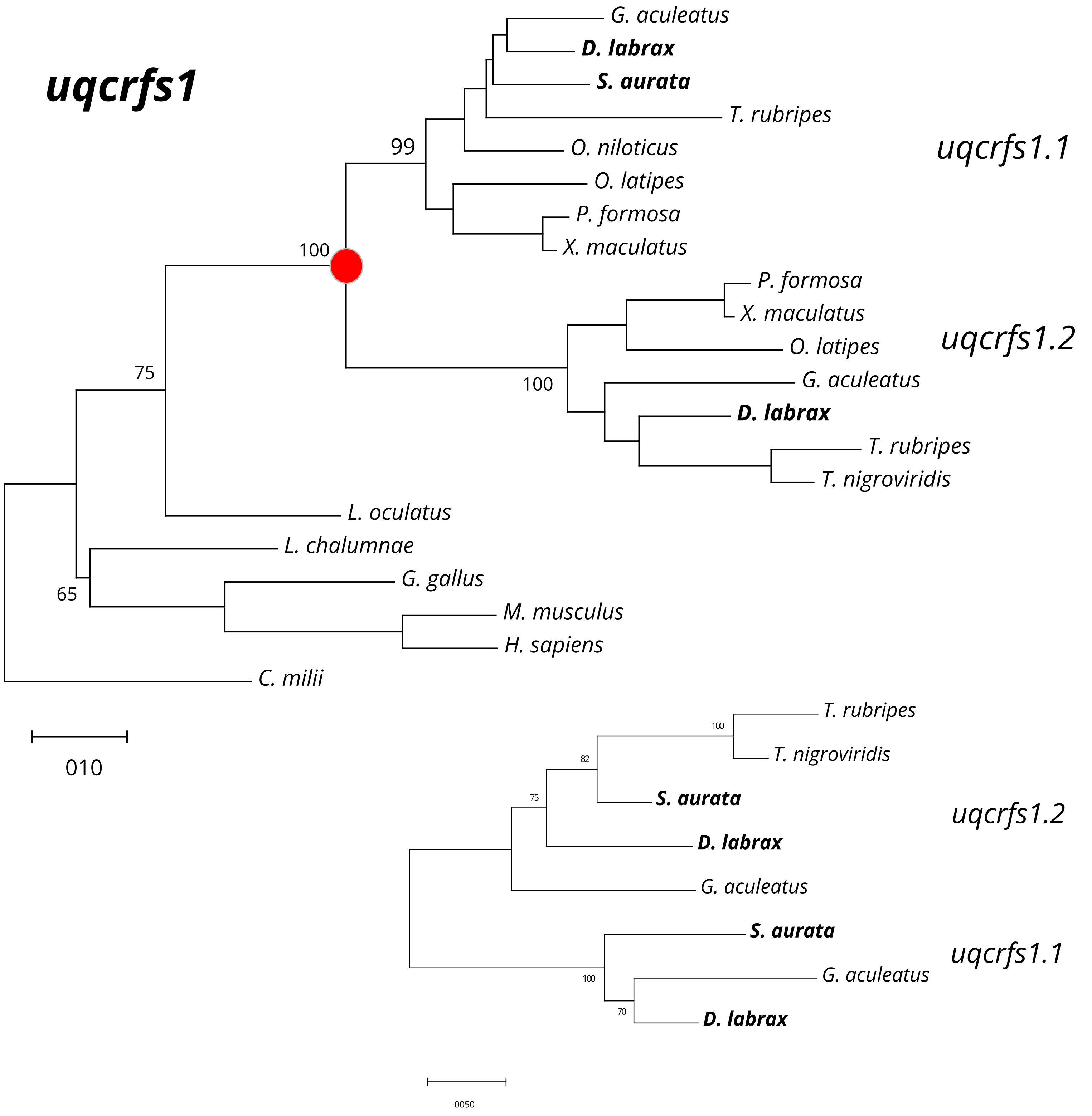

***Figure S7***. *Phylogenetic tree for uqcrfs1 gene family*

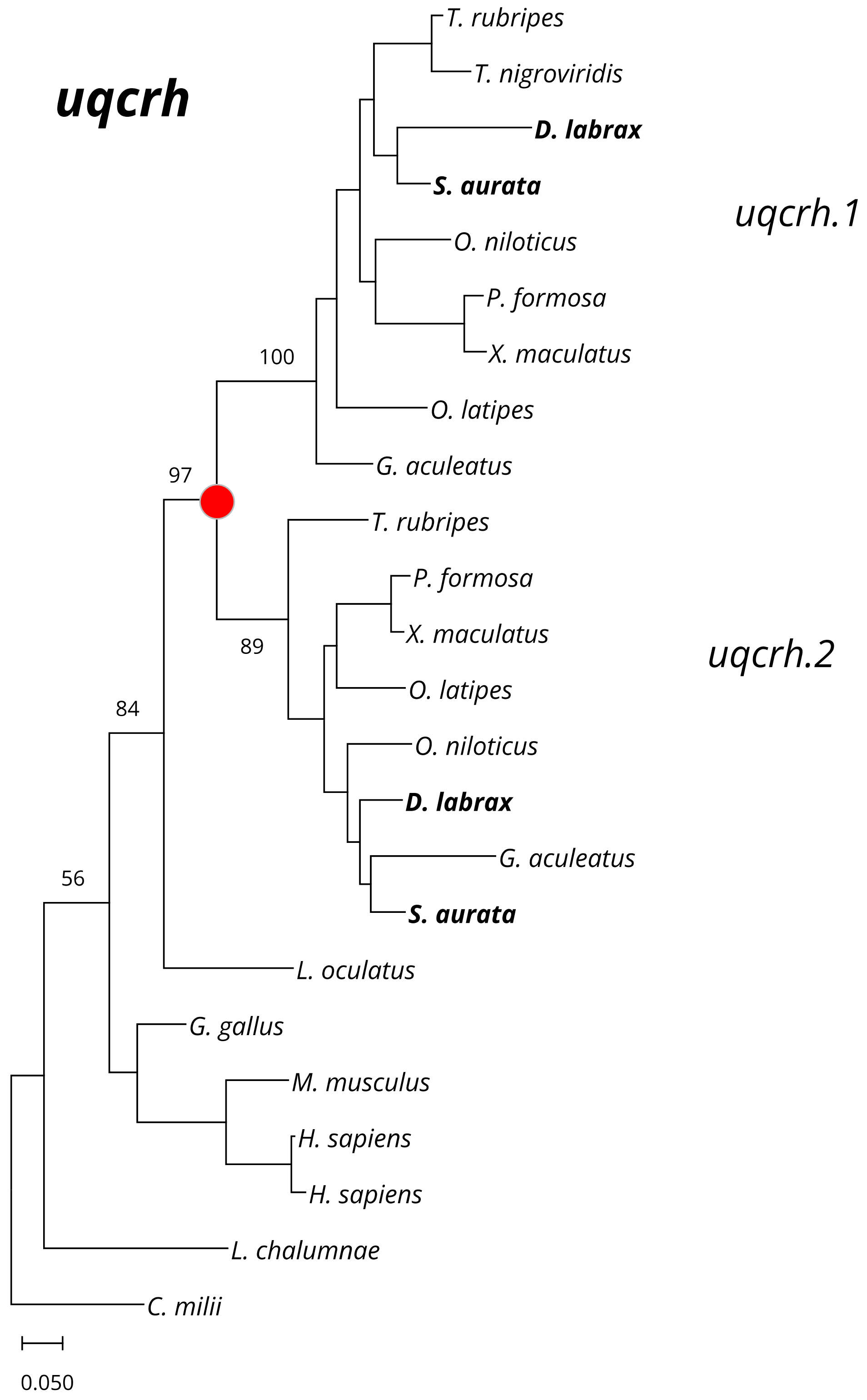

***Figure S8.*** *Phylogenetic tree for uqcrh gene family*

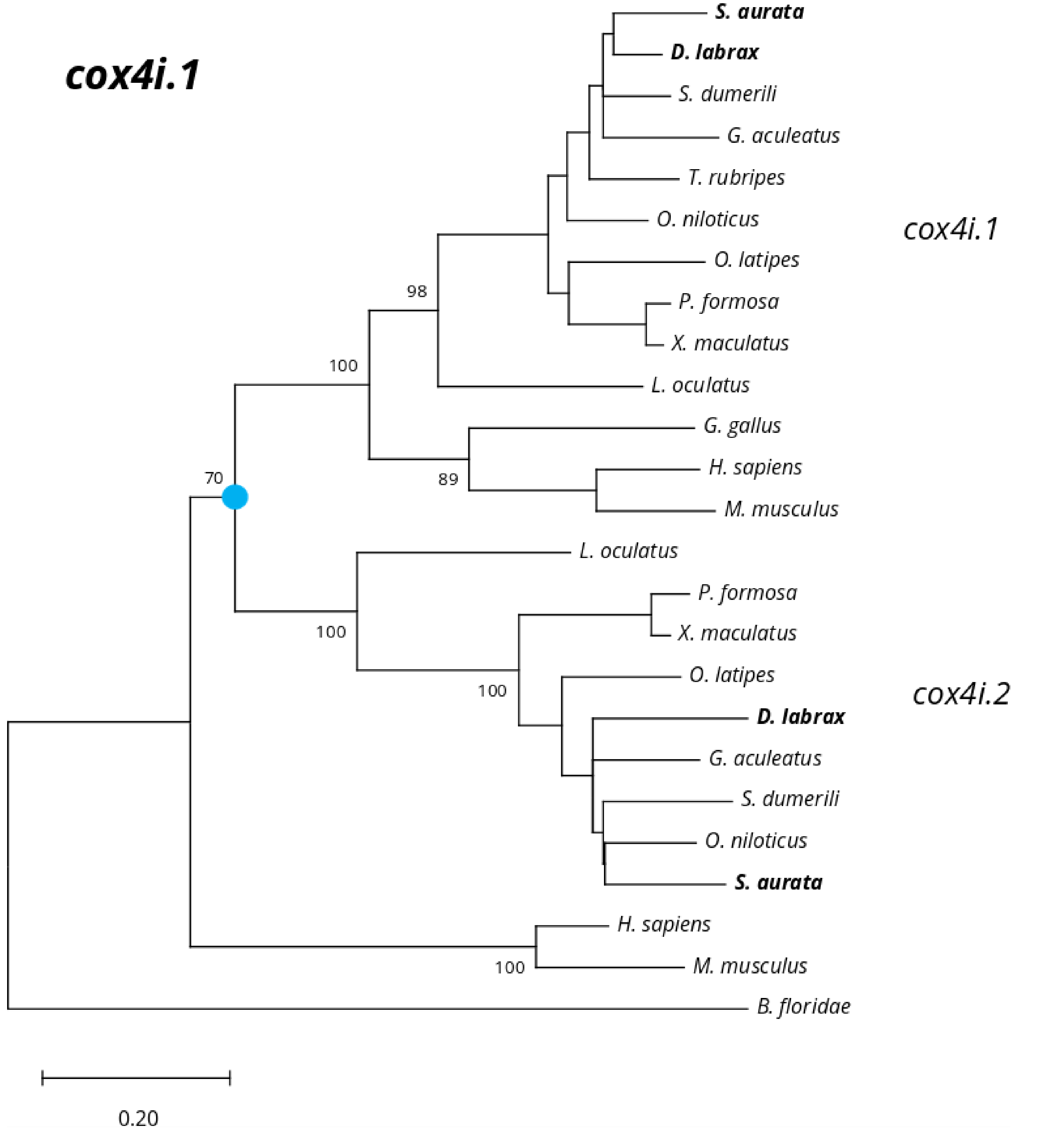

***Figure S9*** *Phylogenetic tree for cox4i.1 gene family*

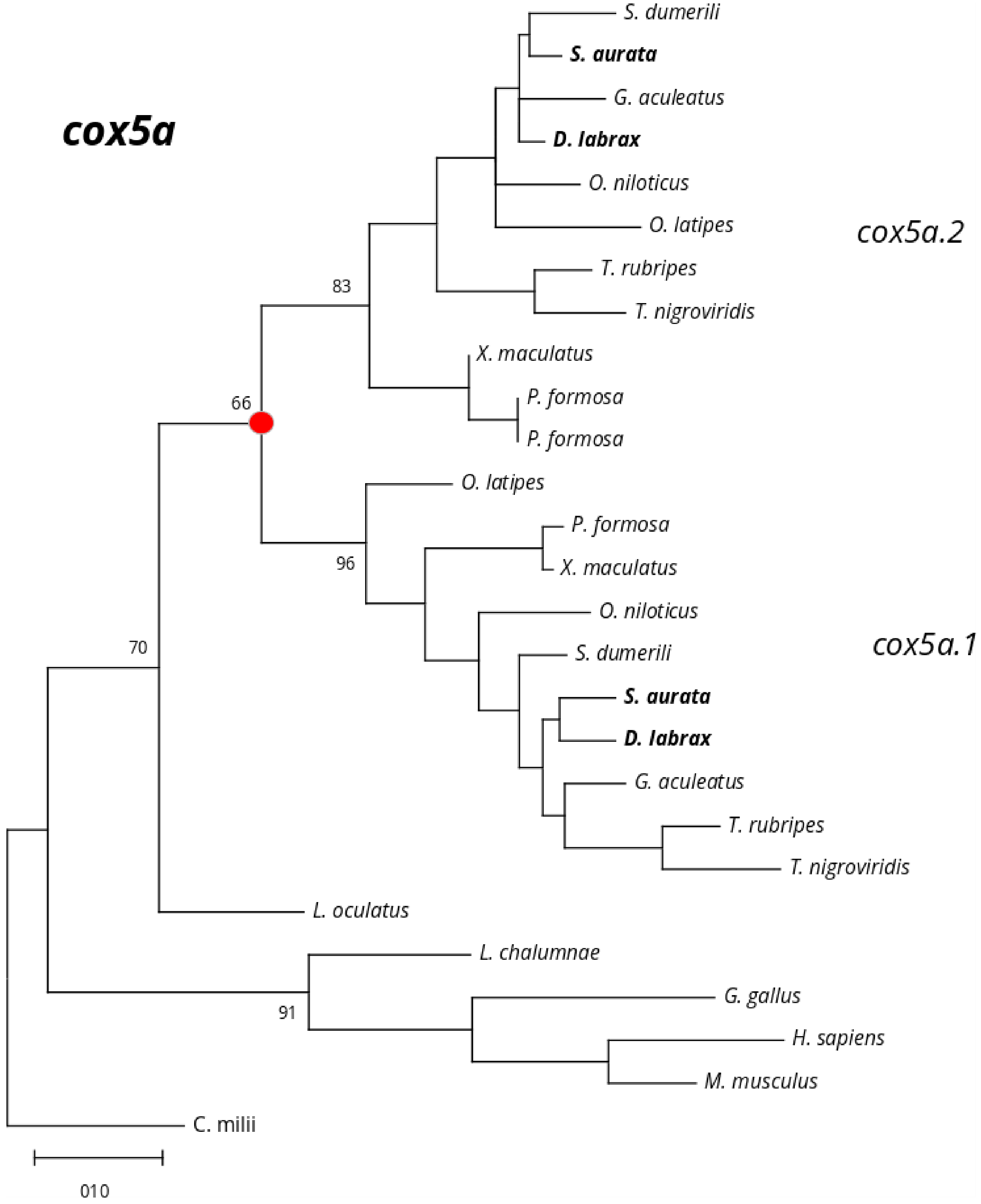

***Figure S10***. *Phylogenetic tree for cox5a gene family*

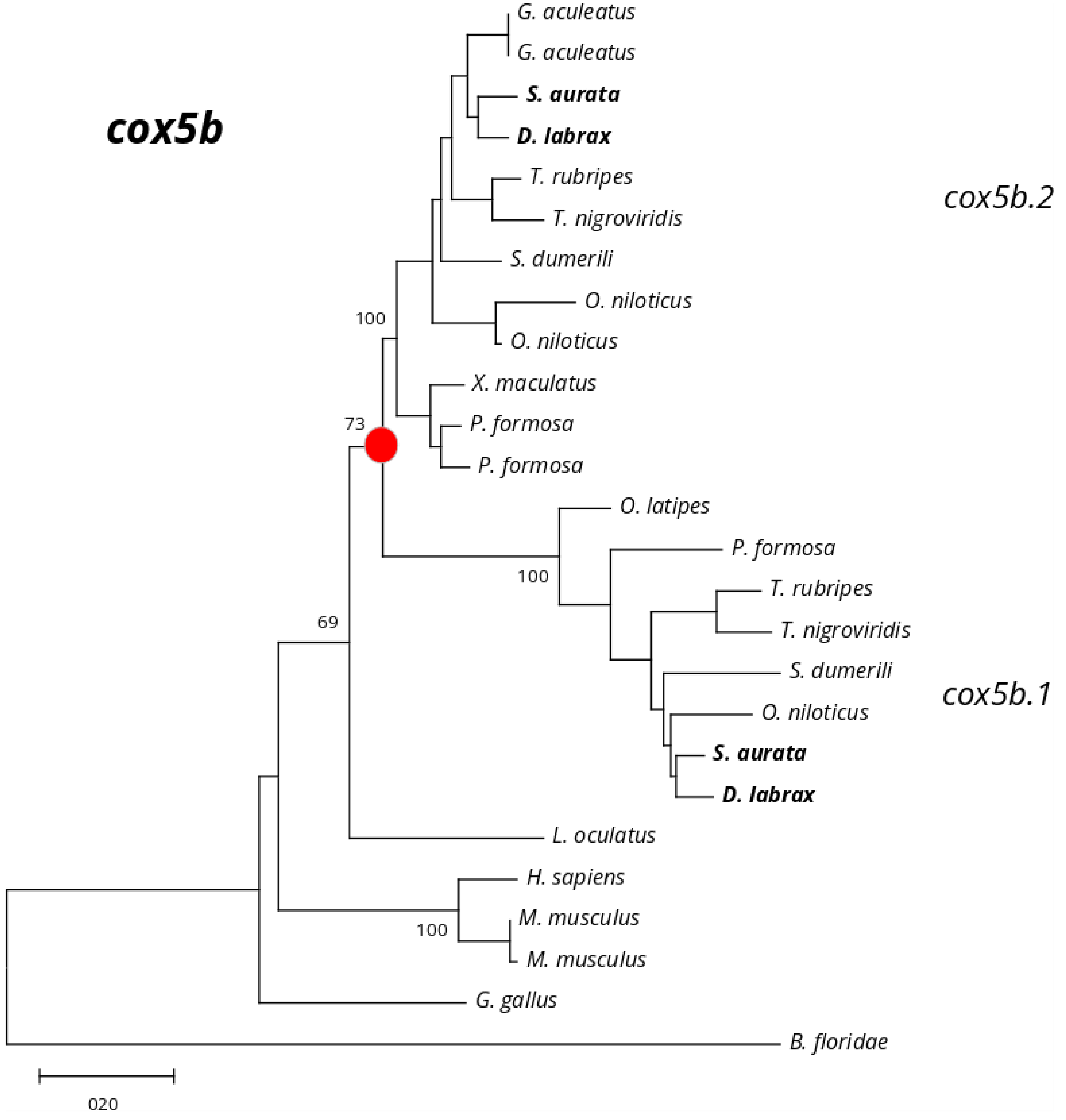

***Figure S11***. *Phylogenetic tree for cox5b gene family*

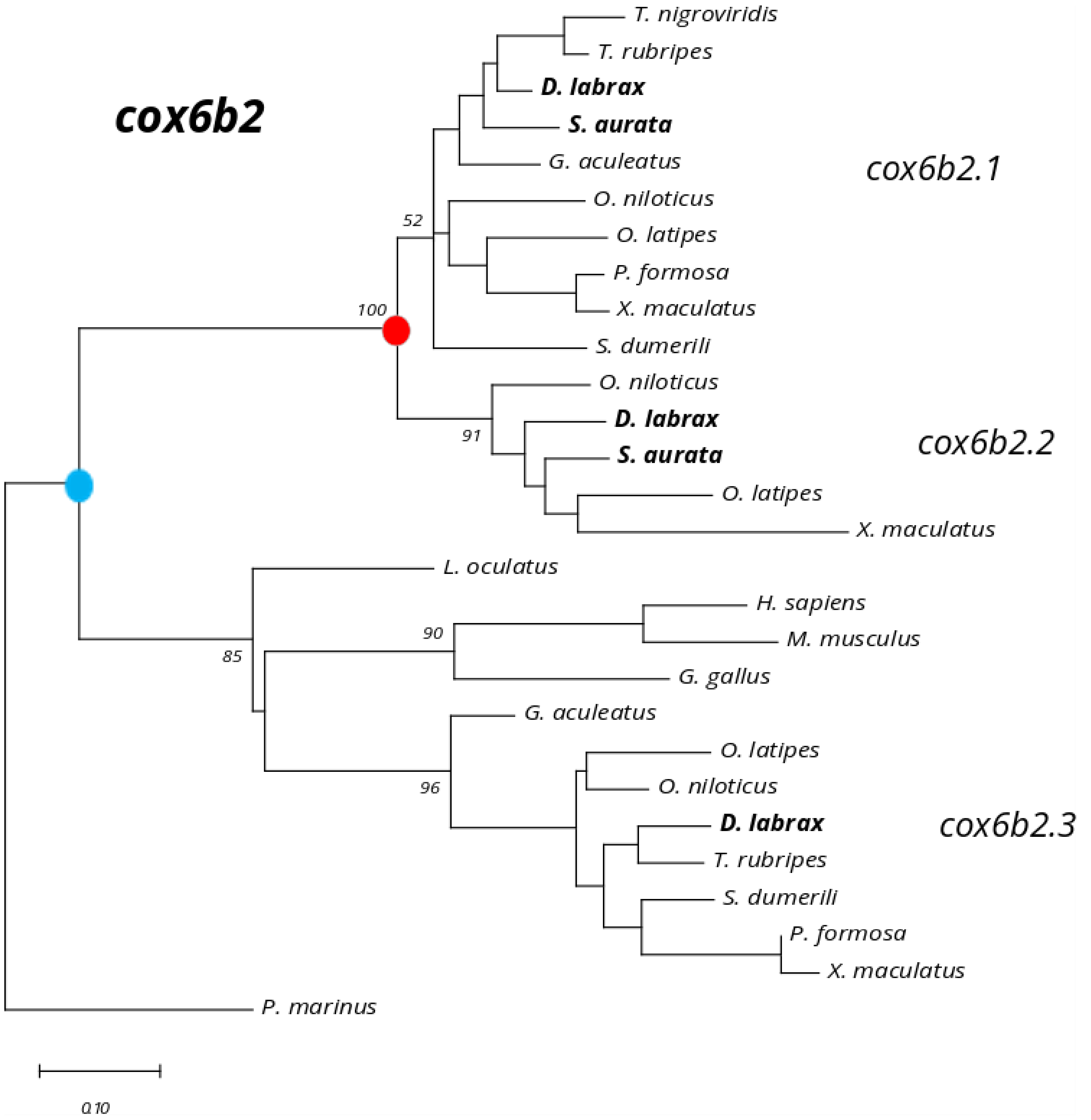

***Figure S12.*** *Phylogenetic tree for cox6b2 gene family*

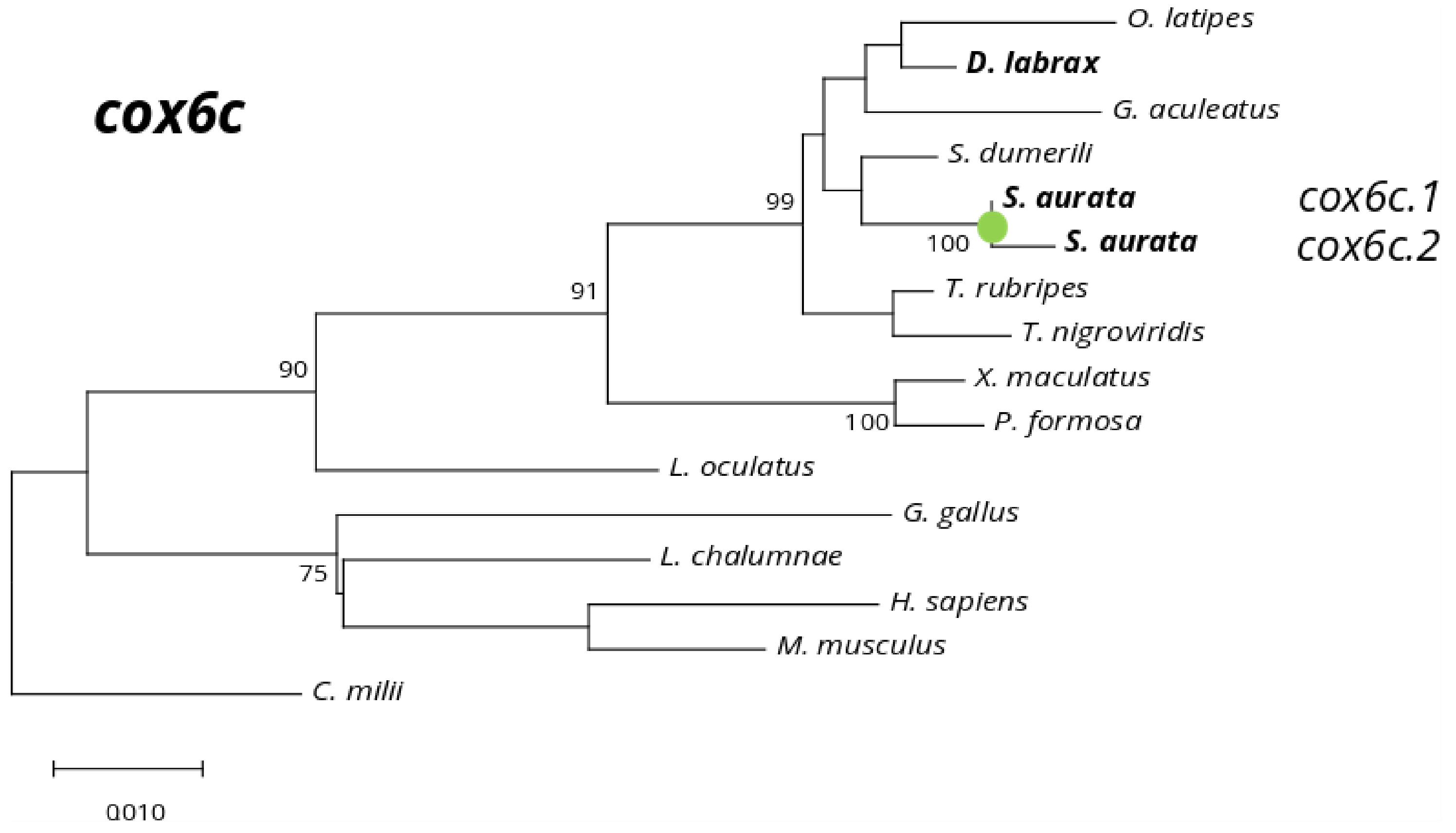

***Figure S13***. *Phylogenetic tree for cox6c gene family*

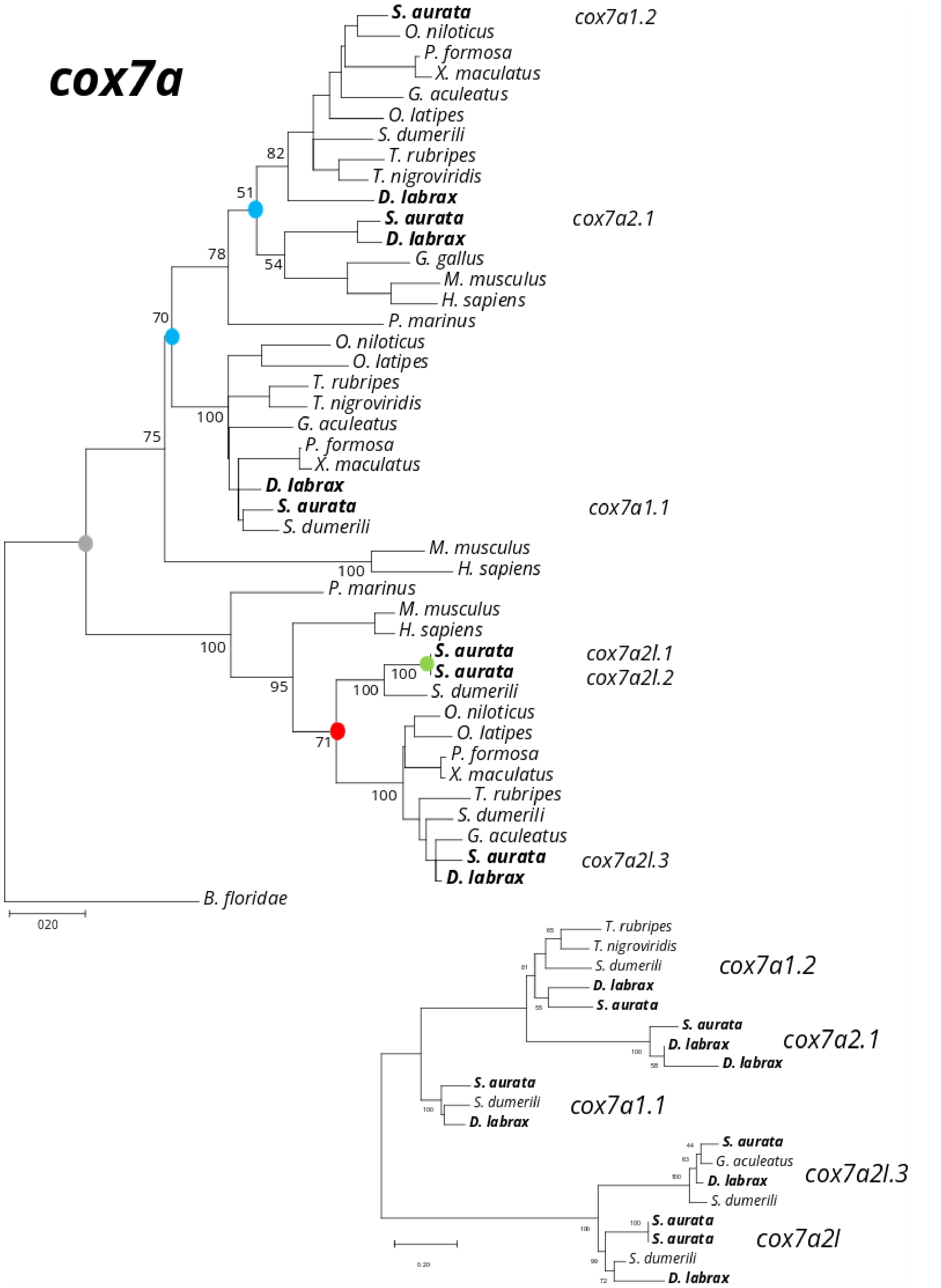

***Figure S14***. *Phylogenetic tree for cox7a gene family*

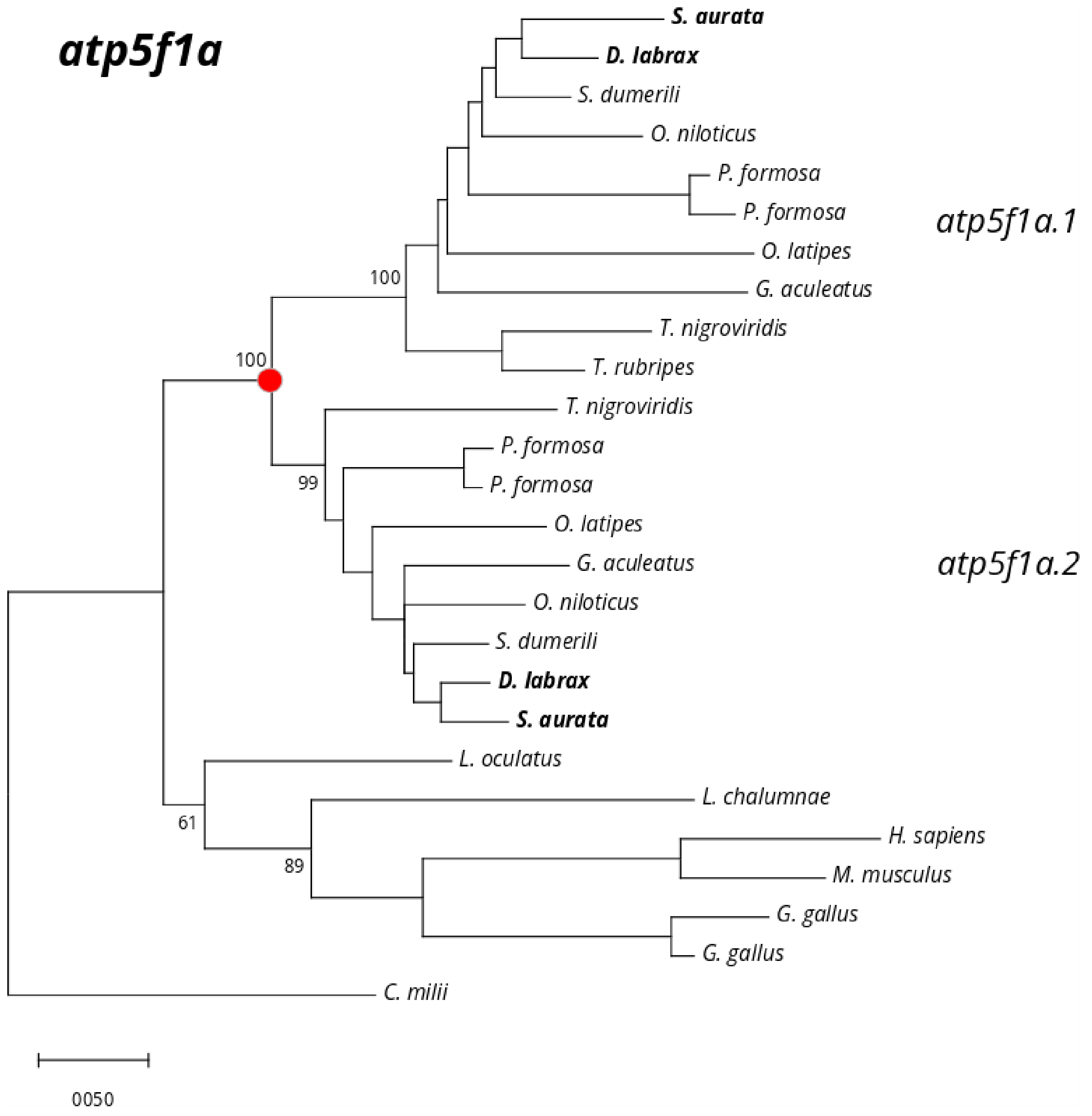

***Figure S15***. *Phylogenetic tree for atp5f1a gene family.*

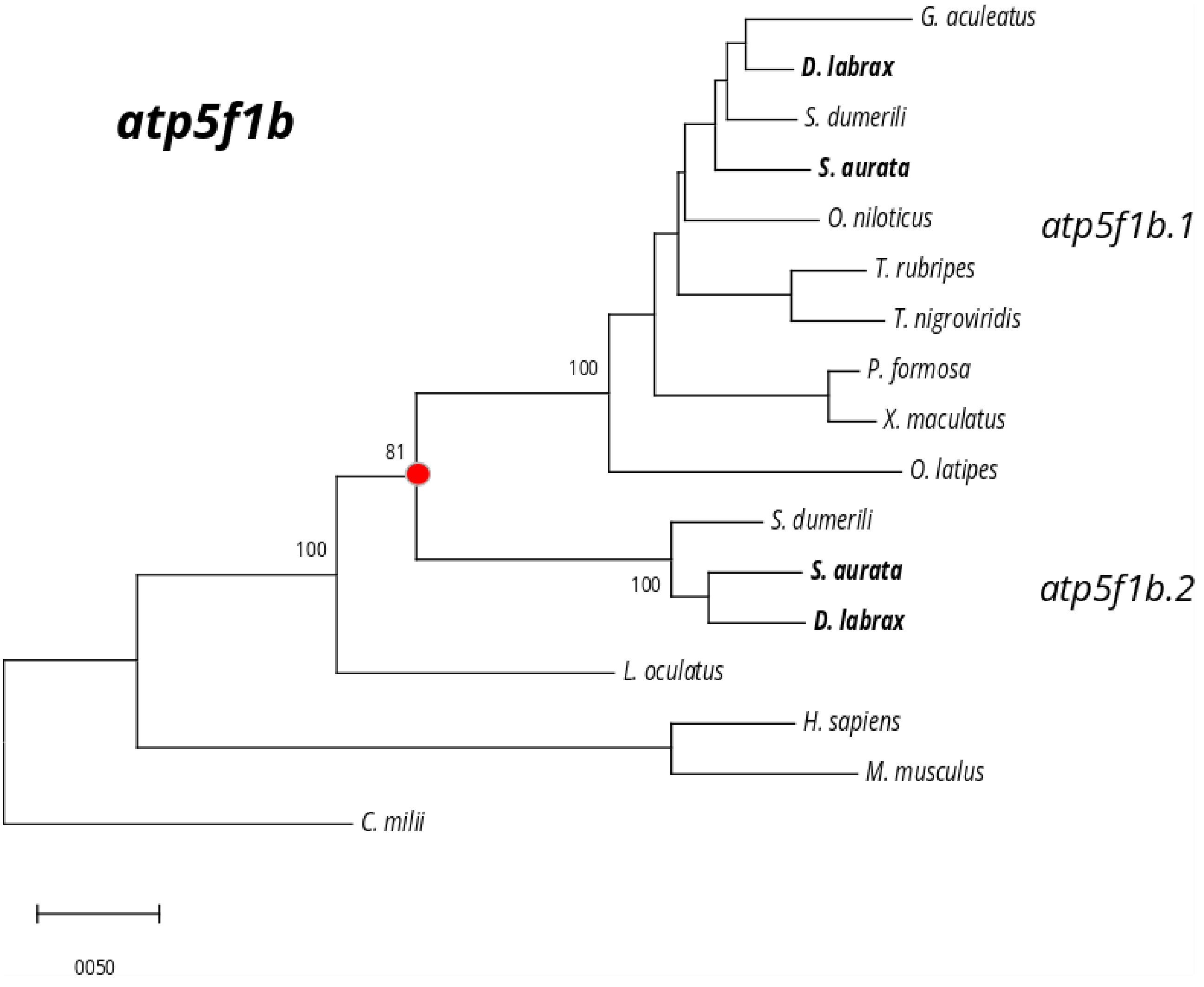

***Figure S16.*** *Phylogenetic tree for atp5f1b gene family*

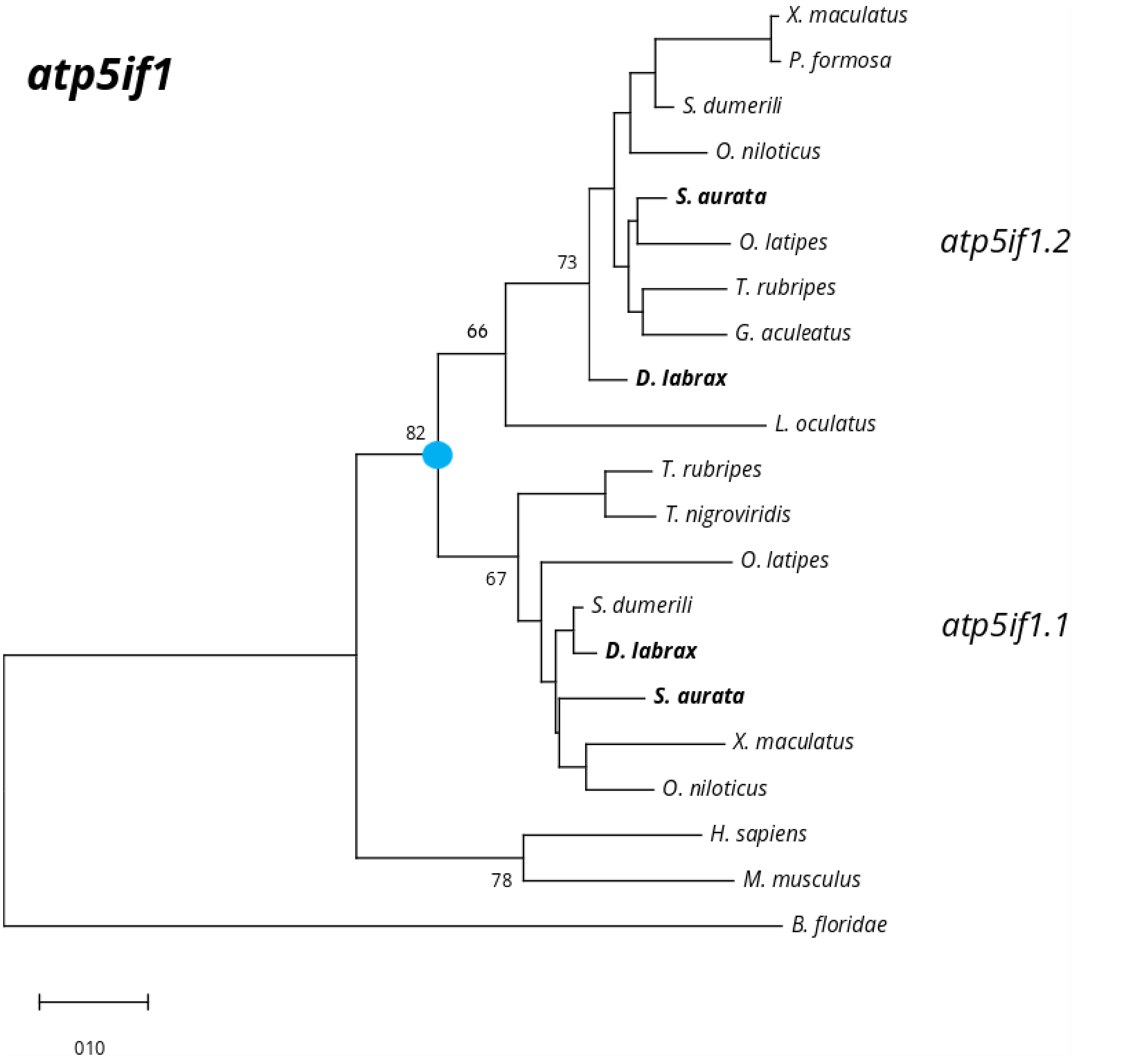

***Figure S17***. *Phylogenetic tree for atp5if1 gene family*

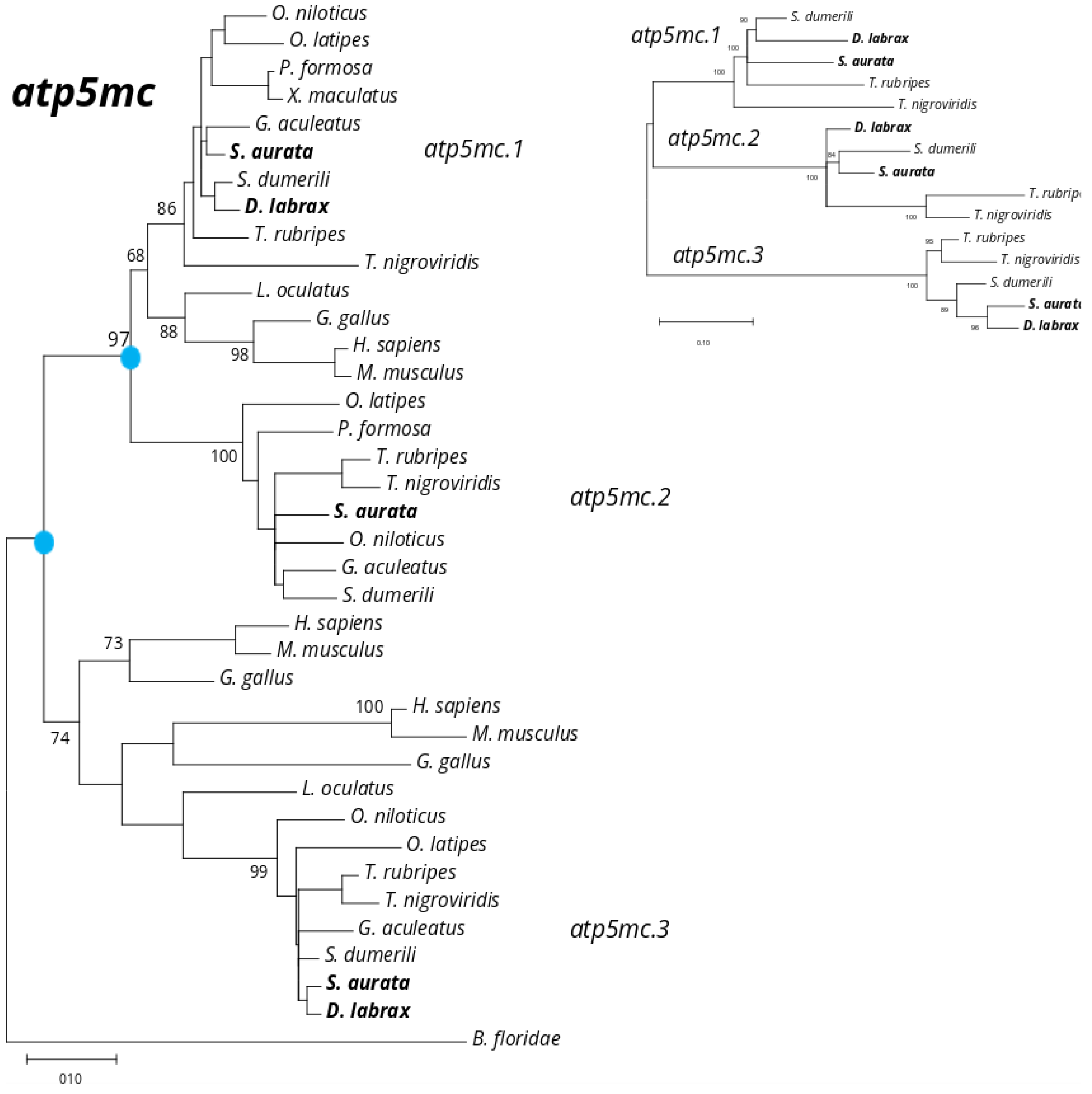

***Figure S18.*** *Phylogenetic tree for atp5mc gene family*

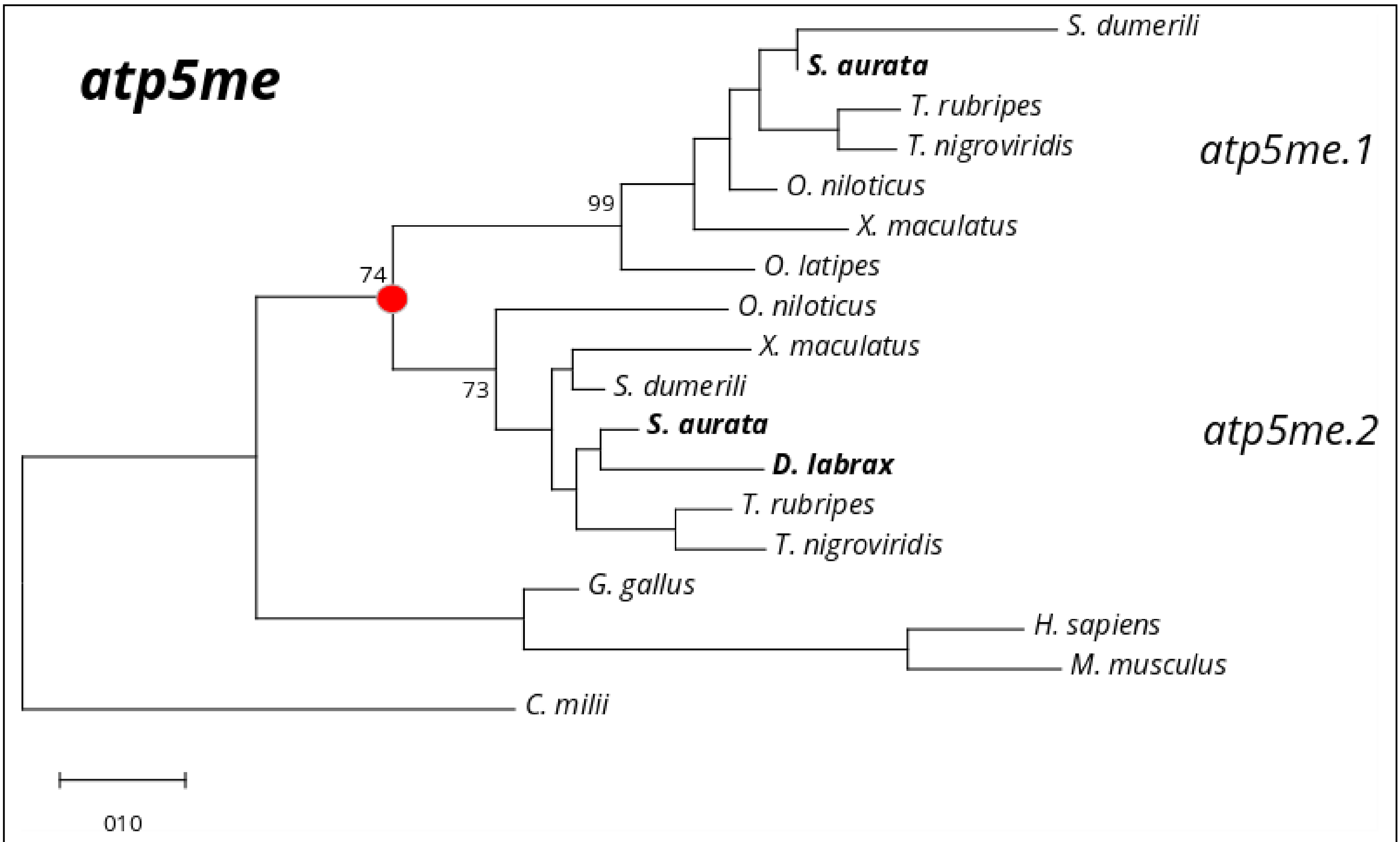

***Figure S19***. *Phylogenetic tree for atp5me gene family*

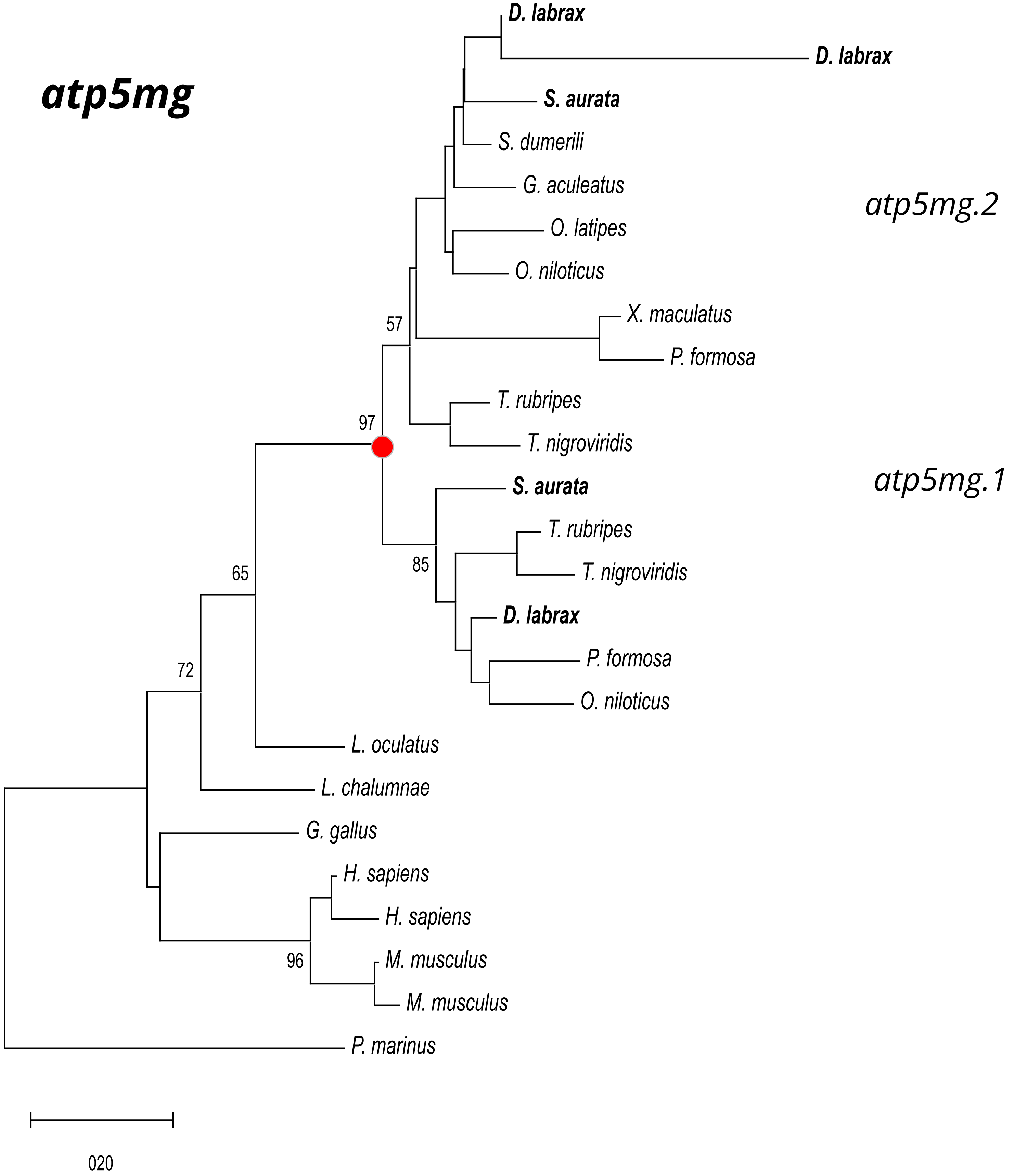

***Figure S20.*** *Phylogenetic tree for atp5mg gene family*

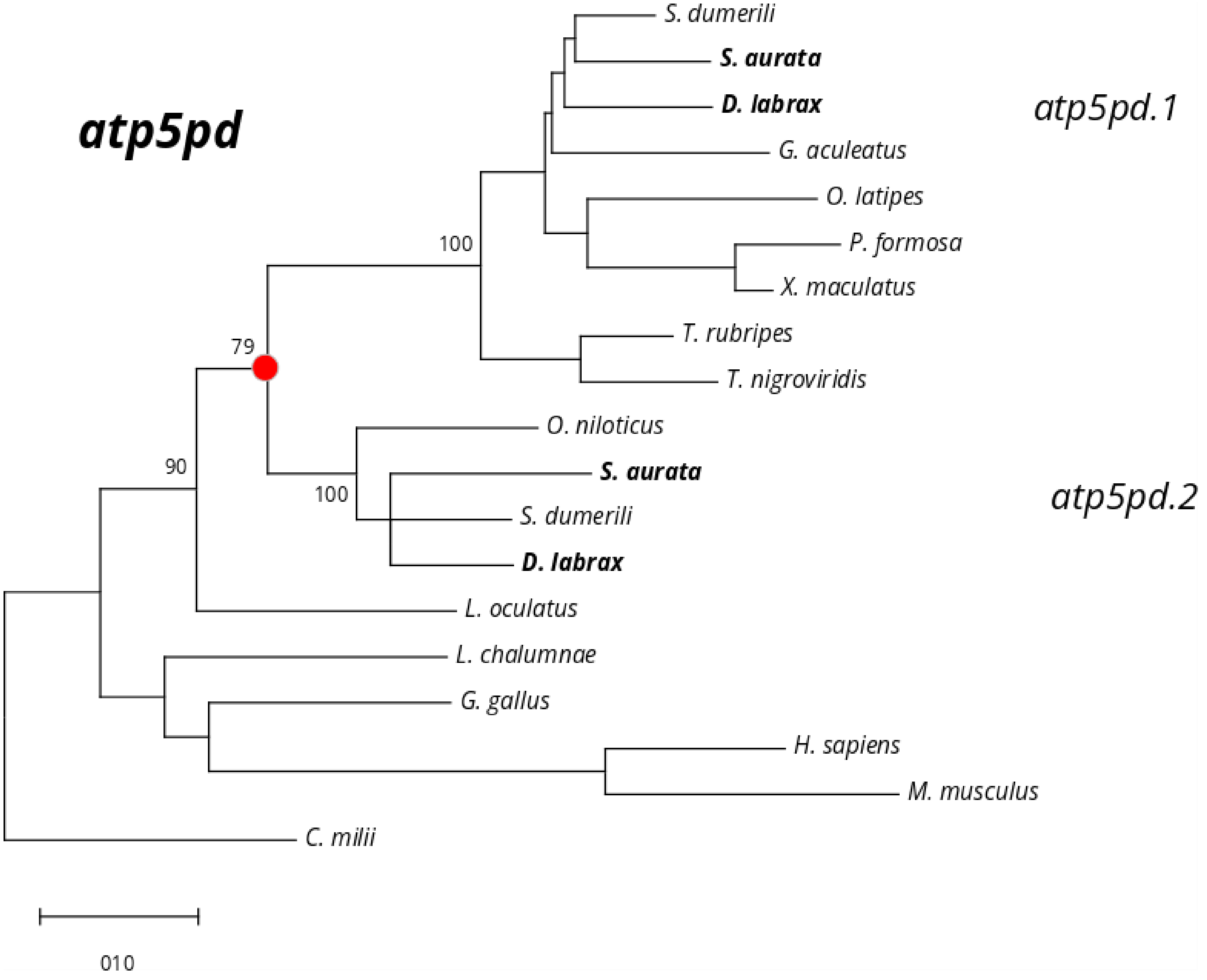

***Figure S21.*** *Phylogenetic tree for atp5pd gene family*

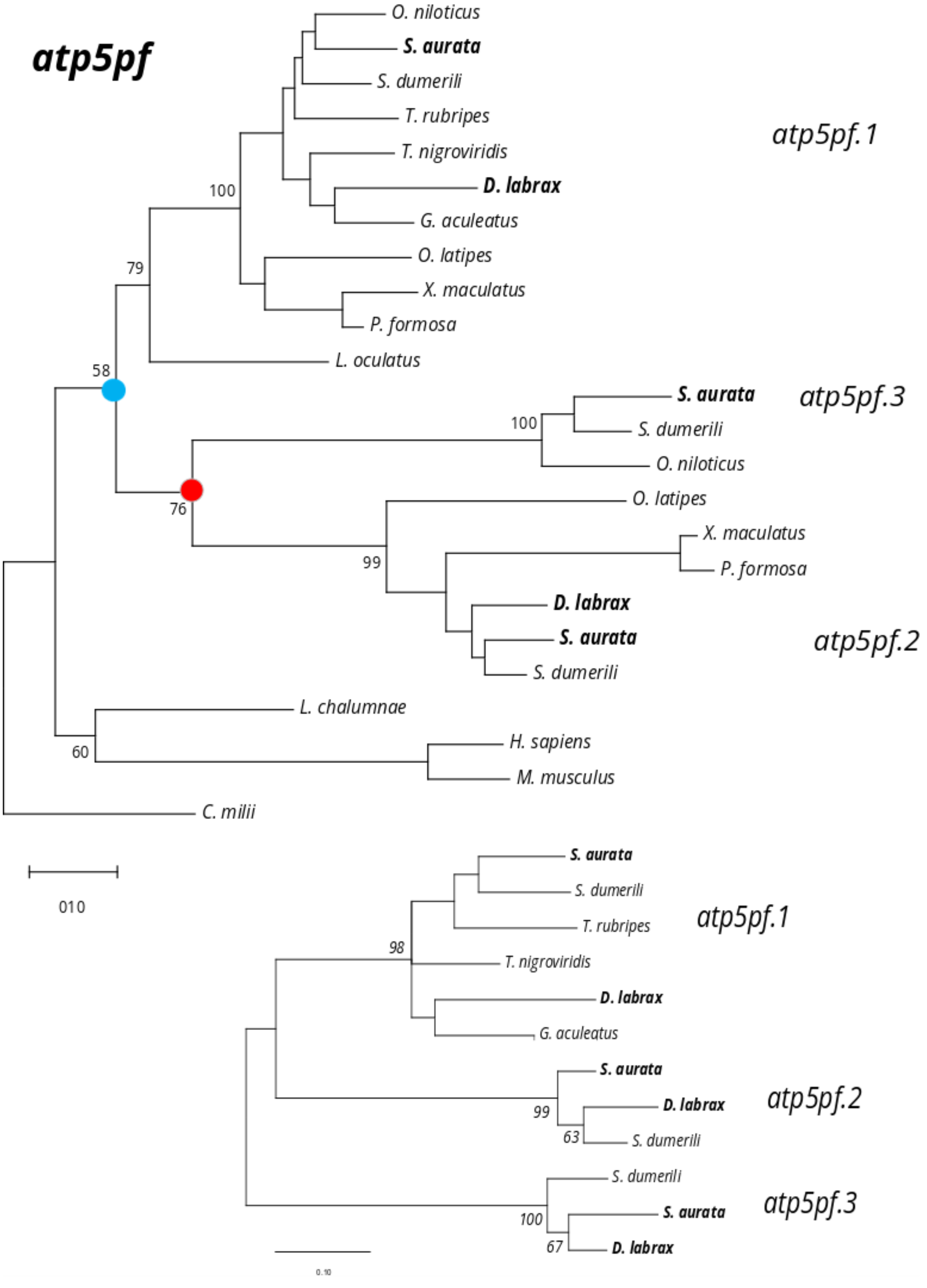

***Figure S22.*** *Phylogenetic tree for atp5pf gene family*

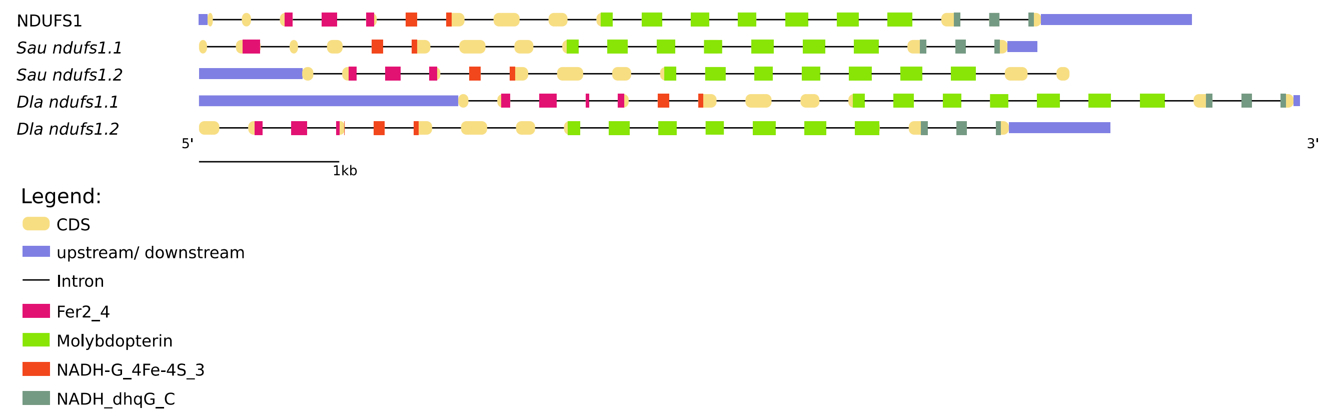

***Figure S23***. *Gene structure and protein domains for ndufs1 gene family in gilthead seabream (Sau), European seabass (Dla) and human (capital letters).*

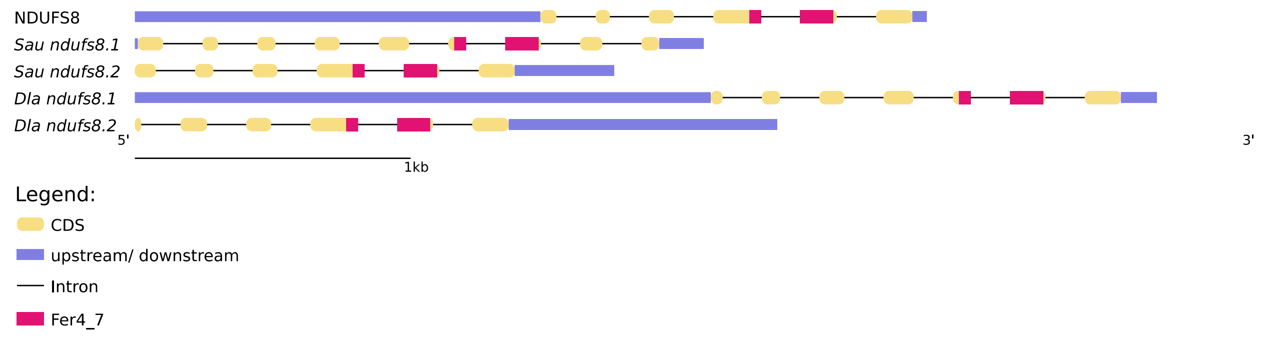

***Figure S24***. *Gene structure and protein domains for ndufs8 gene family in gilthead seabream (Sau), European seabass (Dla) and human (capital letters)*

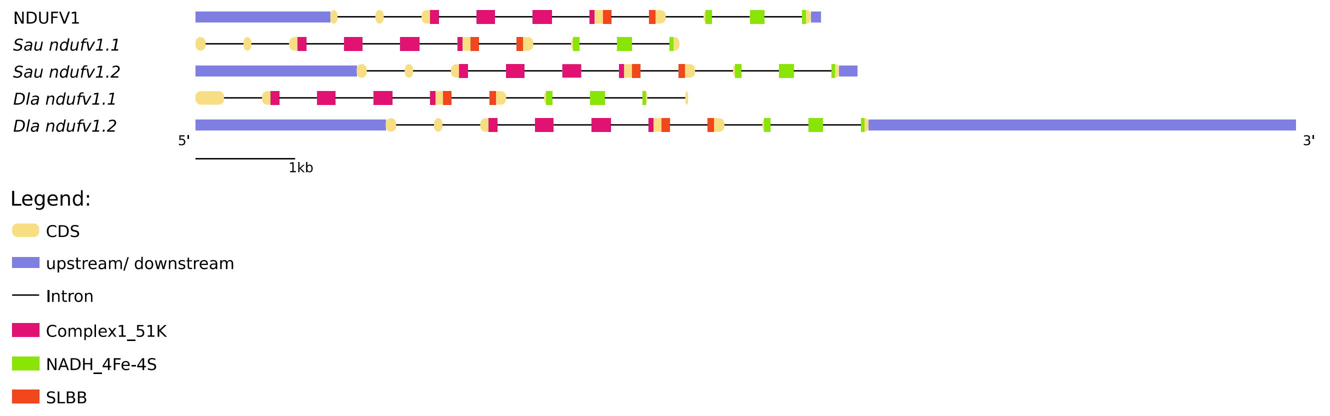

***Figure S25.*** *Gene structure and protein domains for ndufv1 gene family in gilthead seabream (Sau), European seabass (Dla) and human (capital letters).*

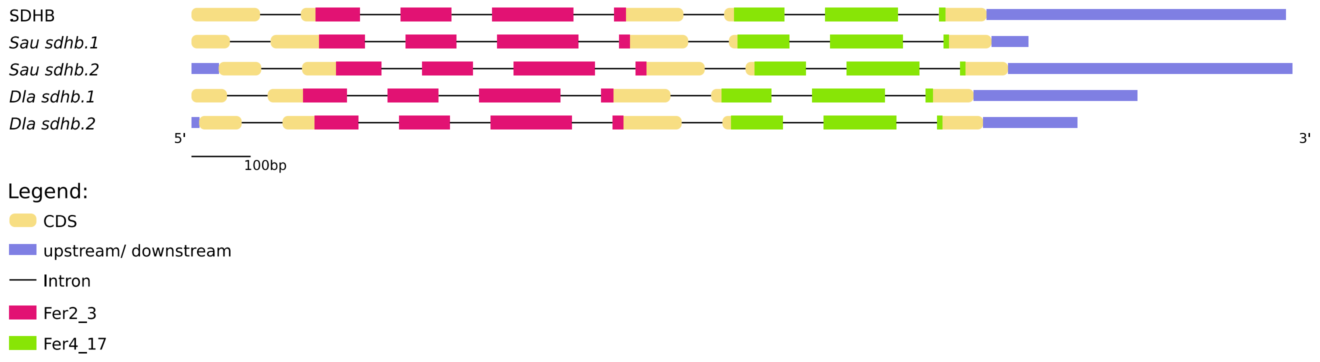

***Figure S26.*** *Gene structure and protein domains for sdhb gene family in gilthead seabream (Sau), European seabass (Dla) and human (capital letters).*

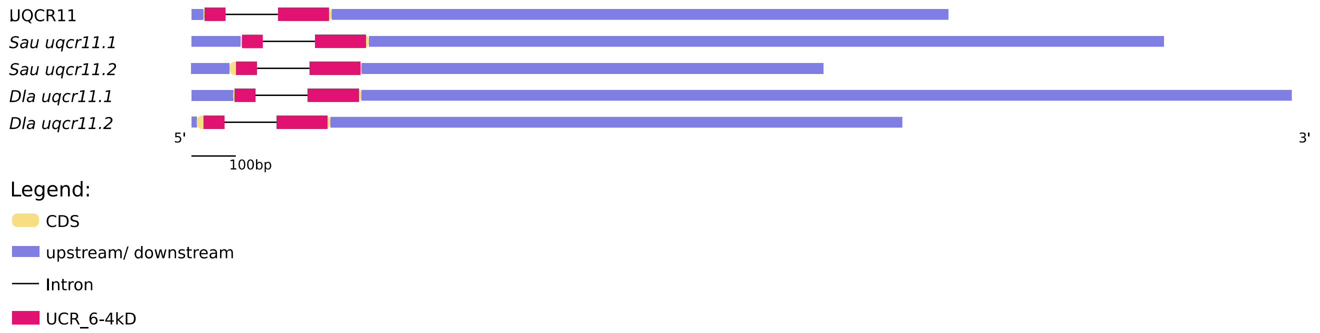

***Figure S27.*** *Gene structure and protein domains for uqcr11 gene family in gilthead seabream (Sau), European seabass (Dla) and human (capital letters).*

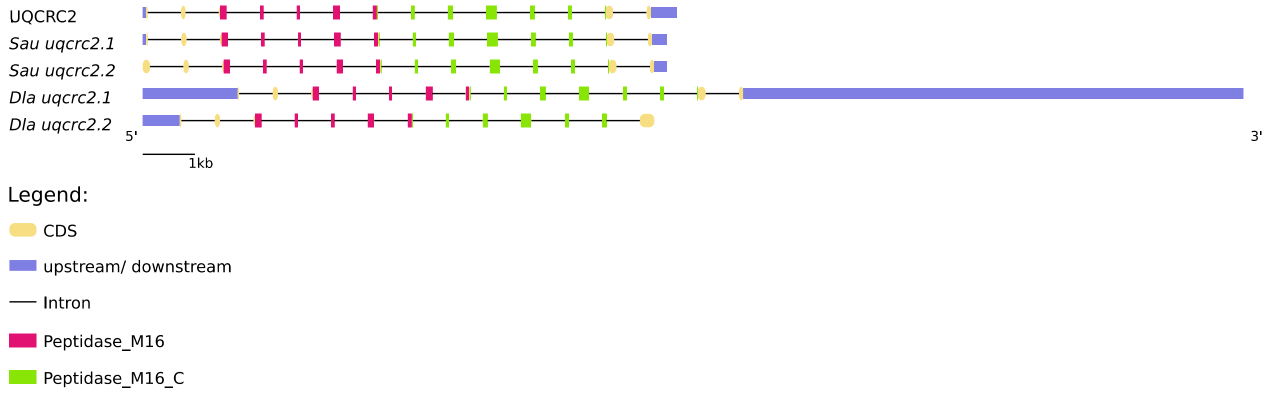

***Figure S28.*** *Gene structure and protein domains for uqcrc2 gene family in gilthead seabream (Sau), European seabass (Dla) and human (capital letters).*

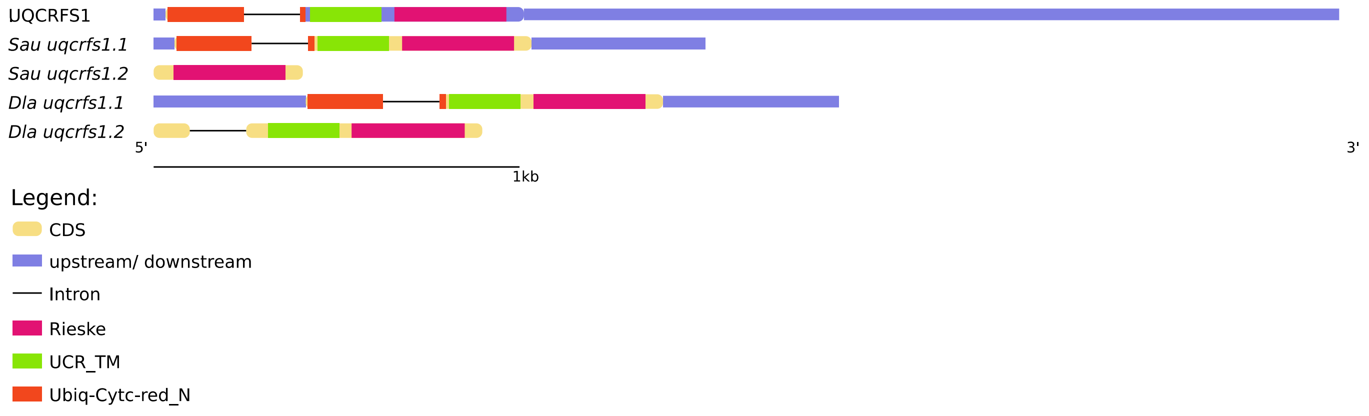

***Figure S29.*** *Gene structure and protein domains for uqcrfs1 gene family in gilthead seabream (Sau), European seabass (Dla) and human (capital letters).*

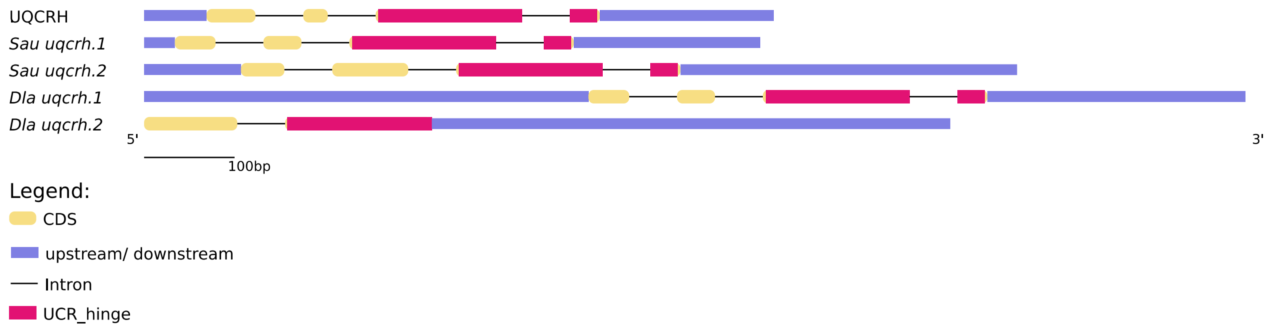

***Figure S30.*** *Gene structure and protein domains for uqcrh gene family in gilthead seabream (Sau), European seabass (Dla) and human (capital letters).*

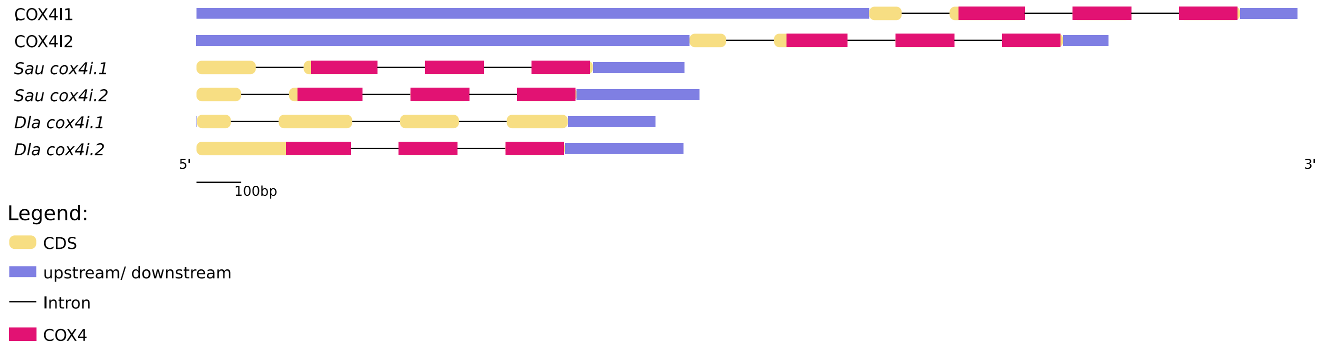

***Figure S31.*** *Gene structure and protein domains for cox4i gene family in gilthead seabream (Sau), European seabass (Dla) and human (capital letters).*

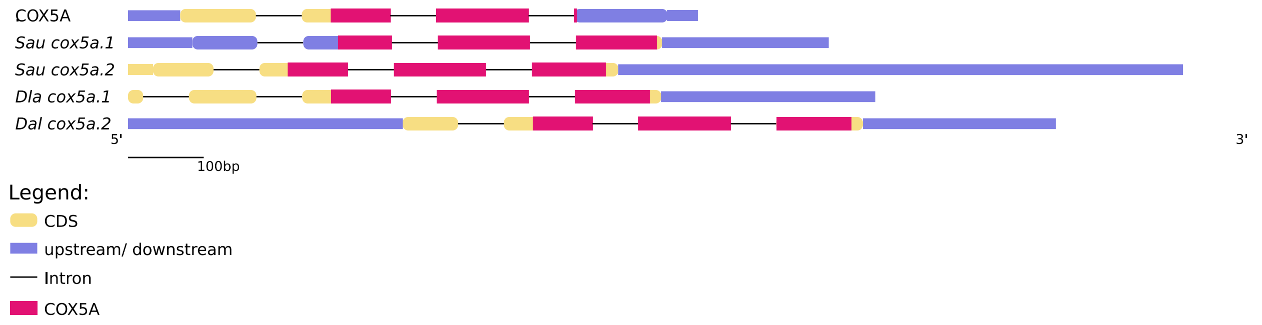

***Figure S32.*** *Gene structure and protein domains for cox5a gene family in gilthead seabream (Sau), European seabass (Dla) and human (capital letters).*

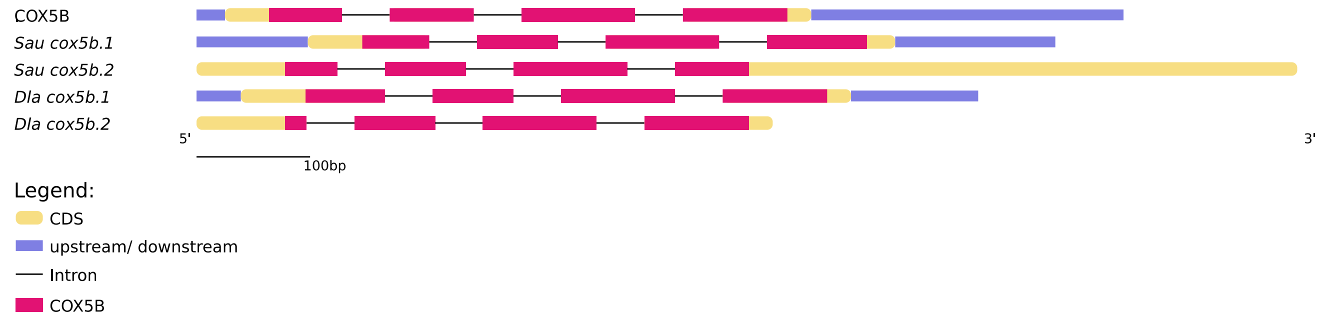

***Figure S33.*** *Gene structure and protein domains for cox5b gene family in gilthead seabream (Sau), European seabass (Dla) and human (capital letters).*

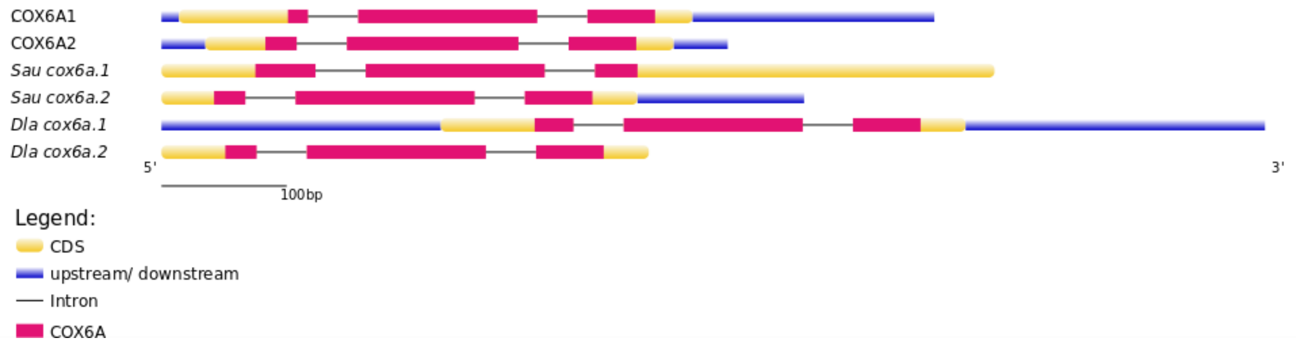

***Figure S34.*** *Gene structure and protein domains for cox6a gene family in gilthead seabream (Sau), European seabass (Dla) and human (capital letters).*

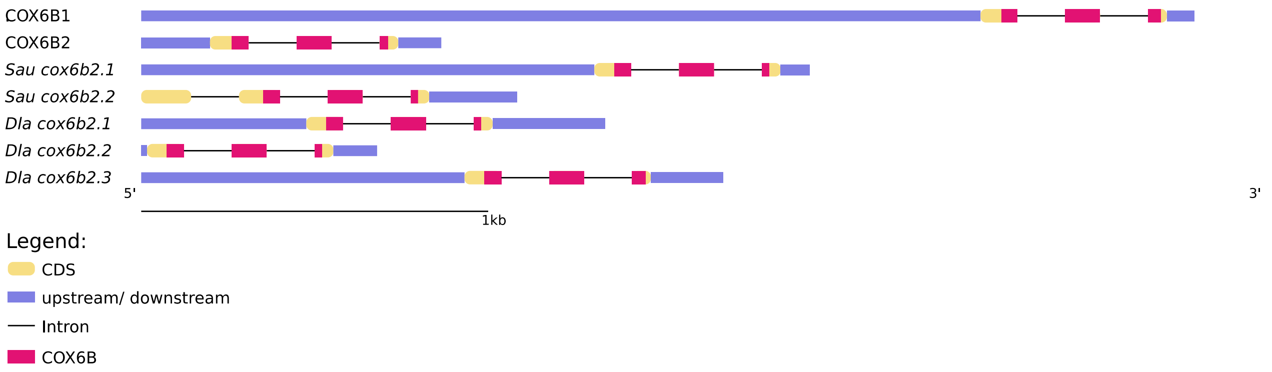

***Figure S35.*** *Gene structure and protein domains for cox6b gene family in gilthead seabream (Sau), European seabass (Dla) and human (capital letters).*

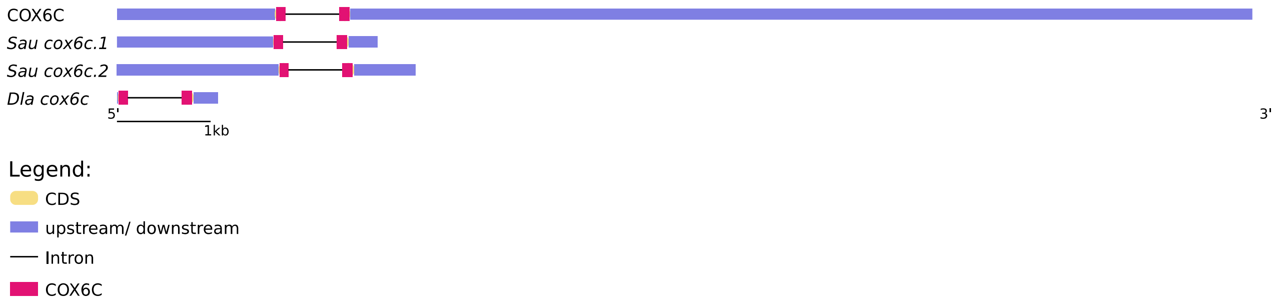

***Figure S36.*** *Gene structure and protein domains for cox6c gene family in gilthead seabream (Sau), European seabass (Dla) and human (capital letters).*

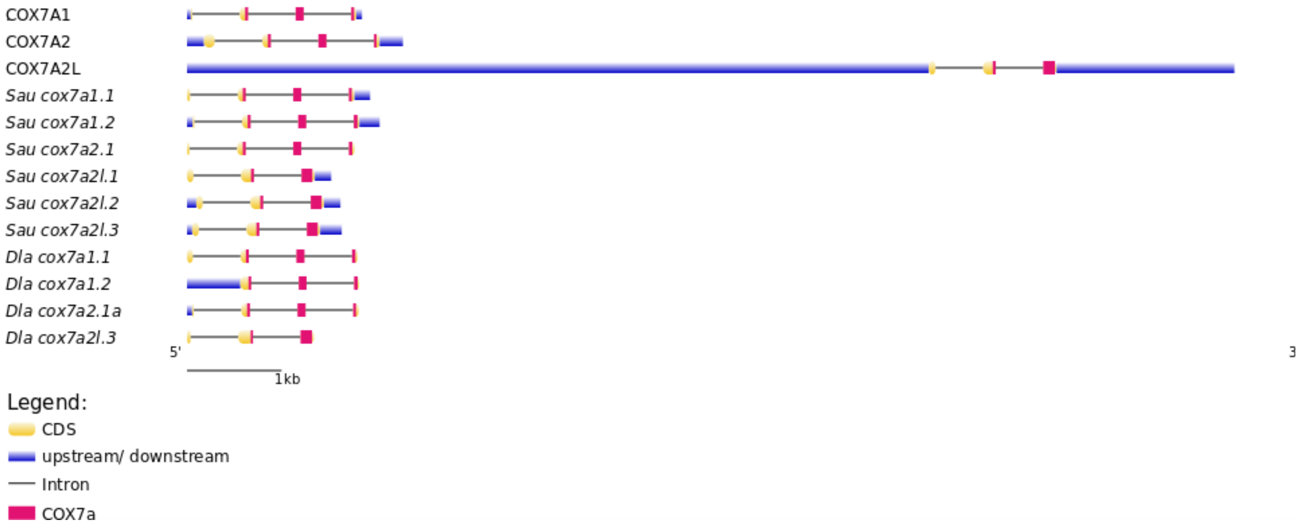

***Figure S37.*** *Gene structure and protein domains for cox7a gene family in gilthead seabream (Sau), European seabass (Dla) and human (capital letters).*

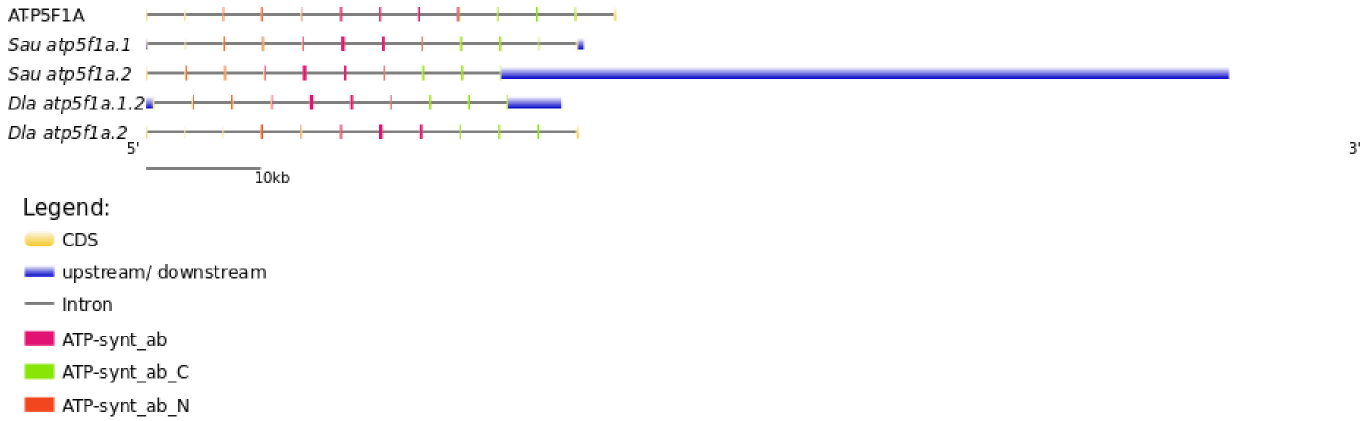

***Figure S38.*** *Gene structure and protein domains for atp5f1a gene family in gilthead seabream (Sau), European seabass (Dla) and human (capital letters).*

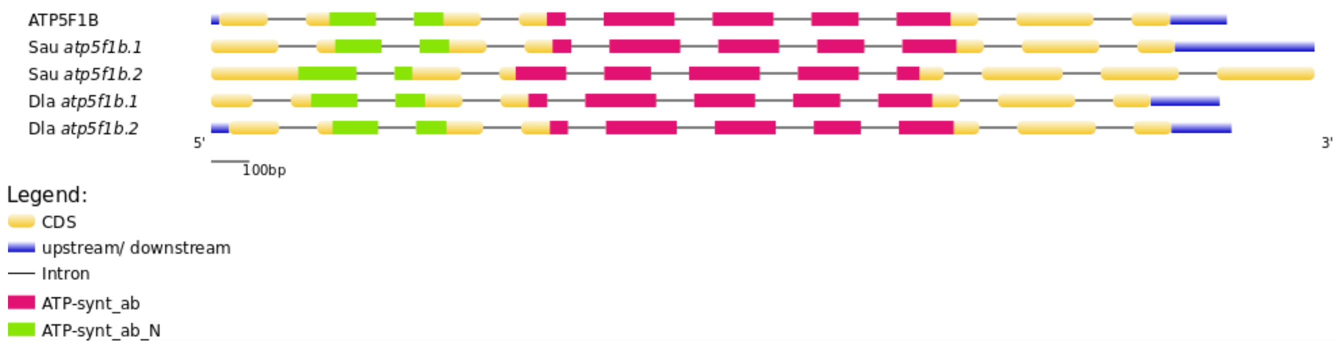

***Figure S39.*** *Gene structure and protein domains for atp5f1b gene family in gilthead seabream (Sau), European seabass (Dla) and human (capital letters).*

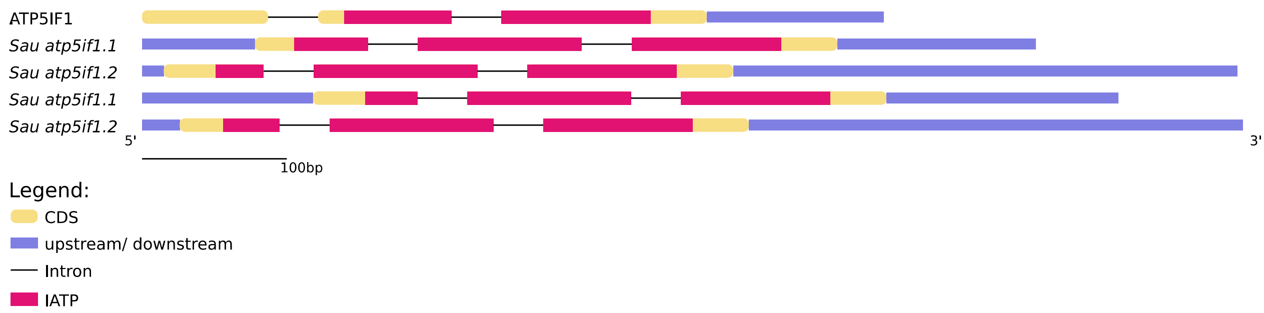

***Figure S40.*** *Gene structure and protein domains for atp5if1 gene family in gilthead seabream (Sau), European seabass (Dla) and human (capital letters).*

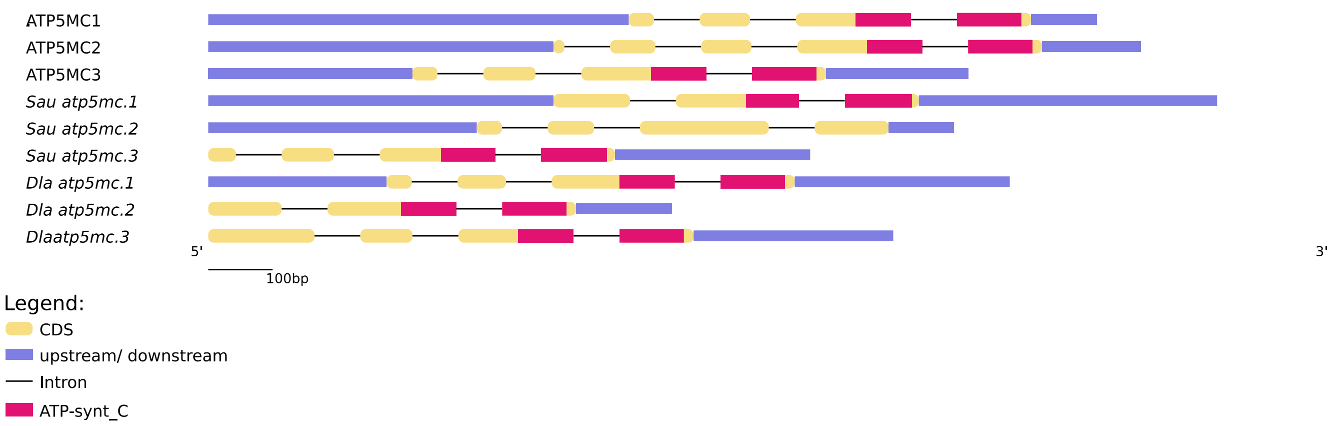

***Figure S41.*** *Gene structure and protein domains for atp5mc gene family in gilthead seabream (Sau), European seabass (Dla) and human (capital letters).*

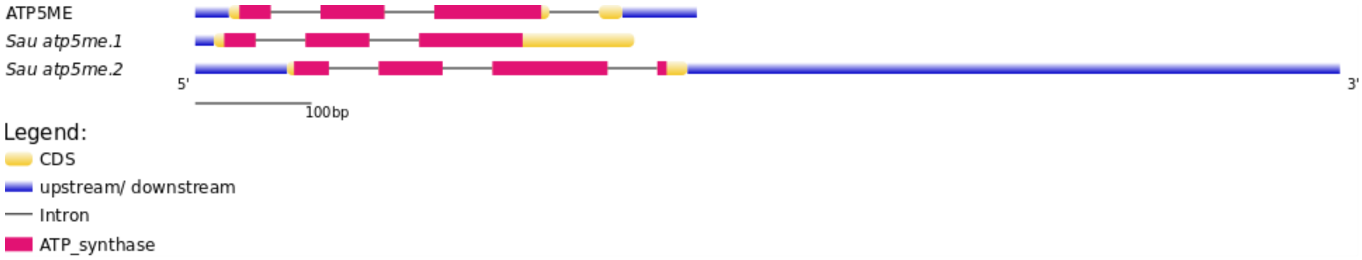

***Figure S42.*** *Gene structure and protein domains for atp5me gene family in gilthead seabream (Sau), European seabass (Dla) and human (capital letters).*

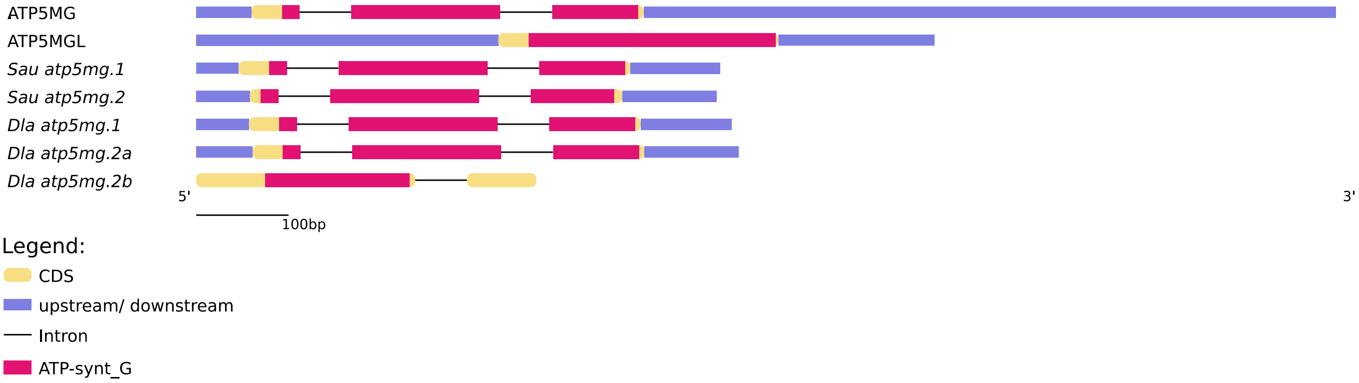

***Figure S43.*** *Gene structure and protein domains for atp5mg gene family in gilthead seabream (Sau), European seabass (Dla) and human (capital letters).*

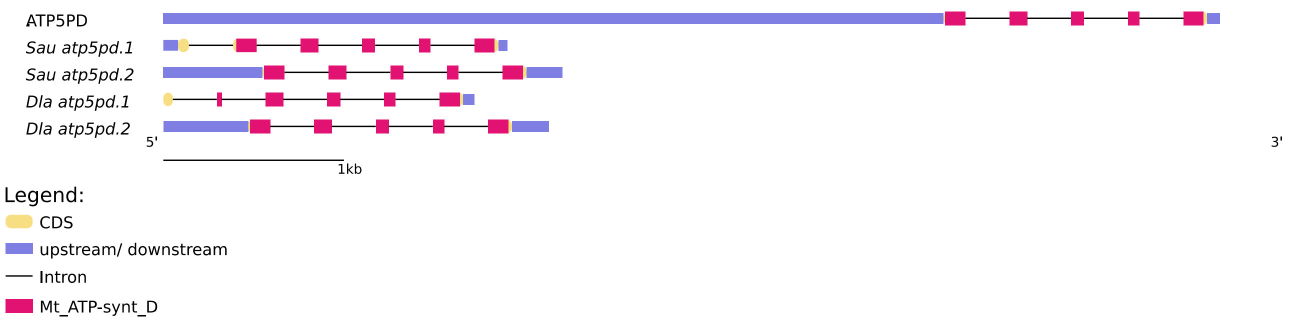

***Figure S44.*** *Gene structure and protein domains for atp5pd gene family in gilthead seabream (Sau), European seabass (Dla) and human (capital letters).*

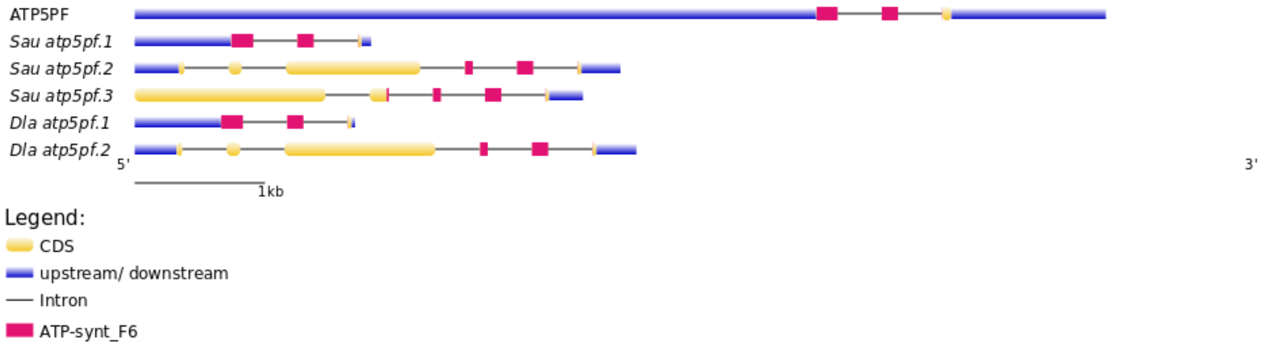

***Figure S45.*** *Gene structure and protein domains for atp5pf gene family in gilthead seabream (Sau), European seabass (Dla) and human (capital letters).*

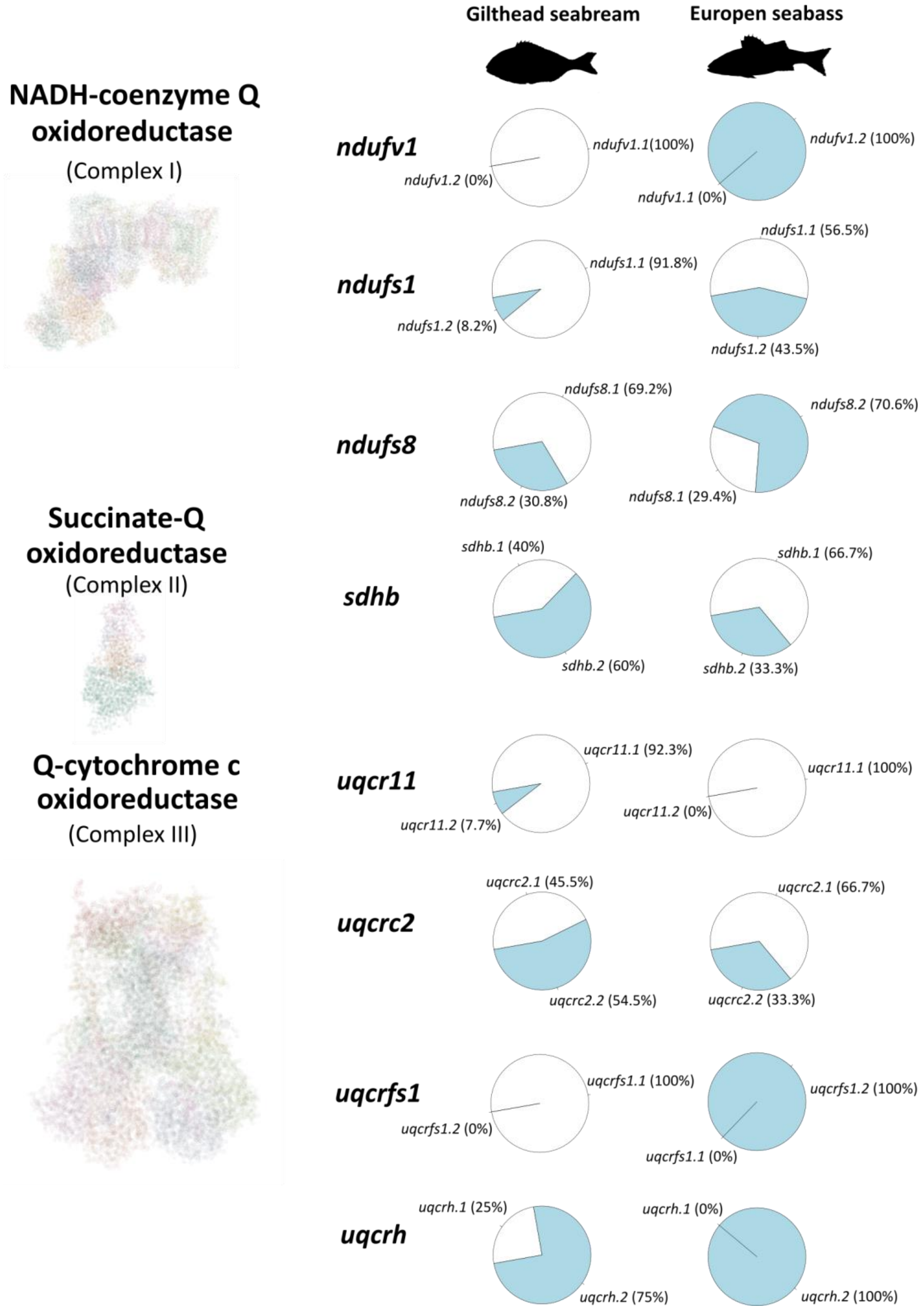

***Figure 46**. Distribution of mutation of paralogs per gene family for seabream (left) and seabass (right), for OXPHOS complexes I, II and III.*

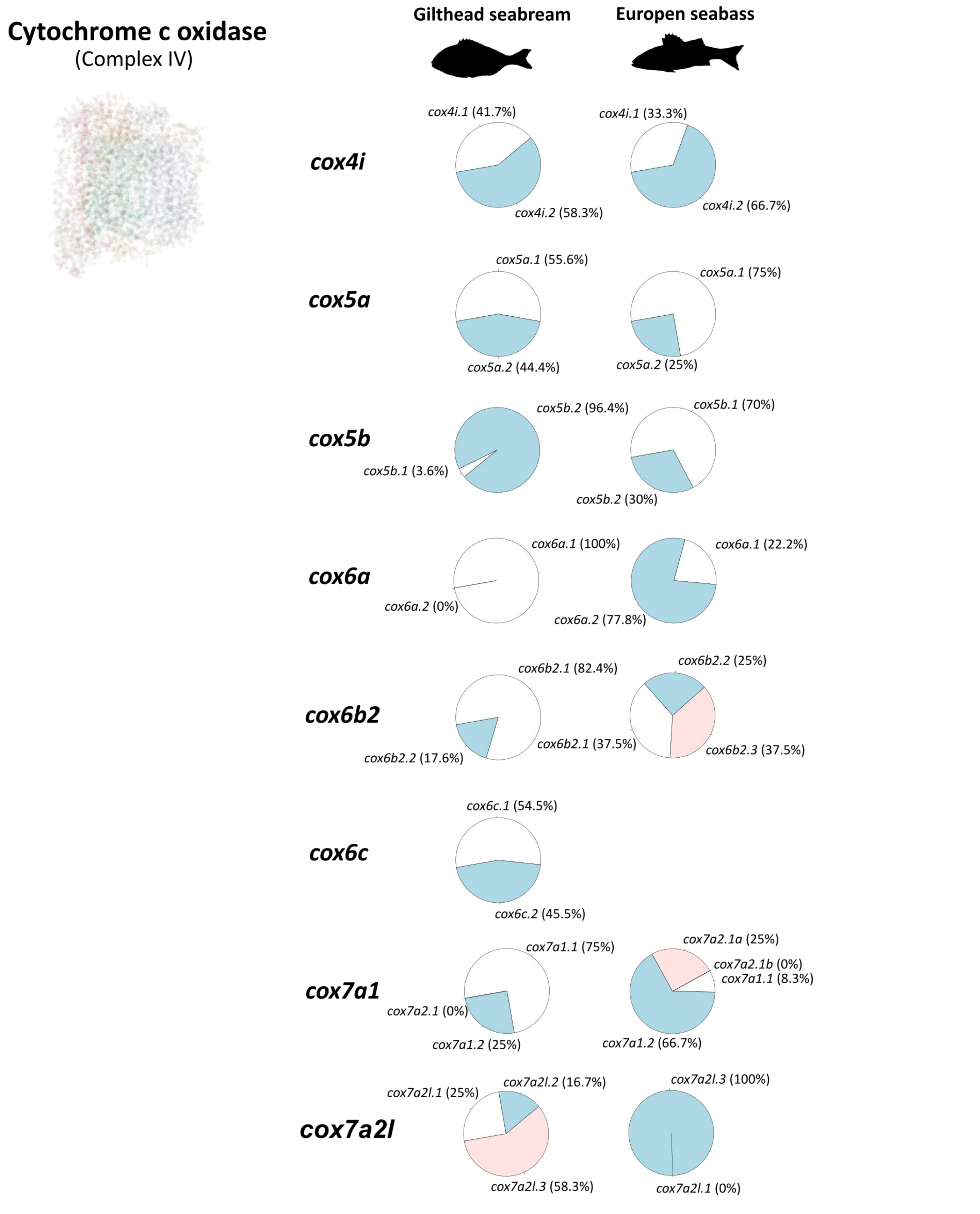

***Figure 47**. Distribution of mutation of paralogs per gene family for seabream (left) and seabass (right), for OXPHOS complex IV*

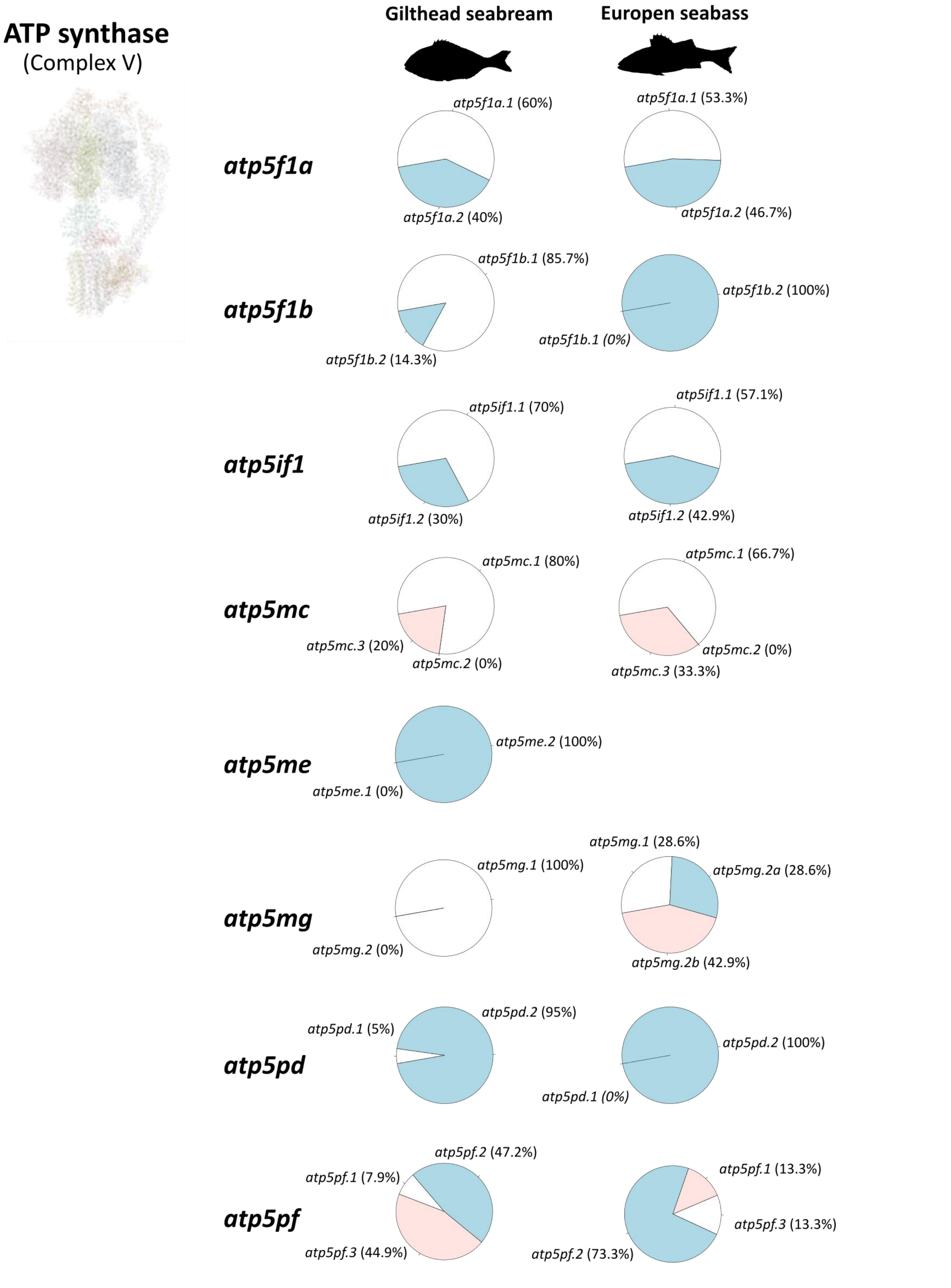

***Figure 48****. Distribution of mutation of paralogs per gene family for seabream (left) and seabass (right), for OXPHOS complex V*
